# The black honey bee genome: insights on specific structural elements and a first step towards pan-genomes

**DOI:** 10.1101/2023.12.06.570386

**Authors:** Sonia E. Eynard, Christophe Klopp, Kamila Canale-Tabet, William Marande, Céline Vandecasteele, Céline Roques, Cécile Donnadieu, Quentin Boone, Bertrand Servin, Alain Vignal

## Abstract

**Background:** The actual honey bee reference genome, HAv3.1, was produced from a commercial line sample, thought to have a largely dominant *Apis mellifera ligustica* genetic background. *Apis mellifera mellifera*, often referred to as the black bee, has a separate evolutionary history and is the original type in western and northern Europe. Growing interest in this subspecies for conservation and non-professional apicultural practices, together with the necessity of deciphering genome backgrounds in hybrids, triggered the necessity for a specific genome assembly. Moreover, having several high-quality genomes is becoming key for taking structural variations into account in pan-genome analyses.

**Results:** Pacific Bioscience technology long reads were produced from a single haploid black bee drone. Scaffolding contigs into chromosomes was done using a high-density genetic map. This allowed for a re-estimation of the honey recombination rate, over-estimated in some previous studies, due to mis-assemblies resulting in spurious inversions in the older reference genomes. The sequence continuity obtained is very high and the only limit towards continuous chromosome-wide sequences seem to be due to tandem repeat arrays usually longer than 10 kb and belonging to two main families, the 371 and 91 bp repeats, causing problems in the assembly process due to high internal sequence similarity. Our assembly was used together with the reference genome, for genotyping two structural variants by a pan-genome graph approach with Graphtyper2. Genotypes obtained were either correct or missing, when compared to an approach based on sequencing depth analysis, and genotyping rates were 89 and 76 % for the two variants respectively.

**Conclusions:** Our new assembly for the *Apis mellifera mellifera* honey bee subspecies demonstrates the utility of multiple high-quality genomes for the genotyping of structural variants, with a test case on two insertions and deletions. It will therefore be an invaluable resource for future studies, for instance including structural variants in GWAS. Having used a single haploid drone for sequencing allowed a refined analysis of very large tandem repeat arrays, raising the question of their function in the genome. High quality genome assemblies for multiple subspecies such as presented here, are crucial for emerging projects using pan-genomes.

## Background

The honey bee *Apis mellifera* was originally found in Europe, Africa and the Middle East, with the most eastern limit of its natural distribution situated in western Afghanistan until a new subspecies was discovered in Kazakhstan [1]. The evolutionary origin of *Apis mellifera* is still unclear, with a possible origin in Eastern Africa or the Middle East, followed by the colonization of Europe through different routes, leading to high genetic differentiation between geographically close populations or subspecies, namely *A. m. mellifera* (otherwise referred to as M-type) in western Europe on one side and *A. m. ligustica* from Italy or *A. m. carnica* (known as C-type) from eastern Europe on the other [2–5]. However, although *A. m. mellifera* is the original subspecies found in western Europe, it has become commonplace amongst breeders, in order to increase production or to facilitate the handling of colonies, to import other subspecies, mainly *A. m. ligustica* from Italy, *A. m. carnica* from Slovenia and *A. m. caucasica* from Georgia, either to be bred as pure lines or as hybrids generated by artificial or directed insemination [6,7]. As a consequence, these imported subspecies and hybrid lines will mate naturally to local *A. m. mellifera* populations, threatening them and prompting the establishment of conservation programmes [8]. However, although it has been replaced in the majority of large professional beekeeper’s facilities by imported honey bees, *A. m. mellifera* is still used by dedicated breeders.

The honey bee reference genome, whose first version was obtained as soon as 2006 [9], was updated twice: a first time in 2014 [10] and a second time in 2019, using long-read sequencing together with Hi-C chromatin interaction and BioNano Optical maps for a chromosome-scale assembly [11]. The sample used for this reference genome is from a commercial line (DH4), which is not precisely genetically defined, but is thought to be mainly of *A. m. ligustica* descent [9]. As a consequence, the genome of the genetically distinct *A. m. mellifera* may not be accurately represented and future pangenome approaches, that were shown in other species to expand the number of genomic regions available for analysis [12,13], would benefit from a high-quality assembly for this important subspecies.

To ensure a faithful representation of the *A. m. mellifera* subspecies genetic background, an individual from the black bee conservatory “Association Conservatoire de l’Abeille Noire Bretonne” in the island of Ouessant, France was selected for sequencing. This very small island (15.5 km^2^) is located 20 km off the coast of Brittany, the conservation population was set up starting in 1987, and further imports of other honey bees banned since 1991. Mitochondrial DNA analyses have shown a low haplotype diversity and the presence of only the M-type in this population [14]. As expected from such a small population, microsatellite analysis has shown a low diversity [15].

Until the latest update [11], the current honey bee genome sequence, Amel4.5 [10] suffered from imperfections, having numerous gaps in the assembly and possible sequence inversions. In order to construct a new *A. m. mellifera* genome assembly with improved continuity, we used the Pacific Biosciences long-read technology and produced all sequence reads from a single haploid drone to avoid assembly problems due to polymorphism. To order and orient our contigs along the chromosomes, we used published sequencing reads from drones originating from three colonies that had previously been used to map meiotic crossovers and non-crossovers in the honey bee [16], allowing also for the production of an updated genetic map and a re-estimation of the honey bee recombination rate.

Our analyses of the assembly allowed the detection of a major family of tandem repeats, running in some instances over more than 10 kb and found at the ends of most sequence contigs. Our assembly allows for the first-time to perform detailed analyses of structural rearrangements, including at the population level, between the genomes of *A. m. ligustica* and other C-type honey bees used by the majority of beekeepers and that of the M-type subspecies *A. m. mellifera* black bee.

## Methods

### Sampling, DNA extraction and PacBio long-read sequencing

Candidate drones for sequencing were sampled at the larval or pupae stage from the black bee conservatory on the island of Ouessant, Brittany, France and extractions were performed from several samples, to select the best DNA quality in terms of molecular weight and quantity. Each sample was ground using a potter (see Additional file 1: Fig. S1) and DNA extraction performed using the QIAGEN Genomic-tips 100/G kit (Cat No./ID: 10243), following the tissue protocol extraction (see supplementary methods). DNA for sequencing was obtained from a single drone OUE7B (see Additional file 1: Fig. S2). Library preparation and sequencing were performed at the GeT-PlaGe core facility, INRAE Toulouse, following the manufacturer’s instructions for “Shared protocol-20kb Template Preparation Using BluePippin Size Selection system (15kb size Cutoff)”. At each step, DNA was quantified using the Qubit dsDNA HS Assay Kit (Life Technologies). DNA purity was tested using the nanodrop (Thermofisher) and size distribution and degradation assessed using the Fragment analyzer (AATI) High Sensitivity Large Fragment 50kb Analysis Kit.

Purification steps were performed using 0.45X AMPure PB beads (PacBio). Thirty µg of DNA was purified to perform 3 libraries. Using SMRTBell template Prep Kit 1.0 (PacBio), a DNA and END damage repair step was performed on 15µg of unshared sample. Then blunt hairpin adapters were ligated to the libraries. The libraries were treated with an exonuclease cocktail to digest unligated DNA fragments. A size selection step using a 7kb (Library 1) or 9kb (libraries 2 and 3) cutoff was performed on the BluePippin Size Selection system (Sage Science) with the 0.75% agarose cassettes, Marker S1 high Pass 15-20kb. Conditioned Sequencing Primer V2 was annealed to the size-selected SMRTbells. The annealed libraries were then bound to the P6-C4 polymerase using a ratio of polymerase to SMRTbell at 10:1. Then after a magnetic bead-loading step (OCPW), SMRTbell libraries were sequenced on 36 SMRTcells on a RSII instrument from 0.05 to 0.2 nM with a 360 min movie.

### Assembly into contigs and alignment to Amel4.5 for chromosome assignments

Raw reads were assembled with Canu 1.3 [17] using standard parameters and a first polishing of the assembly was done with quiver (version SMRT_Link v4.0.0) using standard parameters. The contigs obtained after the assembly step were aligned to the Amel4.5 reference genome using LAST v956 [18].

### Alignment of Illumina sequencing reads and SNP calling for crossing over analysis

All the Illumina paired-end sequences from Liu et al. (2015) [16] were downloaded from the NCBI SRA project SRP043350 (see Additional file 2: Table S1). The reads were aligned to the assembled contigs with BWA MEM v0.7.15 [19], duplicate reads removed with Picard (v2.1.1; http://picard.sourceforge.net), and local realignment and base quality score recalibration (BQSR) performed using GATKv3.7 [19]. SNPs were called in each drone independently with GATK HaplotypeCaller and consolidated into a single set of master sites, from which all individuals were genotyped with GATK GenotypeGVCFs (see scripts in supplementary material). Any SNP with missing genotypes were filtered out. Further quality controls were applied and for each colony, SNPs falling into any of the following categories were discarded: i) non-polymorphic SNPs in the colony, ii) homozygous SNPs in the queen, iii) heterozygous SNPs in drones, iv) SNPs that appeared inconsistent with the observations in the two other colonies and v) SNPs showing inconsistent allelic versions between queen and drone genotypes.

### Phasing and detection of recombination events

For each colony and informative SNP, genotyping results were used to define genotype vectors across all drones for the colony. Identical genotype vectors following one another within a same contig define a segment with no observed crossing over in the drones of the colony and were grouped into bins. Not having access to grand-parental genotypes, genotype phase between two successive bins within a contig was determined by finding which out of the two possible inverse vectors minimised the number of recombination events. Non-crossing over gene conversion events, which can be misinterpreted as double recombination events, occurring usually on short DNA fragment often considered shorter than a few kb, [16] were removed to avoid inflating the size of the genetic map. In our study, non-crossing over gene conversion events were identified as: i) bins of length shorter than 2 kb, occurring between two identical bins, or ii) bins of length shorter than 2 kb for which the number of recombination events happening within this bin is higher than the number of recombination events needed to go from the bin before to the bin after it. Bins detected as non-crossing over gene conversions were merged with their two identical surrounding bins. Both phasing and putative non-crossing over identification were performed iteratively from one bin to the next and independently for each colony. As a consequence, a set of phased vectors minimising recombination events was obtained for each contig in each colony.

### Scaffolding contigs into chromosomes

Using the *a priori* assignment of contigs to chromosomes by alignment to Amel4.5 as a starting point, contigs were ordered and oriented iteratively in order to minimise the number of recombination events between the genotype vectors defined at their extremities. The contig scaffolding was first performed using the data for each colony separately and was thereafter confirmed using markers informative across all three colonies.

### Correction of the assembly with Illumina reads

Genomic DNA from the same individual used for the PacBio sequencing was sequenced with an Illumina NovaSeq6000 instrument, producing over 28 000 000 reads (estimated raw sequencing depth = 37 X), NCBI SRA accession SRR15173860. These were aligned on the assembled genome with BWA MEM version 0.7.12-r1039 [20] using standard parameters. Variant detection was done with freebayes version 1.1.0 [21] and filtered to retain only those with a minimum quality score of 20 and ’1/1’ genotype or ’0/1’ with no read supporting the reference allele. Finally, corrections to the genome assembly were done when alternative alleles were found in the VCF file using vcf-consensus from the vcftools package (version 0.1.12a) [22] with standard parameters.

### Comparison with Amel 4.5 and HAv3.1 assemblies

Estimation of recombination rate and positioning recombination events along the Amel4.5 and AMelMel1.1 assemblies was done following the same procedure as for the de-novo assembly. GC content and sequence coverage for the queens’ genotypes in AMelMel1.1 were measured in 0.5Mb windows and the recombination rates were estimated using a script from Petit et al. (2017) [23] over 1Mb windows. Completeness of the assemblies was estimated with BUSCO 3.0.2 [24] using OrthoDB v9.1 single-copy orthologs [25], from the Metazoa (n=978) and Hymenoptera (n=4415) BUSCO core set. Alignments of AMelMel1.1 to Amel4.5 and to HAv3.1 were done using LAST v956 [18]. Standard output psl files were produced to keep all alignments related to repeat elements, together with psl files from split alignments [18], corresponding to one-to-one alignments. Dotplot visualisation of alignments were produced with custom scripts. Inversions between the two genome assemblies were detected in the split alignment psl file. Liftovers of the HAv3.1 gtf and gff annotation to produce files with AMelMel1.1 annotation coordinates were done using CrossMap [26] and the chained alignment format output from the AMelMel1.1 to HAv3.1 LAST alignments.

### Analysis of repeat elements

Analysis of tandem repeats was done with Tandem Repeat Finder v4.09 (TRF) [27], setting the maximum period size to 2000 bp. The two major classes of repeat sizes: the 91 bp repeat and the 371 bp repeat were analysed by aligning all repeats within a class size with MAFFT v7.313 [28]. Sequences reported by TRF from different parts of the genome start at different positions of the repeated element detected and as a consequence, the multifasta alignments produced by MAFFT were processed with a custom script, to determine an identical arbitrary start point for all sequences, before performing a second alignment with MAFFT. Phylogenetic trees were calculated in Jalview v2.11.2 [29] with the average distance option. Consensus sequences from all sequences selected within the groups defined based on the phylogenetic trees were used for a BLAST search in the AMelMel1.1 assembly and hits following one another at distances shorter than the repeat period size were grouped together.

The previously described monomer consensus sequences: accession X57427.1 for *Alu*I and X89530.1 for *Ava*I were used to detect their presence in the assembly by BLAST.

### Analysis of indels in populations

Indels were detected by aligning the two genomes HAv3.1 and AMelMAl1.1 to one another with minimap2 [30], followed by variant calling with SVIM-asm [31]. Two nuclear mitochondrial DNA (NUMT) were then selected for genotyping in a set of 80 haploid males representing the three major European bee subspecies: *A. m. mellifera (n=35)*, *A. m. ligustica (n=30)* and *A. m. caucasica (n=15)* (see Additional file 2: Table S2). All 80 samples were aligned to both assemblies as described in Wragg et al. (2022) [6] and sequencing depth was estimated using SAMtools [32]. Individual genotypes in the samples sequencing data was determined for the two indels by two methods. One method consisted in using GraphTyper2 [33], that will detect breakpoints due to insertions, deletions or inversions in the pangenome graph built with SVIM-asm using the two assemblies HAv3.1 and AMelMel1.1. The other method consisted in using sequencing depths as an indication of presence or absence of Indels. For a given Indel and for each sample, the sequencing depth for the alignments on the genome in which the Indel is present was calculated and compared to the sequencing depth of the sequences flanking the Indel on both sides. Normalisation was done by calculating the ratio between sequencing depth in the indel and in the flanking sequences. Genotype presence or absence was then done by K-means clustering with K=2.

## Results

### PacBio long-read sequencing and assembly into contigs

All long-read sequence data comes from a single haploid drone selected amongst several tested, as having the highest DNA concentration and a peak of DNA fragment length at 35 kb (see Additional file 1: Fig. S2). A high proportion of reads exceeds 10 kb and a few reads are longer than 70 kb. Their size distribution is shown in Additional file 1: Fig. S3 and S4. After assembly, a total of 200 contigs (gap-free sequence tracts) was obtained. The longest contig is 11.6 Mb and the N50 contig size is 5.1 Mb (see Additional file 2: Table S3 and Additional file 1: Fig. S5). These results are a major improvement in comparison to the 46 kb N50 contig of Amel4.5 and quite similar to the N50 contig of 5.4 Mb observed in the HAv3.1 assembly [11]. Analysis with BUSCO showed that overall, AMelMel1.1 had a slightly larger gene content than both Amel 4.5 and the most recently published reference assembly AmelHAv3.1 [34] (see Additional file 2: Table S4).

### Chromosomal assignation and ordering contigs with crossing-over data

A priori chromosomal assignment of contigs was done by alignment to the Amel4.5 assembly using LAST v956 [18]. Out of the 200 contigs, 110 aligned successfully. Crossing-over data for confirming assignation and ordering contigs along chromosomes was obtained by using the reads from the sequencing of 43 drones from three colonies, initially used to estimate recombination rate in honey bee [16]. Briefly, this data set contains sequence for three queens and their drone offspring (15 to 13 depending on the colony). Three of the drones of colony 1 are sequenced in duplicate and are used for the quality control of SNP calling. Aligning these reads to our contigs allowed the detection of 2,103,924 SNPs, on 176 contigs before any quality control. Out of these, approximately 64.5% were discarded due to an absence of polymorphism within the three colonies analysed, 1% for being homozygous in the queens and 1% for being heterozygous in the drones. Furthermore, 0.2% of the SNPs were discarded for being inconsistent between the three drone replicates and finally 0.4% were discarded for having allelic inconsistencies between queen and drones of the same colony. After all the quality controls and for each of the three colonies, 687,699; 698,123 and 672,728 reliable SNPs (approximately 32% of the initial SNPs), were detected in each of the three colonies on 114, 112 and 113 contigs respectively (see Additional file 1: Fig. S6). In total 120 contigs were at least partially informative across the colonies, with 104 contigs informative in the three colonies and 16 for only one or two. A total of 114,754 polymorphic SNPs was present overall in the 104 contigs informative across all three colonies (see Additional file 1: Fig. S6). Genotype vectors for each SNP across colony drones were then defined, allowing for within-contig crossing-over detection (see Additional file 1: Fig. S7). Genotype vectors from the ends of contigs were then used to join contig ends together by finding for each contig end, the best corresponding end from another contig having either the same genotype vector or a genotype vector presenting a minimal number of crossing-overs (see Additional file 1: Fig. S7). To minimize the number of comparisons, the *a priori* chromosomal assignment by alignment to Amel4.5 (see above) was used.

One hundred and two contigs out of the 110 with chromosome assignment by sequence similarity to Amel4.5, had SNP genotype data and were thus informative for crossing-over detection. At least one crossing-over event, as evidenced by the presence of at least 2 genotype vector bins, could be detected within 86 of these contigs, thus allowing for their orientation. The remaining 16 contigs were oriented based on the alignment to Amel4.5. All these contigs were small, except one contig on chromosome 7. Despite its large size, close to 2.4 Mb, it was indeed difficult to orientate using the genetic map, as no crossing-over could be detected due to an unusually low number of SNPs and a very low local recombination rate. Moreover, its orientation could not be deduced from Amel4.5 or even from the more recent assembly HAv3.1, as both possible orientations induced large inversions when compared to these other two assemblies. Contigs assigned to chromosomes by alignment only (8 contigs) or by crossing-over data alone (16 contigs), were assigned to their chromosomes, but at an unknown (unlocalised) position. All remaining 72 contigs were considered unplaced (see Additional file 1: Fig. S6).

### Tandem repeats at contig boundaries and Orientation of a large inversion on chromosome 7

With long read data, sequence contigs are large, but still don’t cover the entire length of chromosomes, with the exception of chromosome 16. When analysing the contig ends, we found that almost all were composed of tandem repeats arrays usually longer than the read lengths, thus preventing assembly. To orientate the large contig on chromosome 7, positioned as 5^th^ in order along the chromosome by the CO data, we took advantage of the fact that the repeat elements detected by TRF and present at both extremities of the contig have different period sizes (258 and 1296 bp) and consensus sequences. These were compared to the proximal repeats of the 4th and the 6th contigs of chromosome 7. Interestingly, a tandem repeat element of 258 bp was detected at the end of the 4th contig, and of 1296 bp at the end of the 6th contig, period sizes identical to the extremities of the 5^th^ contig, suggesting the correct orientation of the 5^th^ contig. Correspondence between these contig ends was further examined by pairwise alignment of the repeat sequences with NCBI BLAST. The Identity was 100 % between the sequences of identical period sizes, whereas no significant similarity could be found between the others (see Fig. 1 and Additional file 2: Table S9), thus confirming the orientation of the contig. Dotplots comparing AMelMel1.1 and HAv3.1 are shown in Additional file 3 and suggest a very small number of discrepancies, the major one residing on chromosome 7.

**Figure 1.**
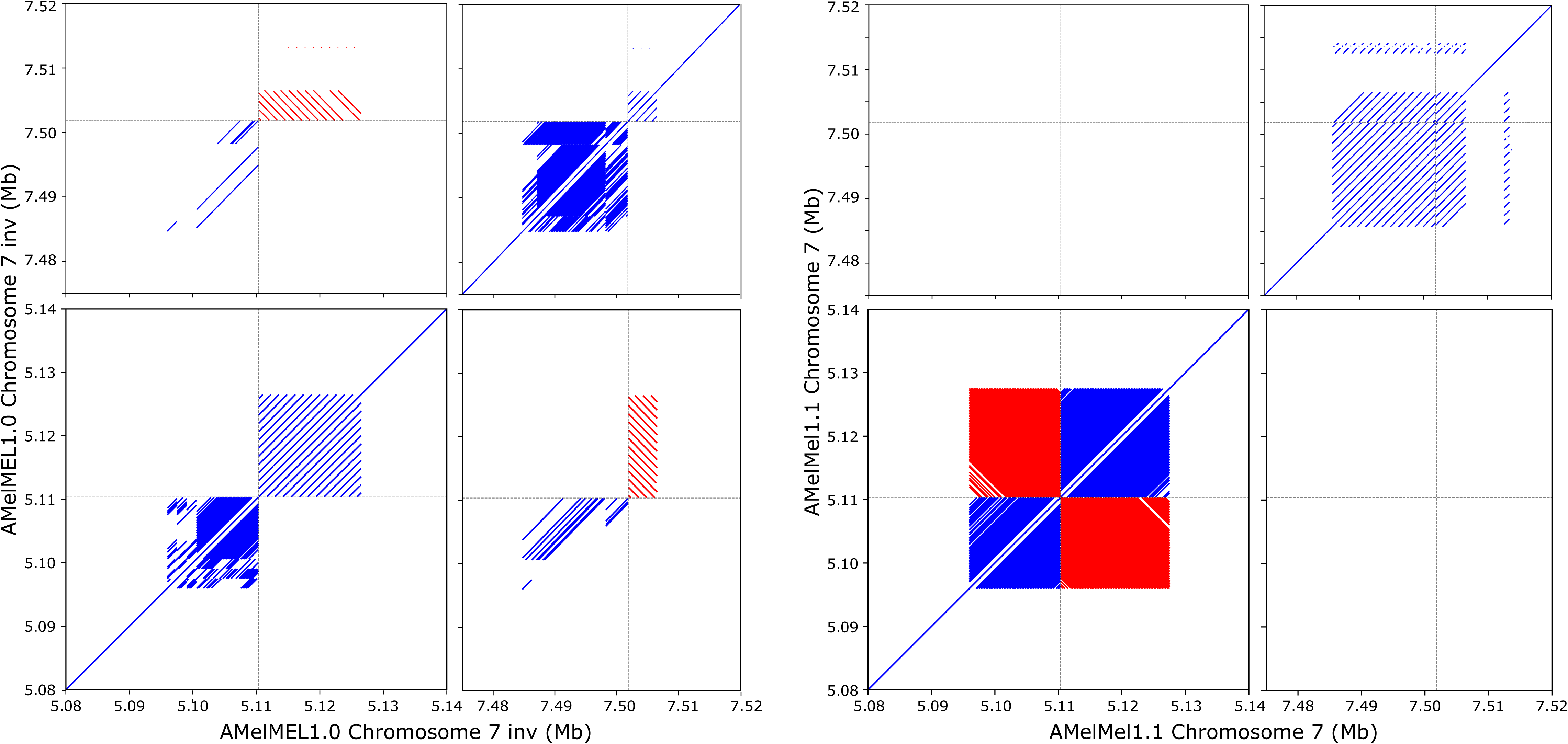
Orientation of the AMelMel1.1 contig presenting an inversion on chromosome 7 when compared to HAv3.1. The repeats present at the boundary between the contigs were used to orient the AMelMel1.1 contig on chromosome 7. Assemblies with one or the other orientation of the contig were self-aligned with LAST. Left: orientation from AMelMel1.0 and right, orientation from AMelMel1.1. For each pair of alignments, only the junction between contigs are shown: the two ends of the contig to orient, the end of the previous and the start of next contigs. Results clearly show the orientation in AMelMel1.1 is the correct one.

### Telomeric and centromeric consensus sequences

The presence of telomeres is an indication of the completeness of the assembly. These were analysed by searching for the accepted TTAGG consensus sequence for Hymenoptera [35] in TRF analysis output, estimating their distance to the ends of chromosomes and comparing the results to that of other 2-7 bp repeats, including non-TTAGG 5 bp repeats. Results (Fig. 2) show that TTAGG are repeated with at least 842 copies when present at the extremities of chromosomes, whereas other interstitial TTAGG repeats have only 117 repeats or less (mean = 21.3, median = 16.7), a size distribution close to that of other pentanucleotide repeats (mean = 24.2, median = 14.4). See also Additional file 1: Fig. S8 and Additional file 2: Table S6, S7 and S8 for data on other STR motifs. In the AMelMel1.1 assembly, no TTAGG repeats were found on chromosomes 3, 7, 12 and 15 and were found only at the beginning of chromosome 1, whereas in the HAv3.1 assembly, they could be found at both extremities of this chromosome, but were absent from chromosomes 5 and 11 [11]. An AATAT repeat was found at the beginning of chromosome 15 in our assembly.

**Figure 2.**
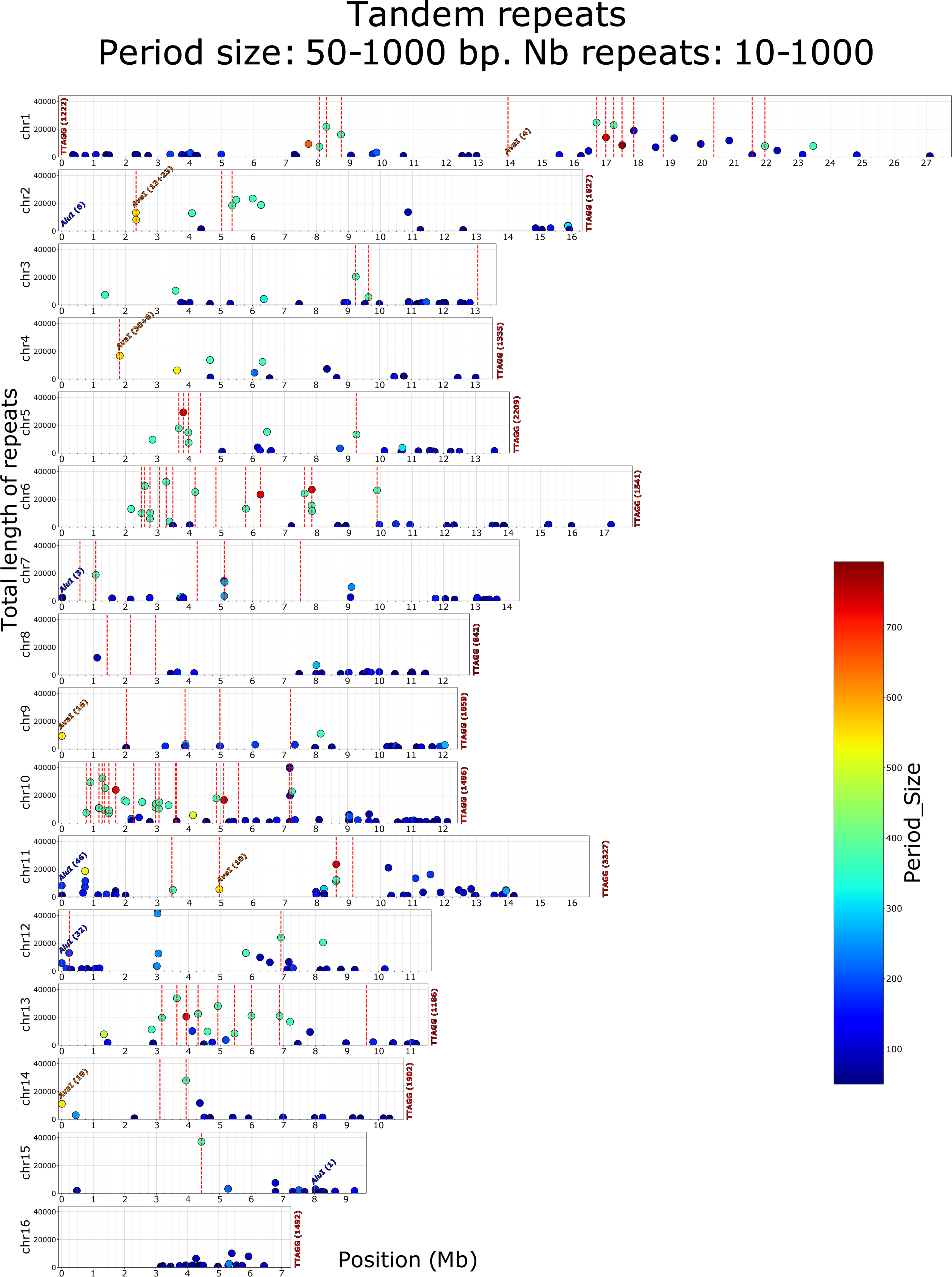
Comparison of Amel4.5 and AMelMel1.1 assemblies for chromosome 3. Abscissa: AMelMel, ordinate: Amel4.5. AMelMel contig borders are represented with vertical dotted lines. Additionally, for both Amel4.5 and AMelMel, the position and number of recombination events detected along the chromosome are represented for each interval flanked by informative markers in the meiosis analyzed. Average SNP density and recombination rate are given for 1Mb windows. Regions indicated in red on the Amel4.5 assembly represent recombination ‘hotspots’ regions where number of recombination events between two informative SNPs is higher than five. See supplementary data for the other chromosomes.

The *Alu*I and *Ava*I repetitive sequences, previously described as being respectively telomeric and centromeric were localised on the AMelMel1.1 assembly by BLAST search and the number of copies per locus detected was counted (Fig 2). The *Alu*I repeat was found at the start of chromosomes 2 (6 repeats), 7 (3 repeats), 11 (46 repeats) and 12 (32 repeats). In addition, a single *Alu*I element was found around position 8 Mb on chromosome 15, at more than 1.5 Mb from the distal end. Curiously, the *Alu*I repeats found on chromosomes 2 and 11 were at the opposite end from the TTAGG sequences we detected (Fig. 2). The *Ava*I repeat was found as arrays at single loci on chromosomes 1, 2, 4, 9, 11 and 14. Only 4 copies in the array were found on chromosome 1, the other arrays having between 10 and more than 30 copies. The *Ava*I repeats are at the start of chromosomes 9 and 14, at the opposite end from the TTAGG repeats. On the other four chromosomes, they are at least at 1.8 Mb from a chromosome end (Fig. 2).

### Recombination pattern

Having used crossing-over detection and a genetic map for contig scaffolding, we could estimate the total genetic map for AMelMel, which is approximately 50 Morgans long, giving an average recombination rate in the genome of 23 cM/Mb, close to the first estimations based on the microsatellite genetic map and to the most recent ones based on SNPs (Table 1). However, although we used the same sequencing dataset as in Liu et al. (2015) [16], we found a drastic reduction in recombination rate between our genetic map and the one they initially published, which is 37 cM/Mb (Table 1). A great difference is that the latter is based on alignments of the sequence reads on Amel4.5. When aligning our assembly with Amel4.5, we find an agreement on the chromosomal assignment of the contigs, but reveal many discrepancies in the orientation of large chromosome segments. At most breakpoint positions between the two assemblies, recombination hotspots are detected on Amel4.5 (Fig. 3 and Additional file 4), suggesting these assembly errors were responsible for the overall higher recombination rate observed in Liu et al. (2015) [16]. This reduction from 37 cM/Mb to 23 cM/Mb is explained by these artefactual recombination hotspots detected on Amel4.5 at the breakpoint positions where the two assemblies disagree, that are absent in AMelMel1.1 (i.e. for chromosome 3 shown in Fig. 3 and Additional file 4 for all the chromosomes).

**Figure 3.**
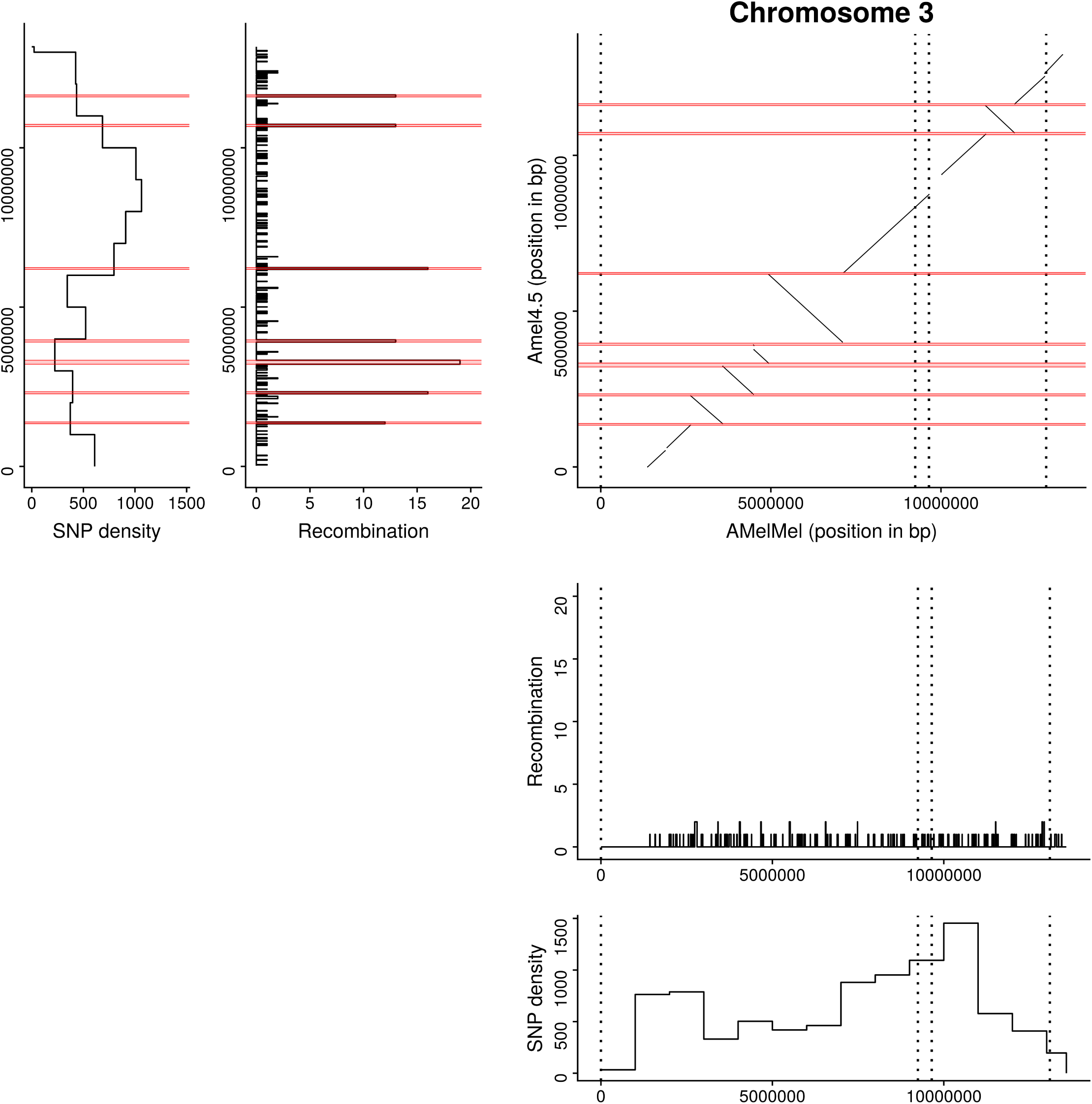
Tandem repeats of period size 90-371 bp detected in the AMelMel1.1 assembly. The colour scale represents the period size of the repeat elements and the Y axis the total length of the repeat array. Vertical dotted lines represent the contig boundaries in the AMelMel1.1 assembly. The position of *Alu*I and *Ava*1 repeats are indicated with the number of repeats in parentheses. The figure shows clearly, that most contigs are separated by tandem repeats of period size close to 371 bp, of length in the order of 10 kb or more. See also Supplementary file 1: Fig S11 for repeats of longer period size. (1000-2000 bp). Although not represented on the graph (period size = 5), TTAGG telomere are indicated with the number of repeats in parentheses, when present at a chromosome end.

**Table 1:**
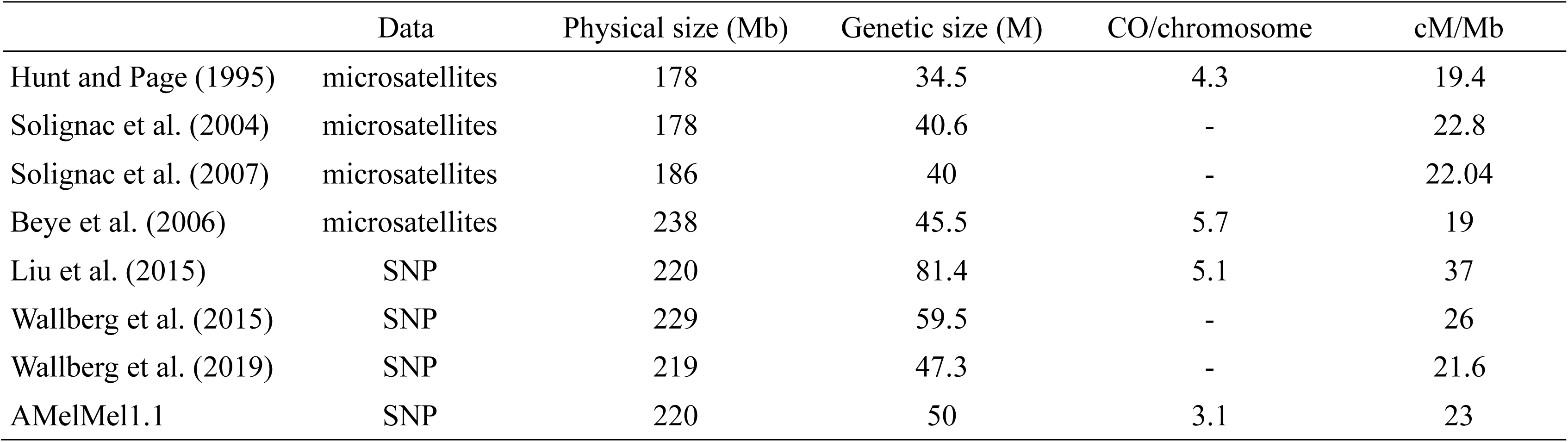
Literature comparison of *Apis mellifera* genetic maps.

|  | Data | Physical size (Mb) | Genetic size (M) | CO/chromosome | cM/Mb |
| --- | --- | --- | --- | --- | --- |
| Hunt and Page (1995) | microsatellites | 178 | 34.5 | 4.3 | 19.4 |
| Solignac et al. (2004) | microsatellites | 178 | 40.6 | - | 22.8 |
| Solignac et al. (2007) | microsatellites | 186 | 40 | - | 22.04 |
| Beye et al. (2006) | microsatellites | 238 | 45.5 | 5.7 | 19 |
| Liu et al. (2015) | SNP | 220 | 81.4 | 5.1 | 37 |
| Wallberg et al. (2015) | SNP | 229 | 59.5 | - | 26 |
| Wallberg et al. (2019) | SNP | 219 | 47.3 | - | 21.6 |
| AMelMel1.1 | SNP | 220 | 50 | 3.1 | 23 |

### High conservation of tandem repeat sequences across chromosomes

We used TRF to further localise and analyse the repeat arrays in the whole honey bee genome. Interestingly, two major period size classes for tandem repeats could be found: one in the size range of 91-93 bp, with a maximum number of 231 repeats, hereafter called the 91 bp repeat and the second in the size range of 367-371 bp, with a maximum number of 100 repeats, called the 371 bp repeat (see Additional file 1: Fig. S9). The 91 repeats are found on all chromosomes, whereas the 371 bp repeats are on all chromosomes except chromosome 16 (see Additional file 1: Fig. S10). Interestingly, very long repeats whose length is within the range of the sequence reads, were often found at the junction between two sequence contigs, confirming they could be responsible for the impossibility to sequence and to assemble these regions properly (see Fig. 2, Additional file 1: Fig11).

We investigated further the nature of the 91 and 371 repeats by analysing their potential homogeneity in terms of sequence content. Summary statistics for the two classes show very different distributions in terms of repeat copy numbers within tandem arrays (see Additional file 1: Fig. S12 and Additional file 2: Table S9). There is a total of 345 arrays of the 91 bp repeat in the genome and 131 arrays of the 371 bp repeats. However, these numbers drop to 43 and 74 respectively when only considering tandem arrays of more than 10 repeats, suggesting that most of the 91 bp repeats have less than 10 elements (see Additional file 1: Fig. S12). To investigate sequence homogeneity within each of the two repeat classes, we selected the repeat sequence defined by TRF for repeats having strictly more than ten copies in tandem within an array. For the 91 bp repeat, we selected for 91 ≤ period size ≤ 93 and for the 371 bp repeat 367 ≤ period size 371, as suggested by the graph shown in Additional file 1: Fig. S9. Then, for each repeat class, we performed a multi-sequence alignment with MAFFT, and produced an average distance tree with Jalview. Results show that out of the 74 sequences of the 371 bp repeat class, 72 were clearly grouped together, having high similarity (Fig. 4), whereas the 43 sequences of the 91 bp repeat class showed lower similarity. We therefore decided to subdivide the 91 bp repeat class into three groups of 20, 10 and 3 sequences, based on the average distance tree (Fig. 4). The remaining ten 91 bp repeat class sequences were singletons. A consensus sequence was made for each of the four group of sequences, and was used for a BLAST search in the AMelMel1.1 assembly. The homogeneity of the 371 bp consensus sequence was confirmed by the detection of a very high number of hits of high similarity covering the overall length of the queries (see Additional file 1: Fig. S13). On the contrary, for the three different consensus sequences used separately for the 91 bp repeat, alignment length and sequence similarities were lower, confirming that it to correspond more to a class size, rather than a specific repeat family based also on sequence composition (see Additional file 1: Fig. S13).

**Figure 4.**
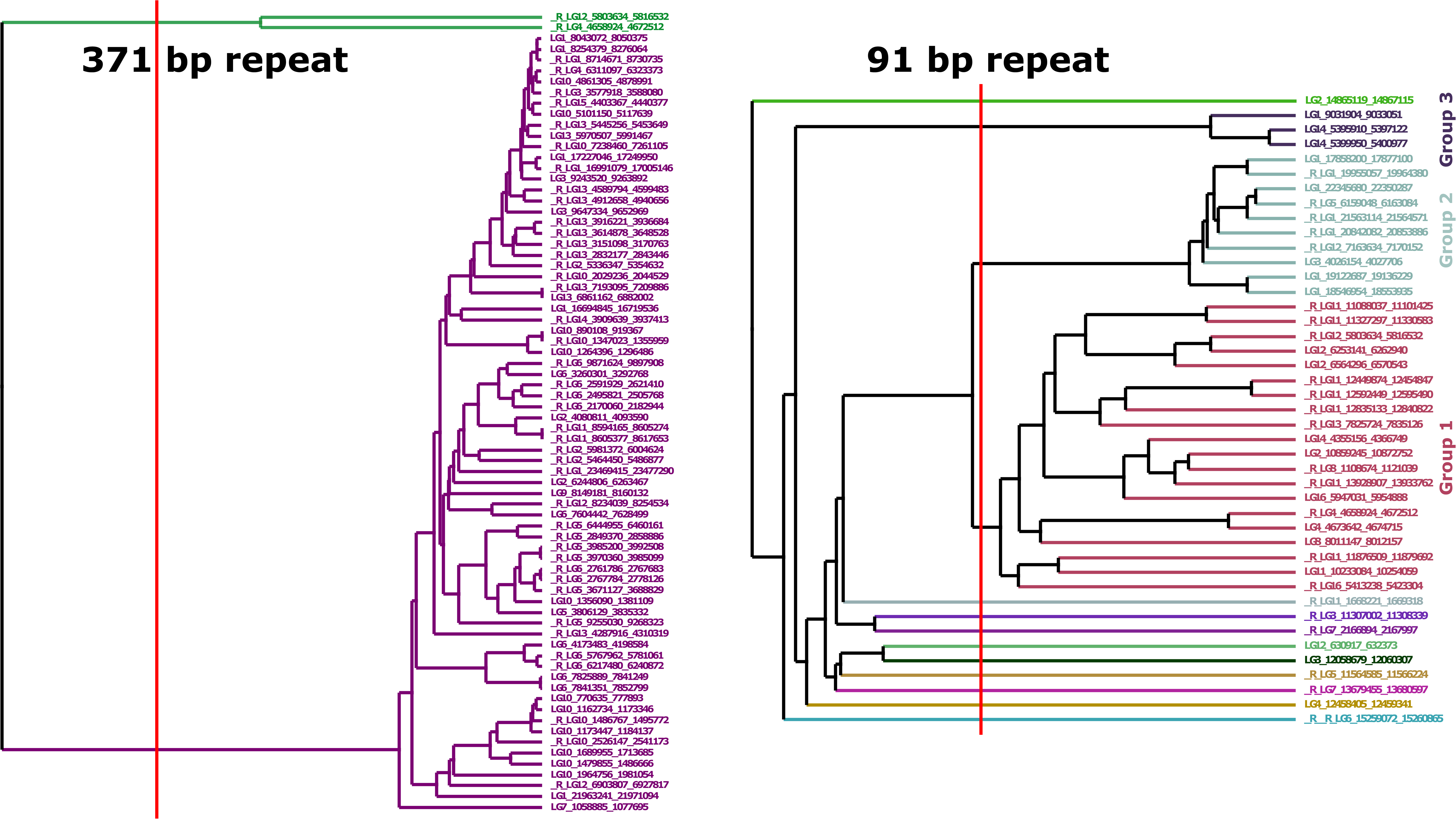
Phylogenetic trees for the tandem repeats of period size 91-93 and 367-371 bp. Only tandem repeats with ten or more elements, such as detected by Tandem Repeat Finder, were considered. Left: phylogenetic tree for the 74 sequences with a period size of 367-371 bp; right: phylogenetic tree for the 43 sequences with a period size of 91-93 bp. The vertical red lines indicate the cut-off that was used to define the groups of sequence based on similarity.

We then searched for the possible existence of the 371 and 91 bp repeats in other organisms. BLAST searches with each of the 371 bp repeat consensus sequences did not allow to find any significant hit in the NCBI nucleic collection database. When searching with each of the 91 bp consensus repeats, four hits were found: three consensus sequences from repeat arrays from chromosome 11 and one consensus sequences from a repeat array from chromosome 12 showed sequence similarity to fragments of predicted lncRNAs *LOC116185390*, *LOC105734921*, *LOC116415009*, *LOC116185696*, from unknown scaffolds of the genome assemblies of *Apis dorsata* and *Apis florea*. However, these lncRNAs are composed of two exons and span close to 1.5 kb in the genomes of *Apis dorsata* and *Apis florea*, suggesting that the 91 bp sequences correspond to only a portion (one out of two exons) of these lncRNAs. To investigate further, we performed BLAST searches with each of the consensus sequences directly on the refseq_genomes databases of *Apis cerana*, *Apis dorsata* and *Apis florea.* A very high number of hits were found, suggesting the 371 bp and 91 bp repeats were also present in these three genomes, with an apparent slightly higher percent identity for the 91 bp repeat (see Additional file 1: Fig. S14).

### Difference in the number of repeats of 5S ribosomal RNA genes

Genes that are repeated in tandem can often vary in numbers between individuals through unequal crossing-over [36]. They are therefore good candidates to study functional variation related to large rearrangements. A typical example of such genes is the 5S ribosomal RNA genes whose copy number can vary greatly in the genome [37–39]. Alignment of a region from the AMelMel1.1 and HAv3.1 assemblies in a region on chromosome 3 containing 5S ribosomal RNA genes, show a variation in the number of these genes between the two genomes (Fig. 5.). The period size of one of the repeat arrays of 5S genes is 357 bp, while that of the second is 373 bp. However, inclusion of this sequence in the multiple sequence analysis of the 371 bp repeat shows that these two sequences are different (see Additional file 1: Fig 15)

**Figure 5.**
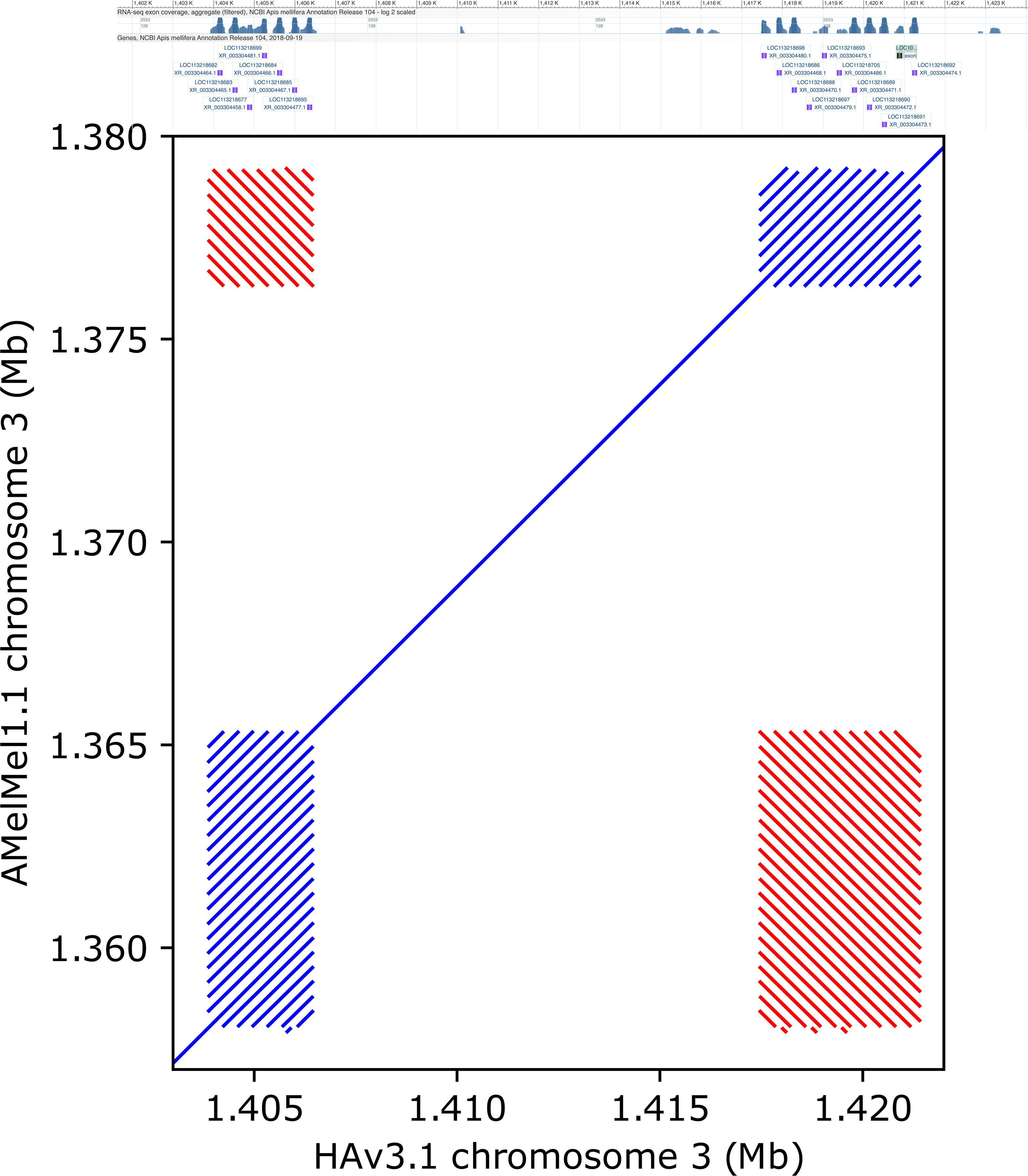
Differences in copy numbers for 5S RNA ribosomal genes. Two of the loci containing 5S RNA genes, present at 15 kb distance on chromosome 3 are shown. Top: screenshot of the NCBI genome viewer for the region showing the annotation for the 5S RNA genes. Bottom: dotplot alignment of HAv3.1 (x-axis) and AMelMel1.1 (y-axis) in the region. The first group of genes in the bottom left contains seven genes in HAv3.1 and twenty in AMelMel1.1 on the forward strand. The second in the top right contains eleven genes in HAv3.1 and eight in AMelMel1.1 on the reverse strand. The red lines off diagonal show the sequence similarity between the two groups of genes and indicate the two gene clusters are in reverse orientation.

### Inversions between AMelMel1.1 and HAv3.1

One-to-one split alignments produced by aligning AMelMel1.1 on HAv3.1 with LAST were used to detect inversions larger than 1000 bp between both genomes. The largest inversion detected is on chromosome 7 and is larger than 1.6 Mb (see Additional file 3 and Additional file 5). It should be noted, that a similar rearrangement on chromosome 7 was previously detected when comparing a genome assembly of an *A. m. ligustica* samples with the HAv3.1 reference [40]. Although close to one hundred other inversions could be detected, their visual inspection on dotplot graphs show that 53 are within complex repeat patterns present at the junction between contigs, 32 within other complex repeat elements and only 12, are in the middle of the high-quality sequence contigs in both assemblies, thus representing well supported inversions. Apart the large inversion on chromosome 7, the smallest is 1055 bp long and the largest 25608 bp long (see Additional file 2: Table S10 and Additional file 5). Interestingly, some inversions will concern genes and can involve repeat elements differentially found in both assemblies. In the example shown in Fig. 6, a local inverted duplicated region seen in the HAv3.1 assembly, is absent in AMelMel1.1. This chromosomal segment contains a portion of the gene model *LOC113218640*, which has no direct annotation in the HAv3.1 assembly, but is described as coding for a *bric-a-brac 1-like* protein. *Bric-a-brac* was shown to be involved in body pigmentation in drosophila [41]. Another interesting inversion is 11 kb long on chromosome 3, in an intron of *Rhomboid*, a gene involved in the formation of wing veins in Drosophila [42]. A more complex rearrangement involves a gene labelled as a probable nuclear hormone receptor *HR38*, involved in synchronizing the reproductive activity in *Agrotis ipsilon* [43] and in the larval-pupal transition in *Leptinotarsa decemlineata* [44]. Other genes involved in the inversions described are reported in Additional file 2: Table S10.

**Figure 6.**
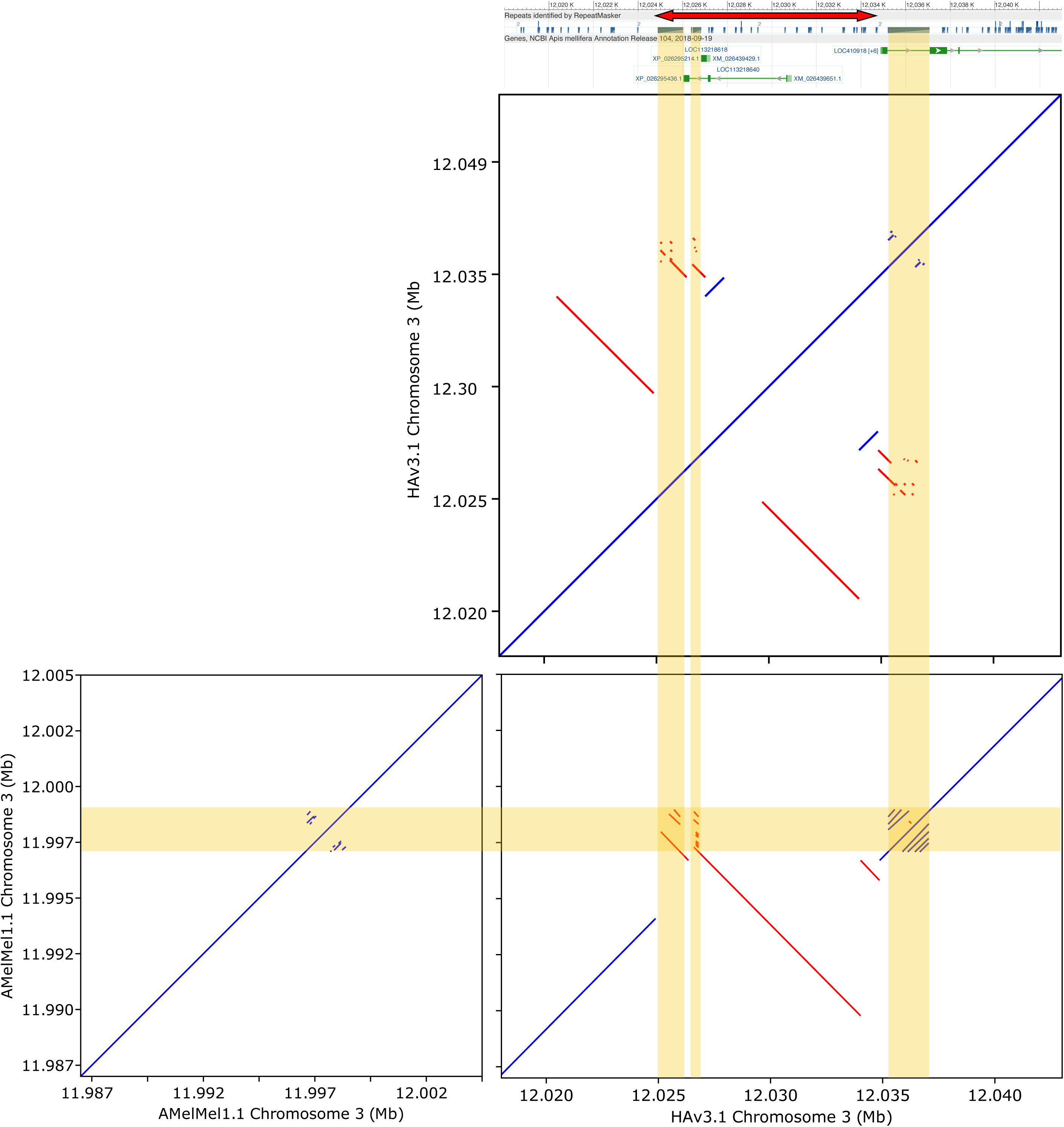
A 10 kb inverted duplication on chromosome 3 between HAv3.1 and AMelMel1.1. Bottom right: a dot plot representation of the alignment with LAST of AMelMel1.1 to HAv3.1 show a 10 kb inversion on chromosome 3. Self-alignments of AMelMel1.1 (left) and HAv3.1 (top) show that the latter has an inverted repeated sequence in the region. The vertical yellow lines show the position of repeats that were previously detected and shown in the NCBI annotation (grey boxes, bottom) and are also found in our LAST alignments. NCBI annotation of genes are in green.

### Using both assemblies for the analysis of two medium-size InDels in honey bee subspecies

To demonstrate the utility of using two reference genomes for analysing structural variants, we studied two indels corresponding to nuclear mitochondrial DNA (NUMT), that were detected by using minimap2 [30] and SVIM-asm [31]. The first one, NUMT_Chr2, is 745 bp long, has 92.7 % identity over 99 % of its length to HAv3.1 mitochondrial DNA, is present in the AMelMel1.1 assembly on chromosome 2 at positions 12,212,275 – 12,213,020 and absent from the HAv3.1 assembly. The second one, NUMT_Chr10, is 576 bp long, has 92.5 % identity over 94 % of its length to HAv3.1 mitochondrial DNA, is present in the HAv3.1 assembly on chromosome 10 at positions 670,675 – 671,251 and absent in the AMelMel1.1 assembly. The presence and absence of these two NUMTs were tested in three honey bee subspecies: *A. m. mellifera* (n=35), *A. m. ligustica* (n=30) and *A. m. caucasia* (n=15), for which Illumina sequencing data was aligned to both reference genomes. Inspection of mean sequencing depth over all 80 samples in the regions of NUMT_Chr2 and NUMT_Chr10 indicates a decrease of the mean depth and an increase of its variance (see Additional file 1: Fig. 16), suggesting the existence of a presence / absence polymorphism. When inspecting the sequencing depth per population, the *A. m. mellifera* samples show a constant value over NUMT_Chr2 and have a depth close to zero over NUMT_Chr10, whereas the *A. m. ligustica* show an inverse tendency (Fig. 7). The *A. m. caucasia* samples seem not to have NUMT_Chr2 in their genomes, whereas a few may have NUMT_Chr10, as although there is a drop of mean sequencing depth on HAv3.1 in the corresponding region, there is still some low coverage (Fig. 7). To genotype our samples individually, we used two methods. The first was to estimate individual sequencing depth in the chromosomal region delimiting the NUMTs, by using AMelMel1.1 as reference genome for NUMT_Chr2 and HAv3.1 for NUMT_Chr10 (see methods). All 80 samples could thereafter be called unambiguously assigned to one of two groups (presence or absence) by K-means clustering (see Additional file 1: Fig. 17). The second method tested was to use GraphTyper2 [33], allowing the genotyping of structural variation using pangenome graphs. Our GraphTyper2 results, showed that the calling of samples was incomplete, with a high proportion of no-calls, and that the fact of using individual bam files of alignments to one or to the other reference genome can greatly influence the call rate (see Additional file 2: Table S11). Indeed, for the detection of variants with minimap2 and SVIM-asm, a reference genome must be specified and bam files of alignments to this specific reference genome must be used to perform the individual genotyping. So, we first used HAv3.1 as reference and the results were a genotyping call rate of 78.7 % for NUMT_Chr2 and null for NUMT_Chr10, as the line describing its potential genotypes didn’t appear in the output file from GraphTyper2 at all. To check if the reference genome could influence the results, we also performed the analysis by using AMelMel1.1 as reference and this time the call rate was 85.0 % for NUMT_Chr2, and 76.2 % for NUMT_Chr10.

**Figure 7.**
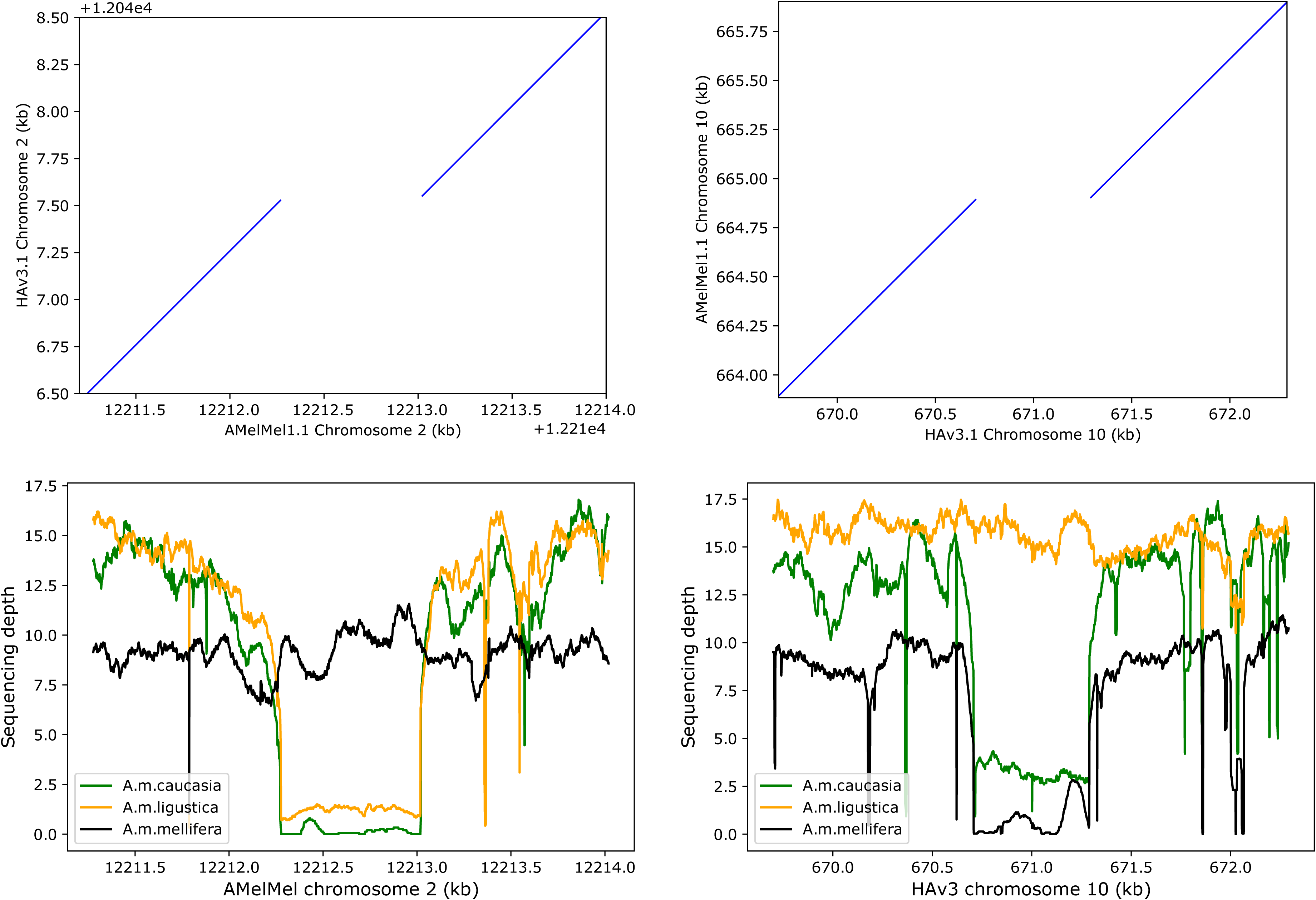
Insertions and deletions in *Apis mellifera* subspecies. Analysis of NUMT insertions detected in only one assembly. Top: dotplot representation of LAST alignments between the two assemblies show a 745 bp variant present in AMelMel1.1 on chromosome 2 and absent in HAv3.1 (left) and a 576 bp variant present in HAv3.1 chromosome 10 and absent in AMelMel1.1 (right). For each variant, sequencing depths were evaluated on the reference in which it is present. Middle: mean sequencing depth over 80 samples (red) shows a drop coinciding with the position of the variants, suggesting that a significant proportion of samples may lack the corresponding segment and standard deviation (blue) increases in the same region, confirming the heterogeneity of the samples for the presence or absence of the variant. Bottom: mean sequencing depth per subspecies, with *A. m. caucasia* (15 samples) in green, *A. m. ligustica* (30 samples) in yellow and *A. m. mellifera* (35 samples) in black. Results suggest most of the *A. m. mellifera* samples contain the insertion present in the AMelMel1.1 assembly on chromosome 2, as the sequencing depth remains constant throughout the region, and not the one present in the HAv3.1 assembly on chromosome 10, as indicated by a sequencing depth close to zero. Inversely, most of the *A. m. ligustica* samples contain the insertion present in the HAv3.1 assembly on chromosome 10 and not the one in the AMelMel1.1 assembly on chromosome 2. Most *A. m. caucasia* samples lack the insertion present in the AMelMel1.1 assembly and a few seem to have the insertion present in the HAv3.1 assembly.

When genotype calls were successfully obtained in both analyses, results were identical and were also concordant with the analysis based on sequencing depth, showing that when genoyping was possible with Graphtyper2, the results were consistent. Two samples were called as heterozygotes for NUMT_Chr2, when using AMelMel1.1 as reference and were counted as “no calls”, given our samples were haploid. Low sequencing depth could have been a possible explanation for the absence of genotyping results with GraphTyper2 in some of the samples, but this does not seem to be the case, as all samples that failed genotyping had at least 8X average sequencing depth in the sequence flanking the NUMTs analysed, whereas successful genotyping could be obtained for samples having as little as 2X sequencing depth (see Additional file: Fig. 18). Substantially, the individual genotyping results confirm the overall impression that the presence or absence of the NUMT insertions are specific to the subspecies analysed, with most, if not all samples having identical within-population genotypes, except for NUMT_Chr10 in *A. m. caucasia*, for which four out of eleven samples have a different allele. Interestingly, NUMT_Chr2 is present in all *A. m. mellifera* and only two *A. m. ligustica* samples, and absent from all other samples, whereas NUMT_Chr10 is absent from *A. m. mellifera* samples and present in all but one *A. m. ligustica* samples and four *A. m. caucasia* samples (Fig. 7, Fig. 8).

**Figure 8:**
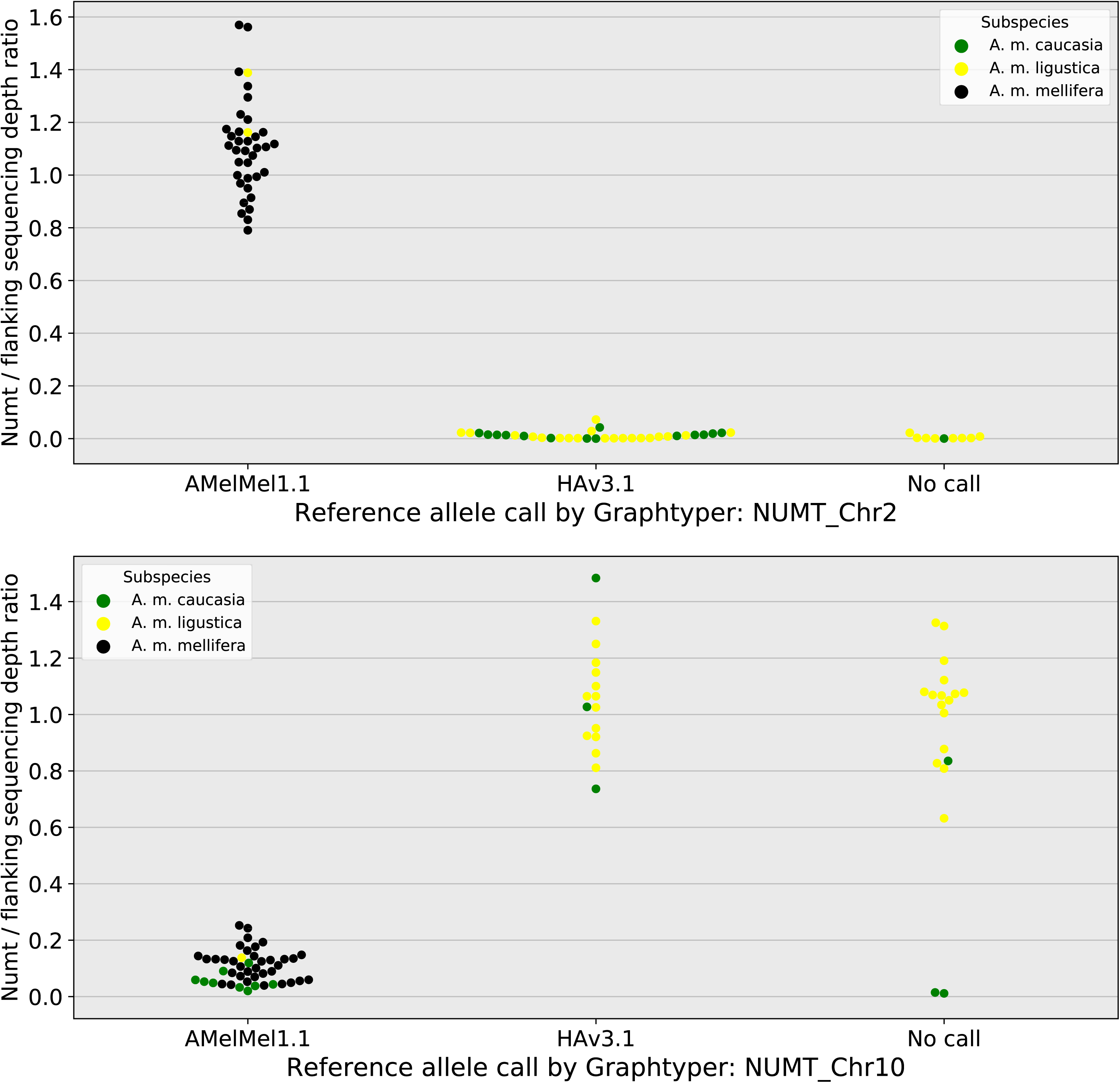
Comparing the indel variant calling between sequencing depth analysis and Graphtyper2. The Presence or absence of the NUMTs in the samples was evaluated by the pangenome graph approach with Graphtyper2 (x-axis) and by estimating the sequencing depth at the position of the NUMTs on the genome in which it is present (y-axis). Sequencing depths were normalised by calculating the ratio between sequencing depth at the position of the NUMT sequence and that of the flanking sequence. Nine out of 80 samples (11 %) could not be called for NUMT_Chr2 and 19 (24 %) for NUMT_Chr10. When alleles could be called by Graphtyper2, results agreed with the data based on sequencing depth.

## Discussion

### AMelMel assembly quality and comparison to other honey bee assemblies

Although five chromosome level genome assemblies for *Apis mellifera* are available [45] ours has the originality of representing *Apis mellifera mellifera*. Indeed, this subspecies is genetically distinct from *Apis mellifera ligustica*, *Apis mellifera carnica* and *Apis mellifera caucasi*a [46] represented by the four other assemblies. Another originality of our study, is that the contigs we obtained were scaffolded into chromosomes using a genetic (recombination) map rather than the now more common HiC chromatin conformation and Bionano optical maps methods. Compared to the current HAv3.1 reference genome [11], our assembly is slightly longer (227 Mb versus 225 Mb), is built from a lower number of contigs (200 versus 228) with very similar N50 contig values (5.1 Mb versus 5.4 Mb). However, the overall final coverage was slightly smaller in our study (137X Pac Bio and Illumina reads versus 192X in HAv3.1). BUSCO statistics are also very similar due to the fact that contig building was based in both cases on PacBio reads with some correction using Illumina reads. Assembly of contigs into chromosomes using the recombination data failed to accurately order and orient in only one instance for a large contig on chromosome seven. Despite this limitation, the orientation of this contig was possible thanks to a careful analysis of tandem repeat elements at its boundaries. Sequencing data for both HAv3.1 and our assembly, AmelMel1.1, are from a single haploid drone, which is a tremendous advantage for the resolution of regions largely composed of repeat elements. This was recently demonstrated in the human Telomere-to-Telomere project, for which a complete hydatidiform mole haploid cell line was used, helping to solve complex structures such as centromeres [47]. Our results show however, that although the sequencing of repeat elements and especially of challenging tandem repeats seems resolved by the use of a single haploid sample and long reads, there are cases in which the total length of monotonous repeats is larger than the reads lengths, preventing local assembly. As a result, for almost all contig boundaries investigated, long stretches of tandem repeats were found (Fig. 2). Interestingly, chromosome 16, which was obtained as a single contig, has no stretch of tandem repeats exceeding 10 kb.

### Genetic maps and recombination rate in the honey bee

Having used genetic recombination data to scafold our contigs, we could build a new recombination maps and give an estimation of 23 cM/Mbp for the overall recombination rate in the honey bee [16], which is of the same magnitude as the latest values from [34] and also congruent with prior values [48–50]. It is interesting to note, that the public sequencing dataset we used, representing 43 drone genome offspring of three queens, gave a much higher estimate of 37 cM/Mb when previously used for generating genotyping data by alignment on the Amel4.5 reference genome [16]. On closer inspection, this higher overall recombination rate in Liu et al. (2015) [16], is due to very specific false recombination hotspots that appear at contig junctions in Amel4.5, when at least one of them is inverted as compared to AMelMel1.1 (Fig. 3 and Additional file 4). This illustrates the importance of the quality of the reference genome for such studies. Errors in the local estimations of recombination rate when using mis-assembled reference genome will in turn affect any analysis based on recombination maps or including linkage disequilibrium.

### Tandem repeats and the current limits for obtaining chromosome-wide contigs

We found a high occurrence of conserved tandem repeats in the honey bee genome, whose length and sequence conservation caused problems for scaffolding contigs into chromosomes, the ultimate goal being each chromosome covered by a single contig. Indeed, long stretches of such repeats were found at the boundaries between contigs. Luckily, the only large contig in the assembly, that could be placed on chromosome 7, but not oriented due to lack of sufficient genetic data, had different tandem repeats at each of its extremities, allowing to decide on a correct orientation. However, other regions may still be problematic, the most striking example being the region between 1 and 3 Mb on chromosome 10. In this region, the contigs are small (< 0.2 Mb) due to a high occurrence of tandem repeats, leading to difficulties in their ordering along the chromosome and their orientation. Moreover, these repeats appear mostly to belong to the highly conserved 371 bp family, preventing their use for contig mapping. This portion of chromosome 10 has also been described as difficult to assemble in other studies [51].

### General chromosome structure: telomeres, centromeres

Cytogenetic studies based on fluorescent *in situ* hybridization of *Alu*I and *Ava*I probes suggest that the honey bee genome is composed of one large metacentric and 15 acrocentric chromosomes [52]. This is to date still considered as the honey bee standard karyotype structure [34,35]. However, other data could question this structure, for instance the suggested positions of the centromeres based on sequence characteristics of the HAv3.1 genome assembly such as the (GC) % and the presence of *Alu*I and *Ava*I repeats on chromosomes 7, 8 and 11 in Wallberg et al. (2019) [34].

Regarding telomeres, we were not able to identify the TTAGG consensus sequences on all the 17 chromosome ends (two for the metacentric chromosome 1 and one for each of the other fifteen acrocentric chromosomes) where they were expected: none were detected on the right arm of chromosome 1 and on chromosomes 3, 12 and 15. Interestingly, some chromosomes also lacked TTAGG repeats in the HAv3.1 assembly, but these were not the same (chromosomes 5 and 11). These discrepancies can be due to problems in the assembly of these repeat regions, either due to variations in the sequence quality between the two datasets or to local variations in repeat content, rendering the assembly of varying difficulty due to biological reasons. It is interesting to note, that in the older assemblies of the bee genome, based on the same DH4 strain as HAv3.1, extended analyses of telomeric and subtelomeric repeats showed that some chromosomes were easier to analyse than others and that no TTAGG repeats were identified for chromosome 5 [35]. Taken altogether, although the current sequencing data supports the actual consensus karyotype structure, we didn’t find that the *Alu*I repeat elements [52] could be considered as a marker of telomeres, as when such repeats were detected at the extremity of a chromosome, this was at the opposite end from the TTAGG repeats (see specifically chromosome 11 in Fig2).

The question of the exact position of the centromeres is a more complex one: the centromeres would be expected at the middle of chromosome 1 and at the proximal end of each of the other chromosomes. The *Ava*I repeat element, considered as a marker of the centromeres [52] was not found on all chromosomes and even when found, the number of repeats in the array could be as small as four, such as the repeat on chromosome 1 (Fig. 2). With the exception of chromosome 11 for which an *Ava*1 repeat was found at the position 5 Mb, the *Ava*1 elements, when detected on a chromosome, were found within 2.5 Mb of the chromosome ends, reflecting the results found on HAv3.1 [34]. However, although the positions of the *Ava*I repeats is identical between the two assemblies, the number of repeat elements vary for each given position. For the moment, the exact position of the centromeres remains uncertain, but the criteria of the eventual presence of an *Ava*I element remains a plausible indication, especially as these seem to be coincident with other specific characteristics, such as low (GC) content [50] or low levels of polymorphism and recombination rates [46]. If these characteristics are indicators of the centromere positions, then chromosome 11 and perhaps also chromosome 7 should be considered sub-metacentric, although in this case, TTAGG repeats would be expected at both of the extremities of these chromosomes, which is not the case in any of the studies to date. Further improvements in genome sequencing and assembly and in obtaining higher-resolution cytogenetic metaphase chromosome preparations will be necessary to elucidate this question.

### Comparing the genomes of two honey bee subspecies

The HAv3.1 assembly is based on a sample from the DH4 line, thought to be mainly of *A. m. ligustica* descent [9]. The comparison with our *Apis mellifera mellifera* AMelMel1.1 assembly allows for the detection of rearrangements occurring between these two distinct genetic types, that can’t be detected through short read sequencing.

Short sequence fragments repeated in tandem, such as the 91 bp and 371 bp repeats described here, tend to vary in copy number through non-allelic homologous recombination (NAHR) or unequal crossing-over [53]. A rapid observation of the LAST alignment data between the two assemblies suggests that the 371 bp repeat element can vary greatly in copy number and the 91 bp element to a much lesser extent, although these preliminary observations will require more thorough analyses. No obvious function was found for these elements to date, except for the fact that a BLAST search found that the 91 bp element shows similarity of sequence to one out of two exons of *Apis dorsata* and *Apis florea* lncRNAs, suggesting these are incomplete and consequently not active in the repeat arrays. However, the annotation of the lncRNAs in *Apis dorsata* and *Apis florea* is only based on the alignment of short reads RNA-seq. More work is needed to confirm this finding concerning the 91 bp repeat and further comparisons with other bee genomes whose sequences are underway [54] will help understand these interesting genome elements. The 5S ribosomal RNA genes are another interesting case of variation in gene number and studies in mouse and human have shown that this variation may be important for a balanced dosage of rRNA, that can have possible implications in diseases [37,38]. It would be interesting to see if the variations of 5S gene numbers observed here is a difference between the two honey bee subspecies investigated or if intra-population variation can be found.

After screening out rearrangements that could be due to errors associated with assembly problems, such as inversions of complete small contigs, thirteen inversions larger than 1 kb were detected between the two genomes. Out of these, a large 1.6 Mb inversion on chromosome 7 is likely an error in HAv3.1, as it was also seen when sequencing a closely related sample from the *Apis mellifera ligustica* subspecies [40]. Out of the twelve remaining inversions, some involve genes, present either at one of the breakpoints, having inversions within their structure (usually introns) or whose structure remains intact, but are in reverse orientation. As usual, interesting functions that may explain some of the phenotypic differences found between the two subspecies represented by our dataset will be found (see Additional file 2: Table S10). Even when restricting to genes for which functions were observed in insects, three genes stand out. One is *Bric-a-brac 1-like*, whose implication in body pigmentation in *Drosophila* [41], could be linked to our two reference genomes representing light (yellow) and dark coloured honey bee subspecies. Another is *Rhomboid*, previously shown to be involved in the formation in wing veins in *Drosophila* [42]. A third is the hormone receptor *HR38*, shown to be involved in the synchronisation of reproductive activity in the moth *Agrotis ipsilon* and the larval-pupal transition in the Colorado potato beetle *Leptinotarsa decemlineata* [43,44].

### Nuclear mitochondrial DNA segments and perspectives for pangenomics

To test the utility of having two reference genomes for genotyping structural variants, we tried genotyping two NUMTs, present in one or the other HAv3.1 and AMelMel1.1 assembly, with Graphtyper2. Results show that Graphtyper2 could not call genotypes for all samples. In the first instance, this is surprising, given the fact that this test of presence or absence of a 745 bp fragment in the case of NUMT_Chr2 and a 576 bp one for NUMT_Chr10 is done on haploid samples, simplifying the problem, as each of the NUMTs should be either present or absent in each individual tested. This may be caused by the fact that Graphtyper2 extracts reads that were previously mapped to the structural variant regions on a linear reference genome, thus possibly introducing a bias. It is however surprising, that when HAv3.1 was used as reference for the primary mapping of reads, NUMT_Chr10 could not be genotyped at all. This reference-bias could be overcome by using more recent methods in which the reads for the genomes to genotype are mapped directly on the pan-genome graph, although such methods are more complex to use in practice, due to problems such as the definition of genome coordinates [55].

## Conclusions

In conclusion, we present here a genome assembly for the honey bee *Apis mellifera* that is from a different subspecies than the current reference genome. One originality of the assembly process was to use recombination data rather than optical maps or HiC for scaffolding contigs into chromosomes. We characterise for the first time long tandem repeats that are present in the genome and are responsible for most sequence discontinuities and show that these belong to two main repeats families yet to be further characterised and whose potential function in the genome remains to be investigated. Finally, we show the interest of having two reference-quality genomes for the detection of structural variants, such as inversions and insertions-deletions and demonstrate the possibility of using a pan-genome approach for genotyping such variants in honey bee populations.

## Declarations

### Ethics approval and consent to participate

Not applicable

### Consent for publication

Not applicable

### Availability of data and materials

The AMelMel1.1 assembly has been deposited on the NCBI under the accession number GCA_003314205. The reads of the 36 corresponding PACBIO_SMRT runs are in SRA under the accessions SRR9587836 to SRR9593684. Scripts and supplementary description of bioinformatic analyses are available in GitHub: https://github.com/avignal5/PacificBee/tree/main.

### Authors’ contributions

KC-T, WM and AV performed sampling and high molecular weight DNA extraction; CV, CR, CD performed the sequencing; CK performed assembly into contigs and contig quality checks; SEE, BS and AV did the chromosome-level assembly using recombination data; AV did the tandem repeats analyses; QB and AV did the NUMT detection and analysis. SEE and AV drafted the manuscript.

All authors read and approved the final manuscript.

## Additional files

**Additional file 1 Supplementary methods and Supplementary Figures S1-S18**

Format: pdf

Title: Supplementary methods and supplementary figures S1-S11

Description:

**Additional file 2 Tables S1-S11**

Format: Excel file

Title: Supplementary tables S1-S11

Description:

**Additional file 3 AMelMel-Hav3**

Format: pdf

Title: Comparison of AMelMel1.1 and HAv3.1 genome assemblies.

Description: Dot plot alignments of the AMelMel1.1 and HAv3.1 genome assemblies.

**Additional file 4 AMelMel-Amel4_5**

Format: pdf

Title: Comparison of AMelMel1.1 and Amel4.5 assemblies

Description: Dot plot alignments of the AMelMel1.1 and HAv3.1 genome assemblies.

**Additional file 5 Inversions**

Format: pdf

Title: Inversions larger than 1 kb detected between the AMelMel1.1 and HAv3.1 genome assemblies.

Description: Inversion structural variants larger than 1 kb, detected after aligning the AMelMel1.1 and HAv3.1 genome assemblies with LAST.

## Supporting information

Alignment of AMelMel to HAv3.1

Alignment of AMelMel to Amel4.5

Chromosomal inversions between AMelMel and HAv3.1

