## Supplemental figures for "The black honey bee genome: insights on specific structural elements and a first step towards pan-genomes"

### Supplementary methods

#### DNA extraction

To ensure a sufficient quantity of high molecular weight DNA was extracted for sequencing from a single drone, several samples were extracted using different protocols. We found that one of the most crucial steps was the grinding of the sample. And the most satisfying results were obtained using a mortar and potter (figure 1).

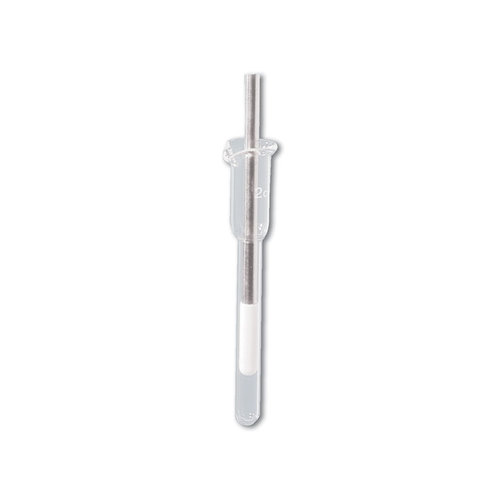

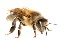

**Figure 1: mortar and potter used for grinding individual drones.** Our best results were obtained with two larval samples OUE7B (303 mg), OUE8B (336, 5 mg). After grinding, the samples underwent a 50 min lysis at 50°C followed by an overnight lysis at 37°C. OUE8B was lysed at 50°C for 3 h. After one centrifugation step, the DNA was immobilized on QIAGEN Genomic-tips 100/G kit (Cat No./ID: 10243). After several washing steps, DNA is eluted from the column, then desalted and concentrated by alcohol precipitation. The DNA was resuspended in TE buffer. DNA quantification was calculated using a Qubit 3.0 Fluorometer assay (Qubit dsDNA BR Assay Kit) while size distribution and degradation assessed using a Fragment analyzer (AATI) (figures 2A and 2B).

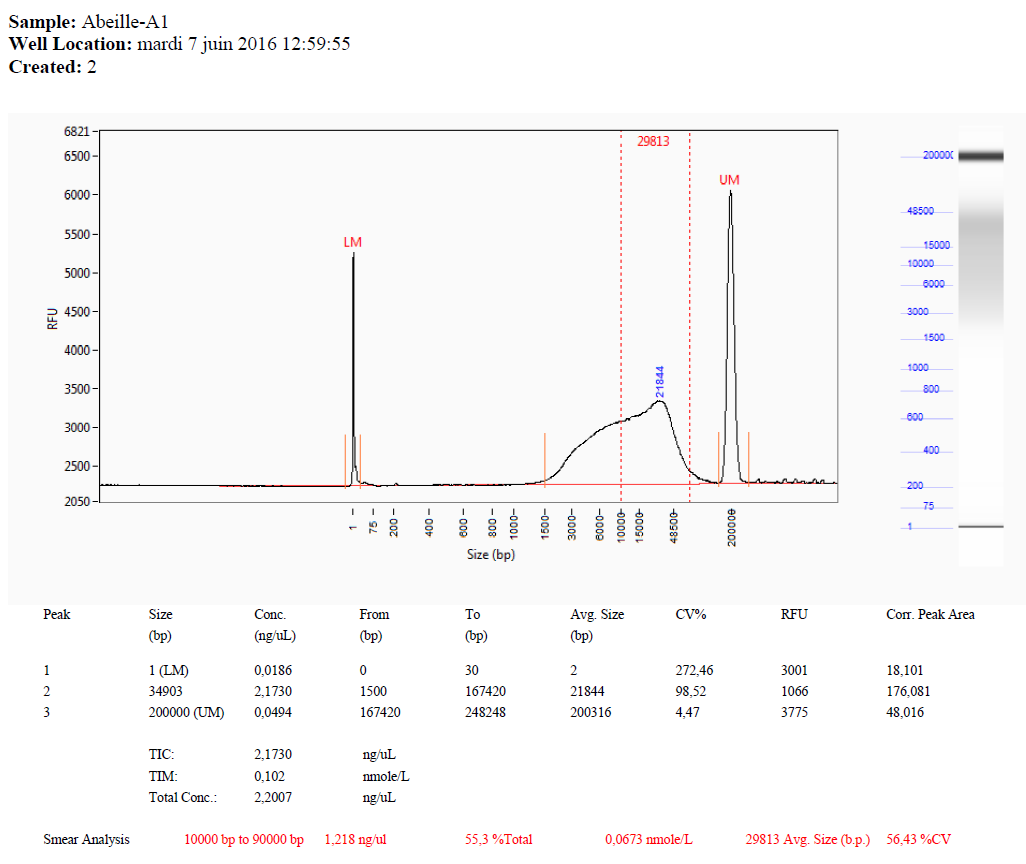

**Figure 2A**: Fragment analyzer results for sample OUE7B.

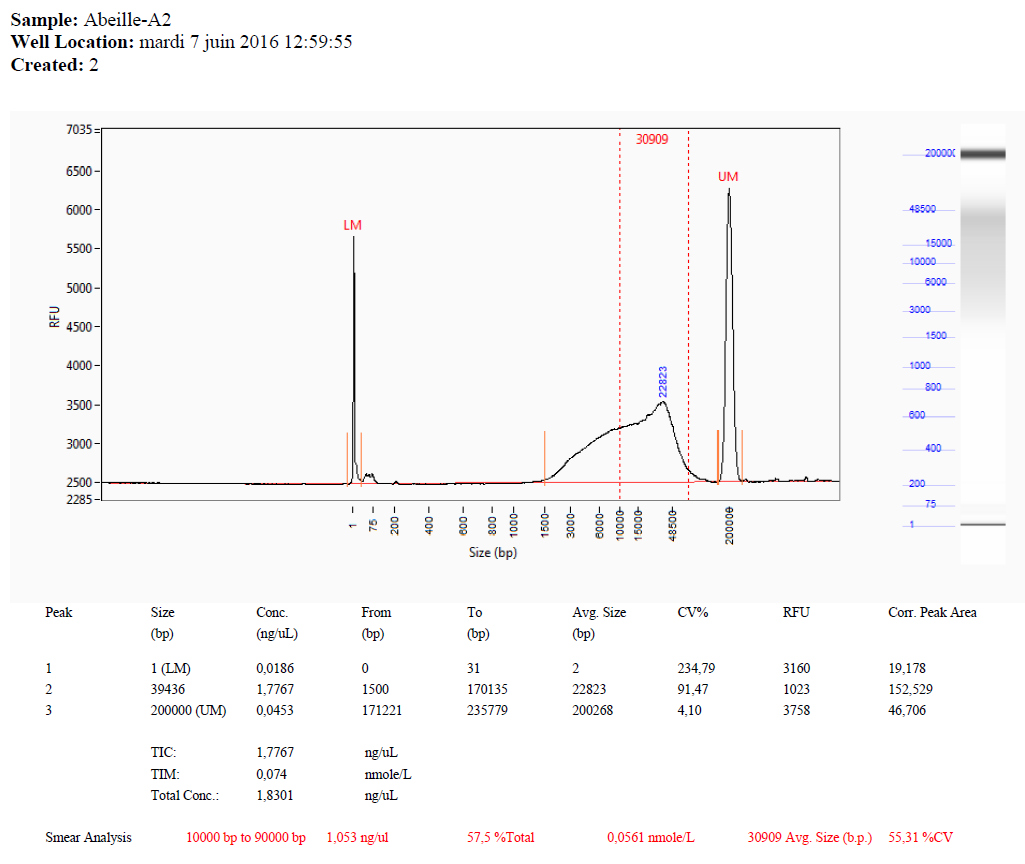

**Figure 2B: Fragment analyzer results for sample OUE8B.** The size of the DNA fragments was satisfactory for both samples and the quantities of DNA obtained for the 2 samples being equivalent: 100 ng/µL and 110 ng/µL for OUE7B and OUE8b respectively, which is largely sufficient for the construction of libraries for all the necessary sequencing runs, we chose to carry the construction of libraries using only the OUE7B sample.

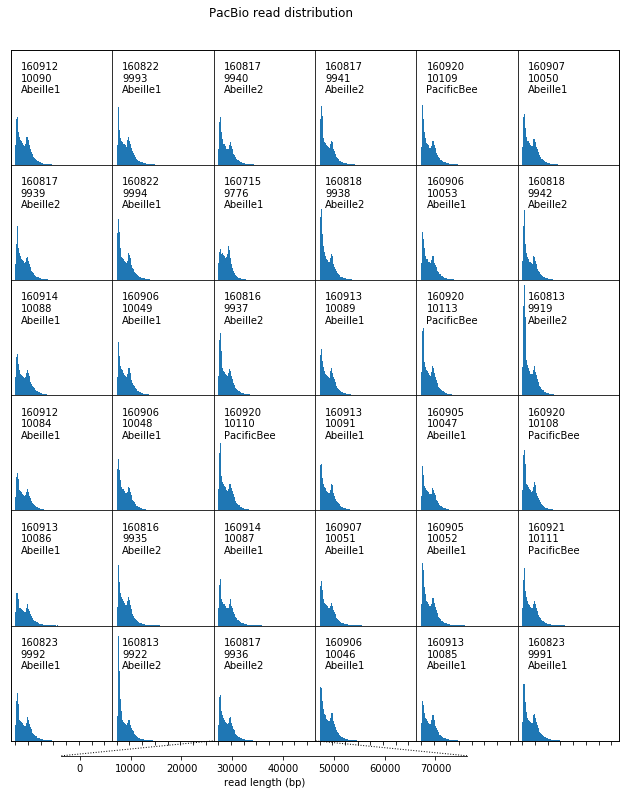

**Figure 3: distribution of read lengths per SMRTcell.** Three libraries “PacificBee, “Abeille1”, “Abeille2”, all constructed from the same drone samples OUE7B, were sequenced on 36 SMRTcells on a RSII instrument.

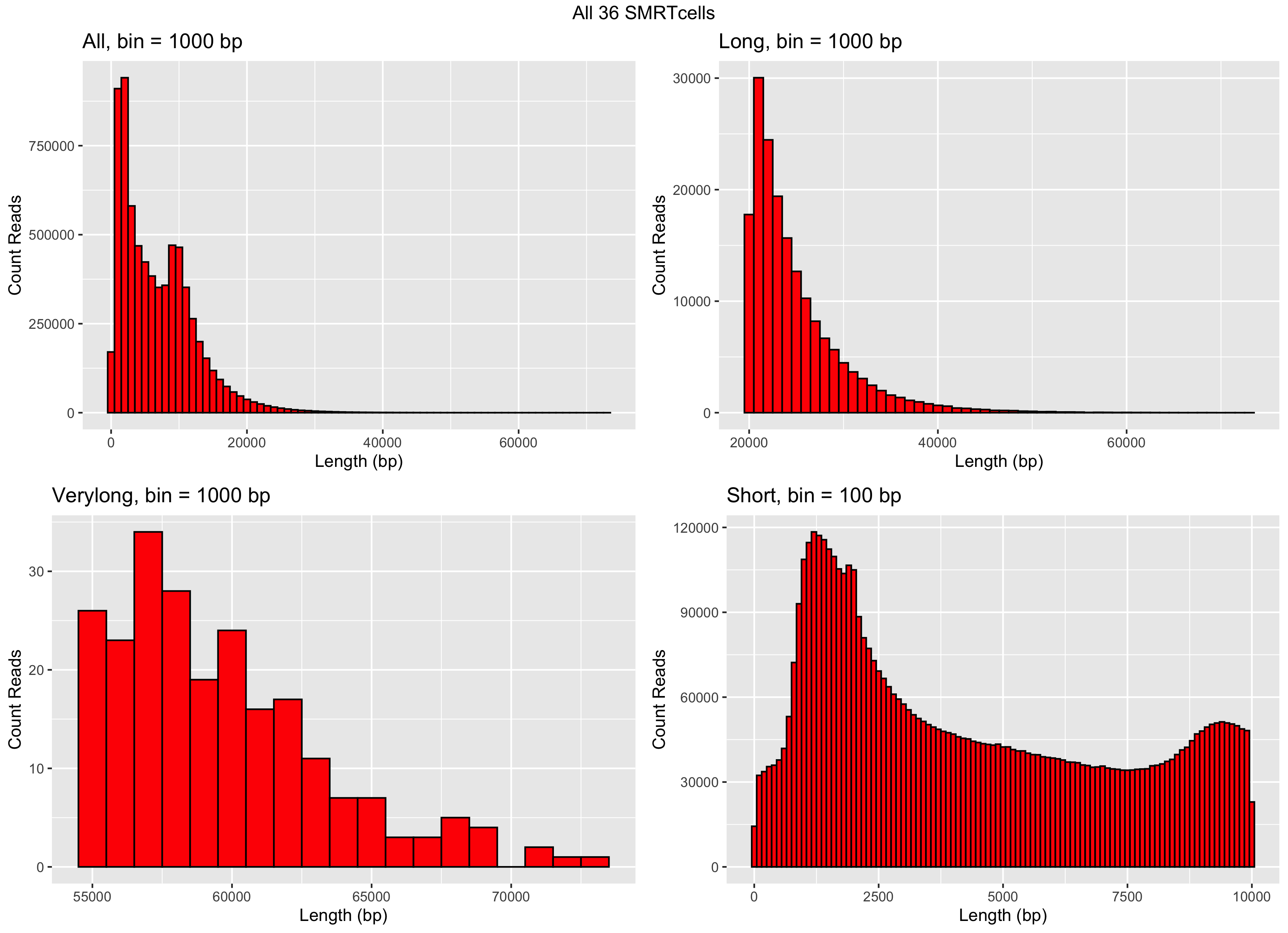

**Figure 4: Size distribution of sequencing reads from all 36 SMRTcells.** A few reads (four) have a length in excess of 70 kb.

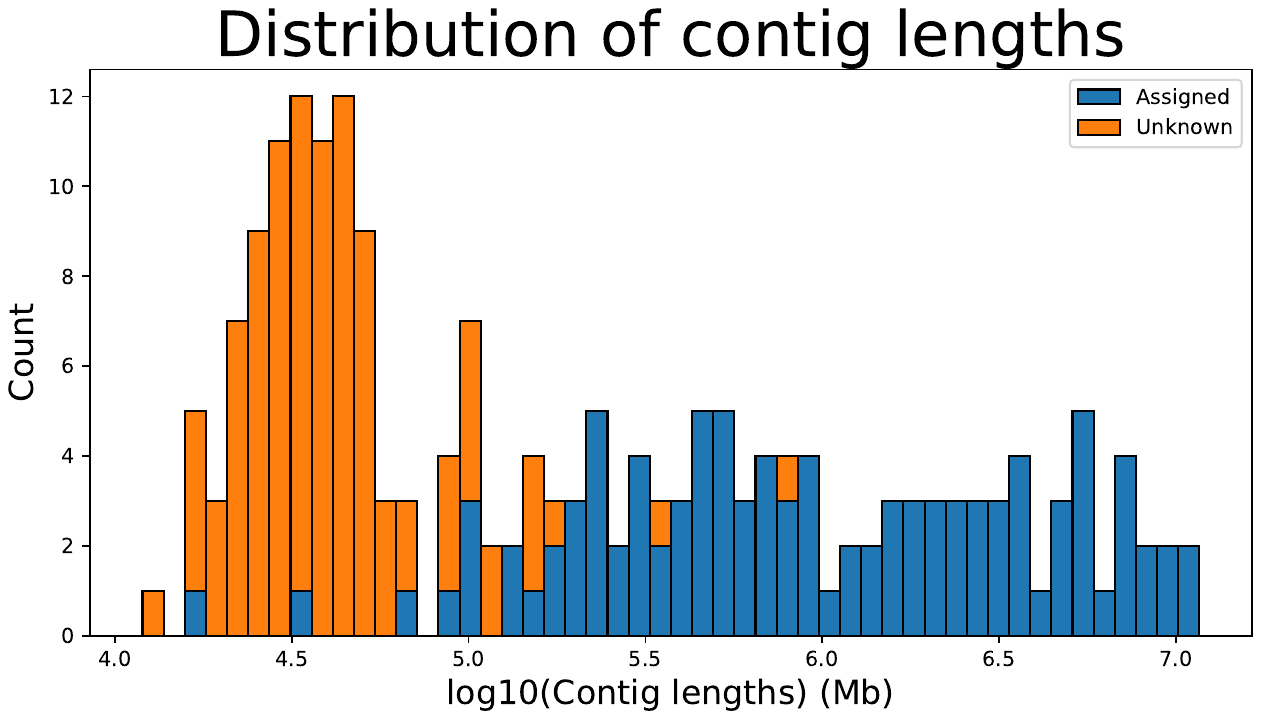

**Figure 5: Size distribution of the length of the contigs assembled with Canu.** Blue: contigs assigned to chromosomes. Orange: contigs that couldn’t be assigned to chromosomes.

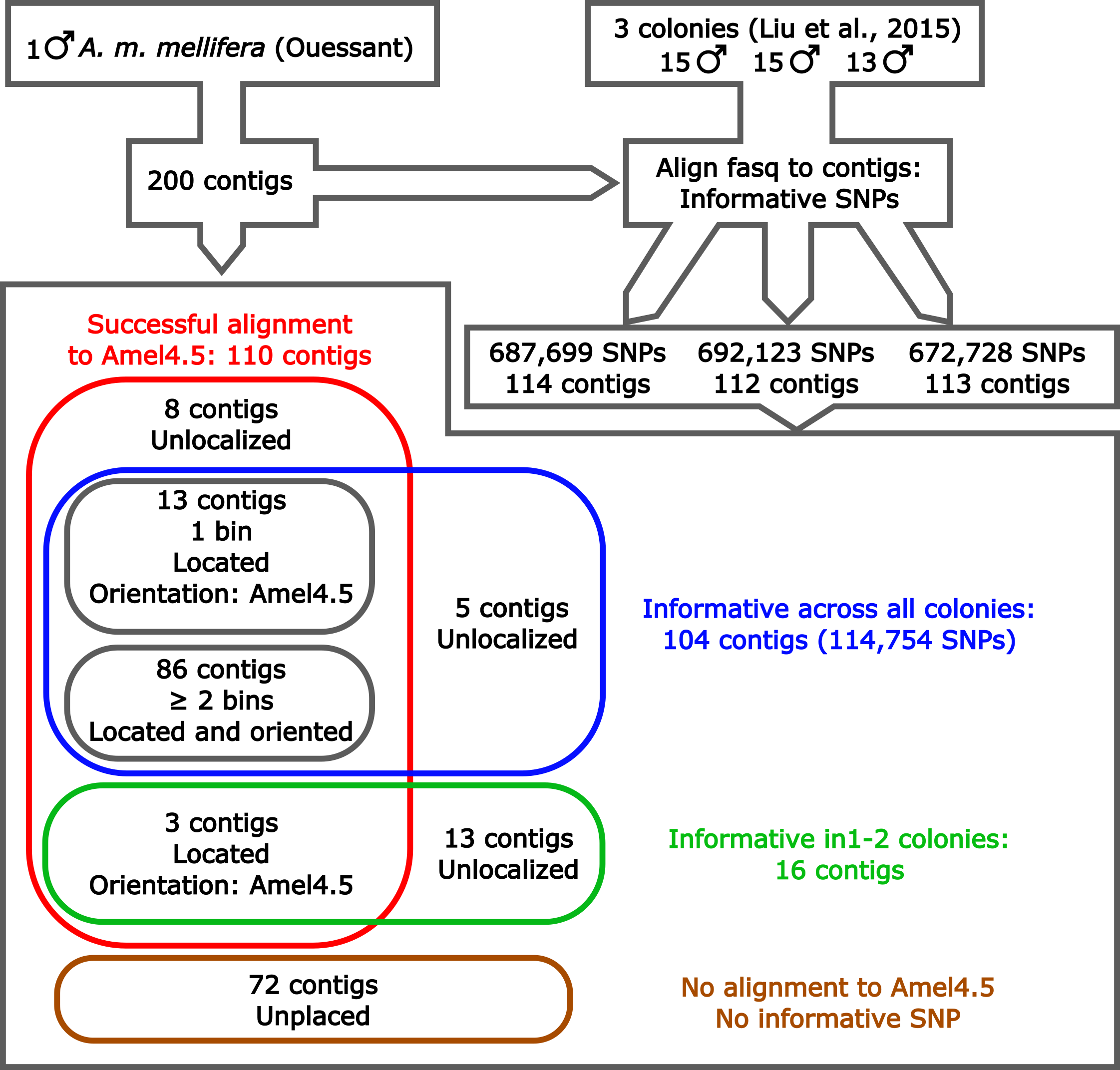

**Figure 6: Alignment of contigs to Amel4.5 and detection of SNPs in the data from Liu et al., (2015).** One hundred and ten contigs were assigned by alignment to Amel4.5 and 120 were informative in at least one of the three colonies from Liu et al (2025). Out of these, one hundred and two could be ordered along the chromosomes by using the crossing-over data and 86 having at least one crossing-over event detected within the contig (≥ 2 genotype vector bins), could be oriented using this same data.

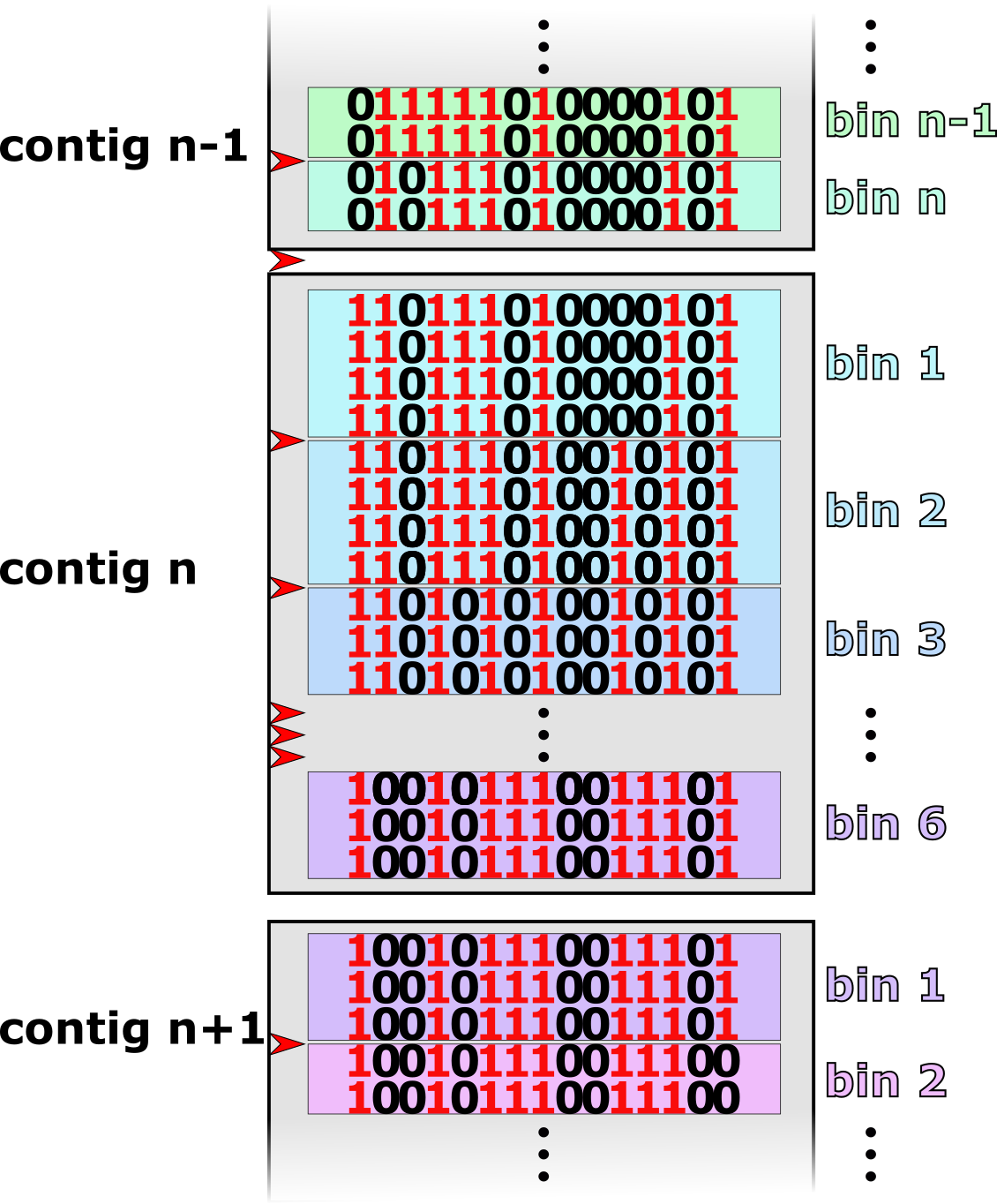

**Figure 7: Ordering contigs by using crossing-over events.** Vectors of ‘0’ and ‘1’ represent genotypes detected for drones from a same colony. These allow to detect crossing-over events (red arrows) having occurred in the queen gametes having produced the drones. To facilitate analyses, successive identical vectors were grouped together into bins. Within-contig crossing-overs can be observed and used to estimate the cM/Mb ratio in the genome. Bins from contig extremities are used to order contigs by the detection of contig ends from different contigs having minimal crossing-over events. For instance in the example shown, no crossing-over was detected between contig n and contig n+1 and one crossing-over was detected between contig n-1 and contig n.

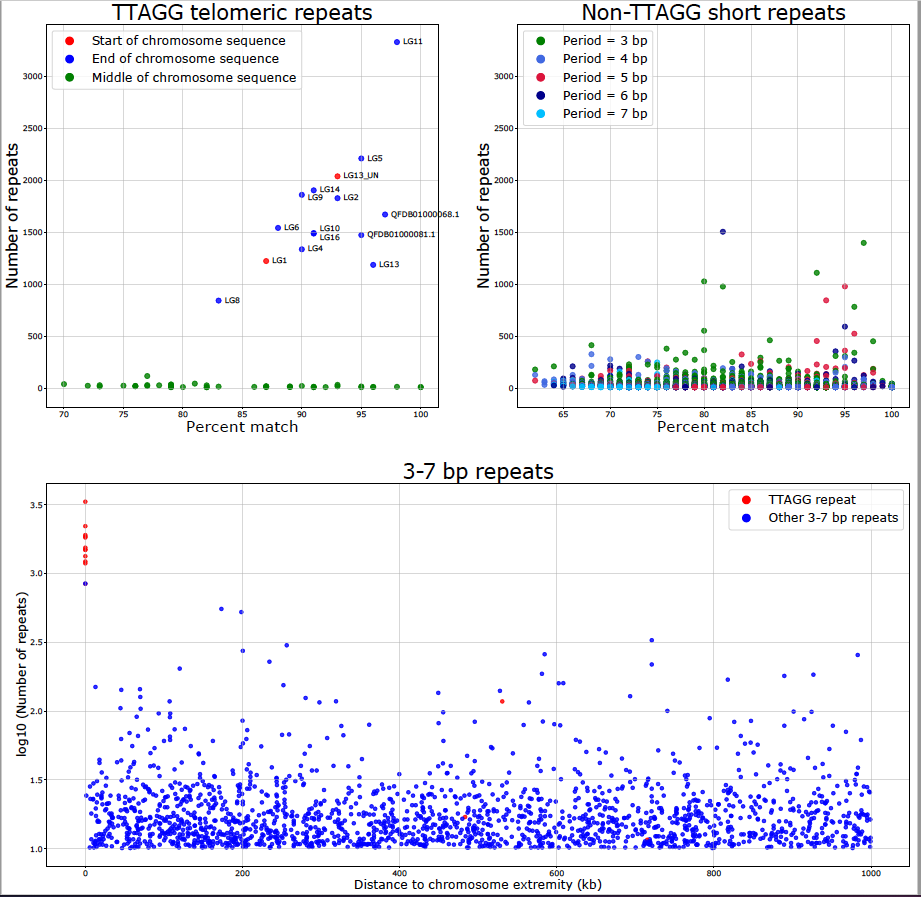

**Figure 8. Telomeric repeats and other repeats of small period size.** Top left: number of repetitions of the consensus TTAGG telomeric repeat; red: at the start, blue at the end and green in other positions of a chromosome sequence. Top right: The number of repetitions of other short repeats. Bottom: distance to the chromosome end for TTAGG and other short repeats. The only non-TTAGG repeat with a high number of copies present at the end of a chromosome is an AATAT motif repeated 845 times with 93% matches.

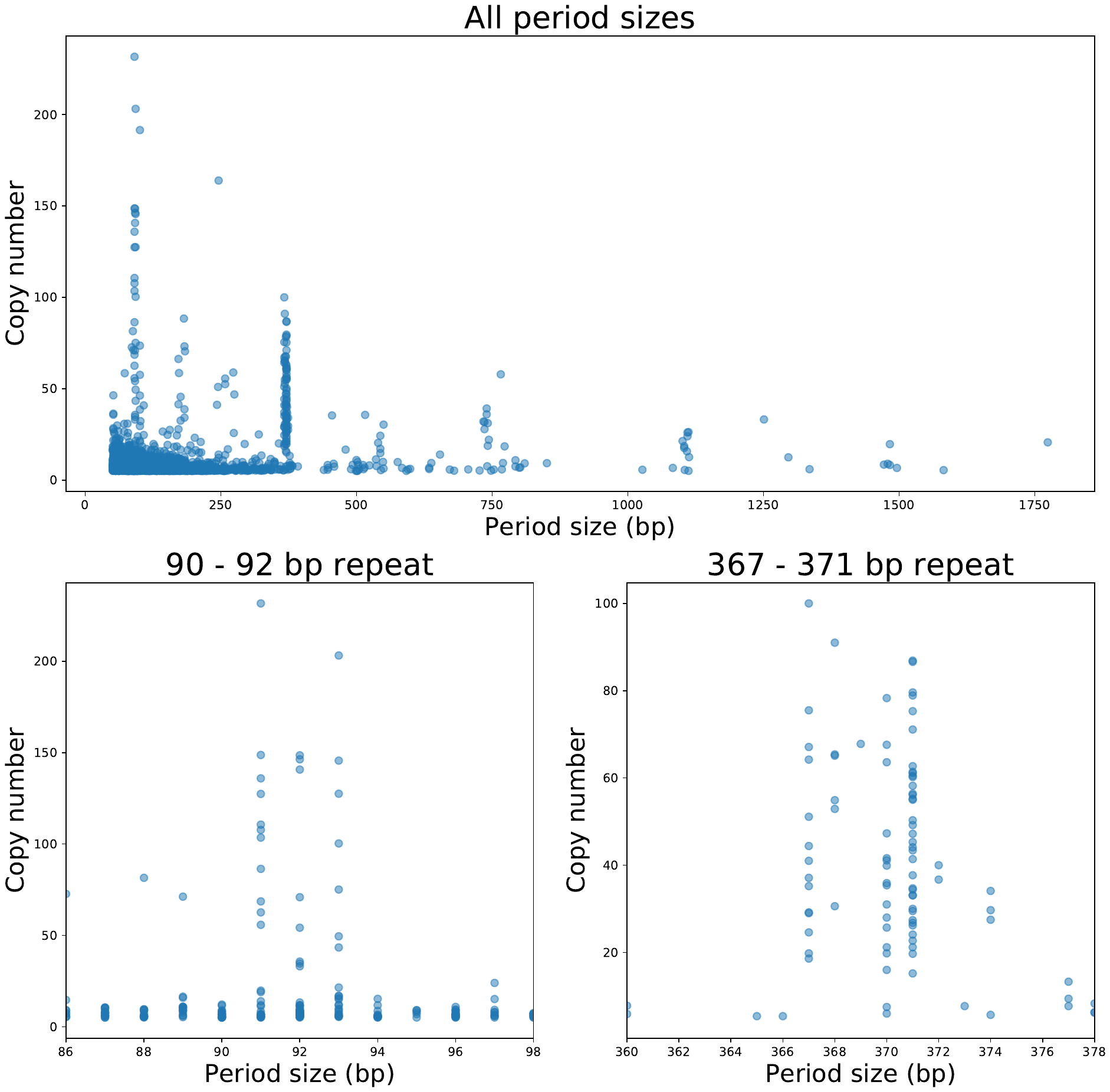

**Figure 9: Period size and copy number of tandem repeats.** Top: period sizes between 50 and 2000 bp, as detected by Tandem repeat Finder. Bottom left: close-up on the peak at 91-93 bp. Bottom right: close-up on the peak at 367-371 bp.

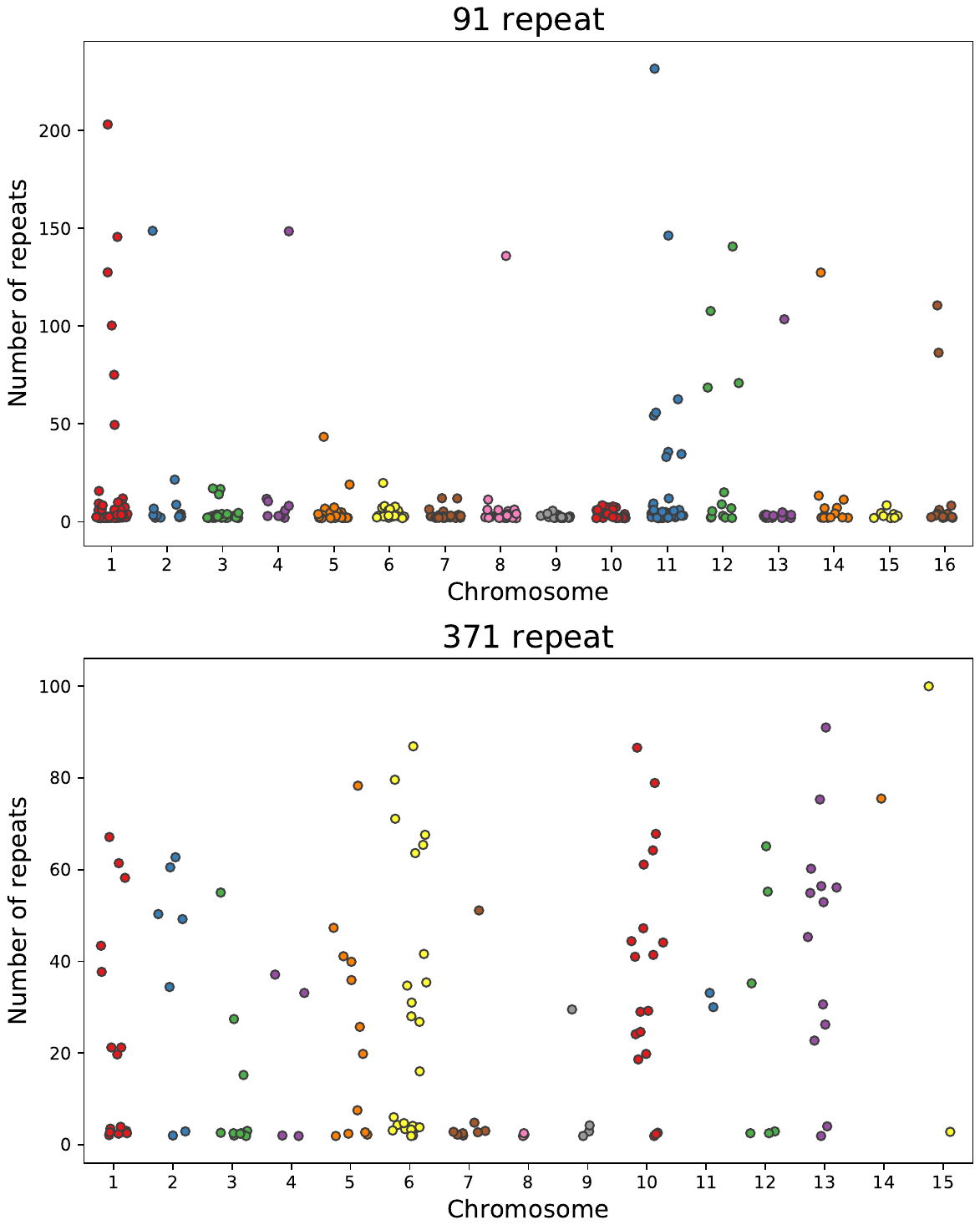

**Figure 10: Chromosome distribution of the 91 bp repeat and the 371 bp repeat.** The 91 repeat corresponds to the size range 91-93 and the 371 repeats to the size range 367-371. There are no 371 repeats on chromosome 16.

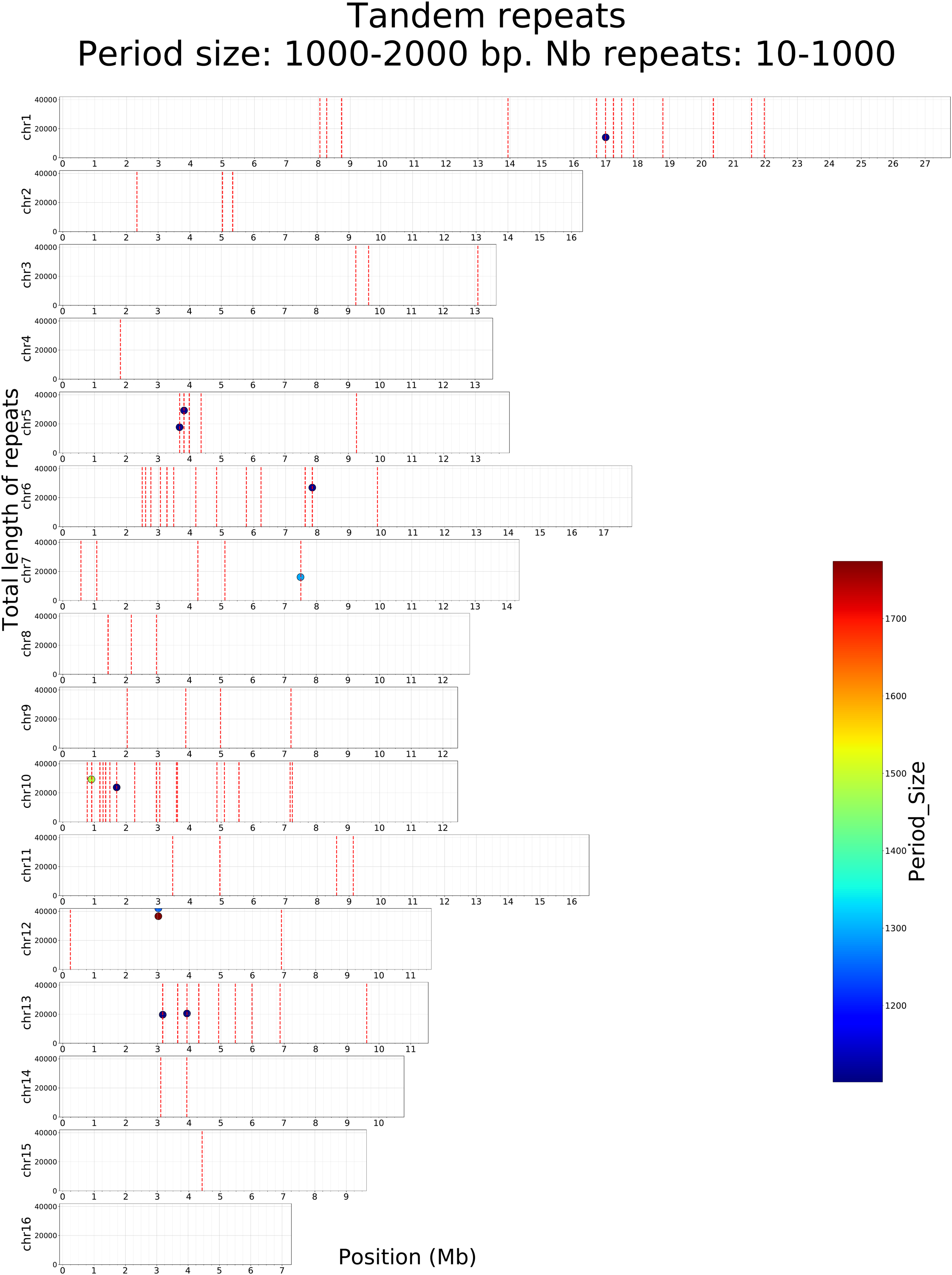

**Figure 11: Tandem repeats of period size 1000-2000 bp detected in the AMelMel assembly.** The color scale represents the period size of the repeat elements and the Y axis the total length of the repeat array. Vertical dotted lines represent the contig boundaries in the AMelMel1.1 assembly.

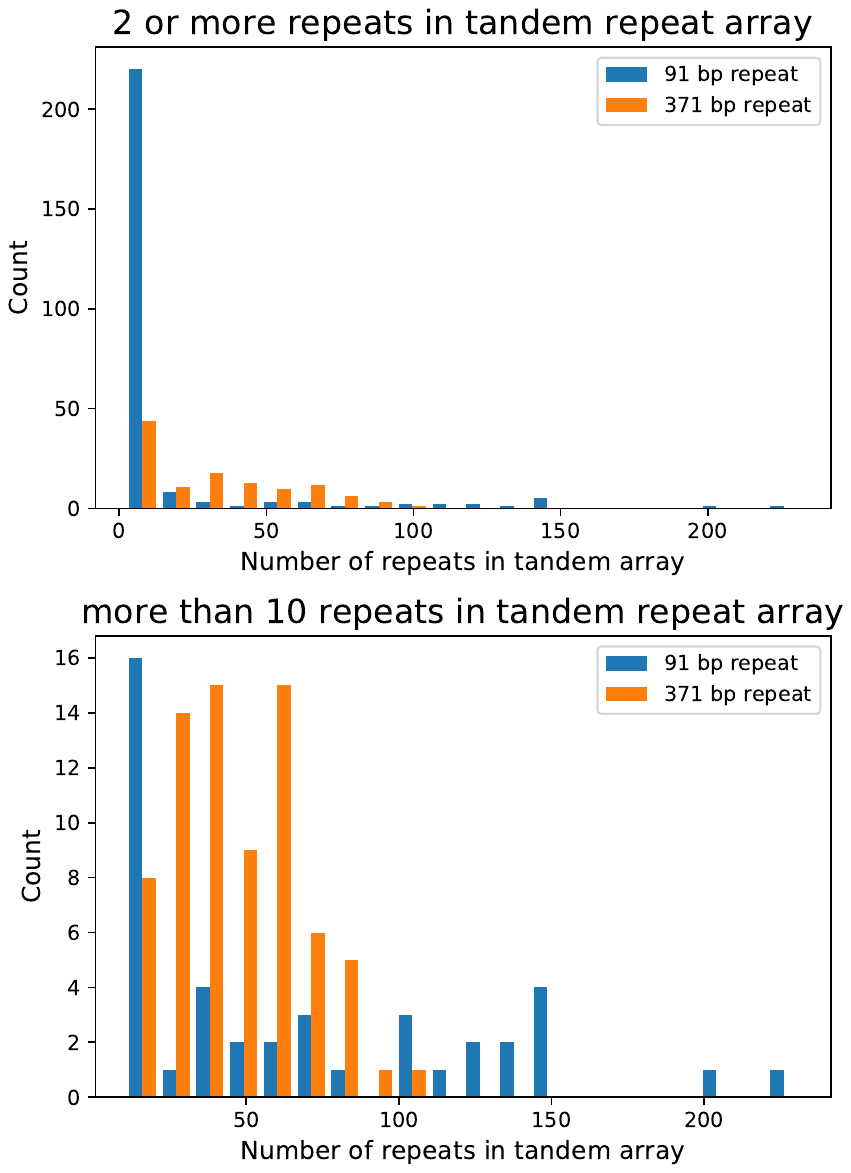

**Figure 12: Distribution of the number of repeats in tandem arrays for the 91 bp and 371 bp repeats.**

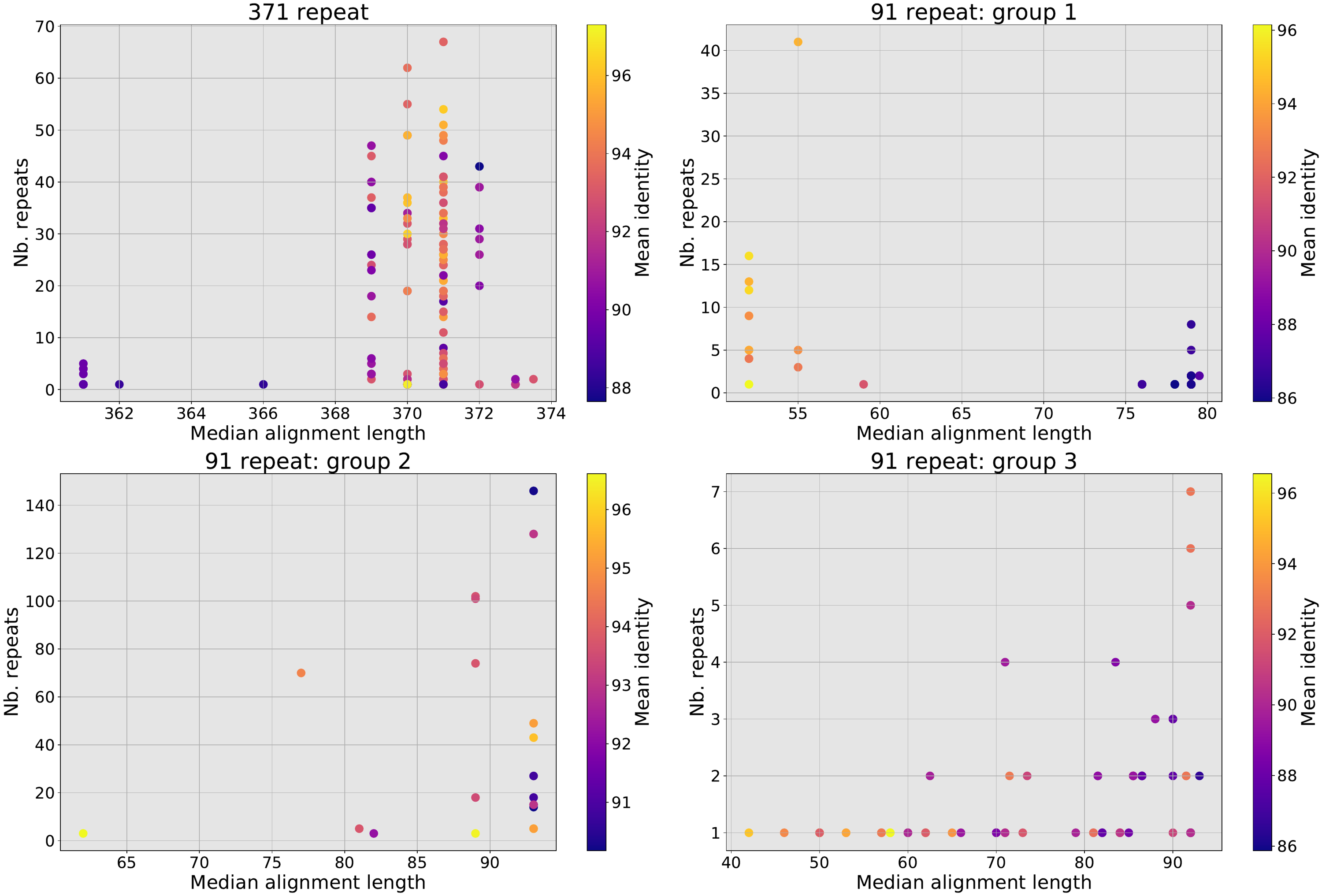

**Figure 13:** Summaries from the BLAST hits on AMelMel1.1 obtained with the consensus sequences from the major group for the 371 bp repeats and the three major groups for the 91 bp repeats. BLAST hits were grouped together when found in tandem arrays in the genome. Abscissa: median alignment length for each hit within tandem arrays, ordinate: number of repeats within tandem arrays, colour: mean identity within tandem arrays.

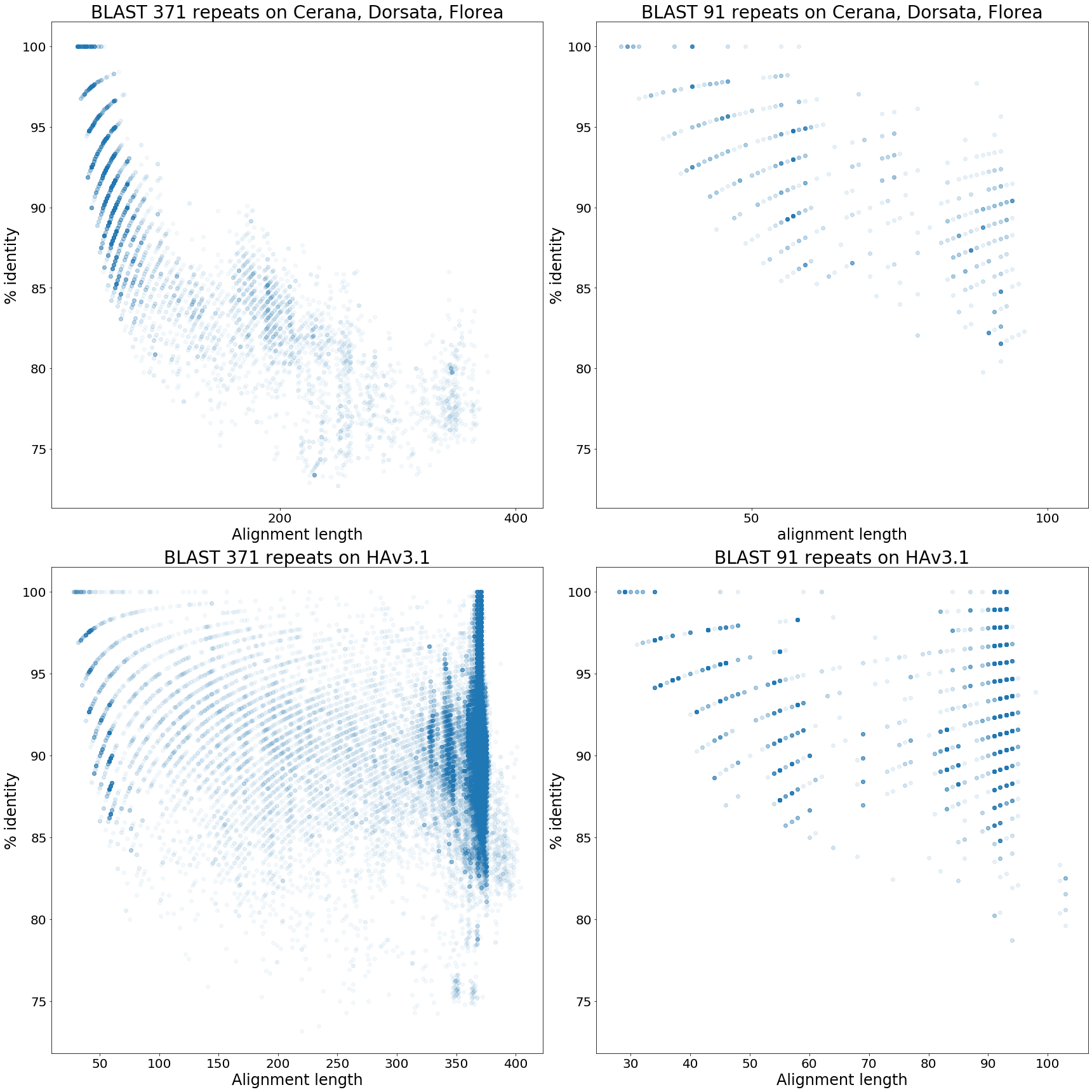

F**igure 14: BLAST searches with the 371 bp and the 91 bp on *Apis* genomes.** The graphs present the alignment lengths and % identities of all the hits of each of the 371 bp repeats (left) and the 91 bp repeats (right) on the *A. cerana*, *A. dorsata* and *A. florea* genome databases (top) and on the *A. mellifera* HAv3.1 assembly (bottom).

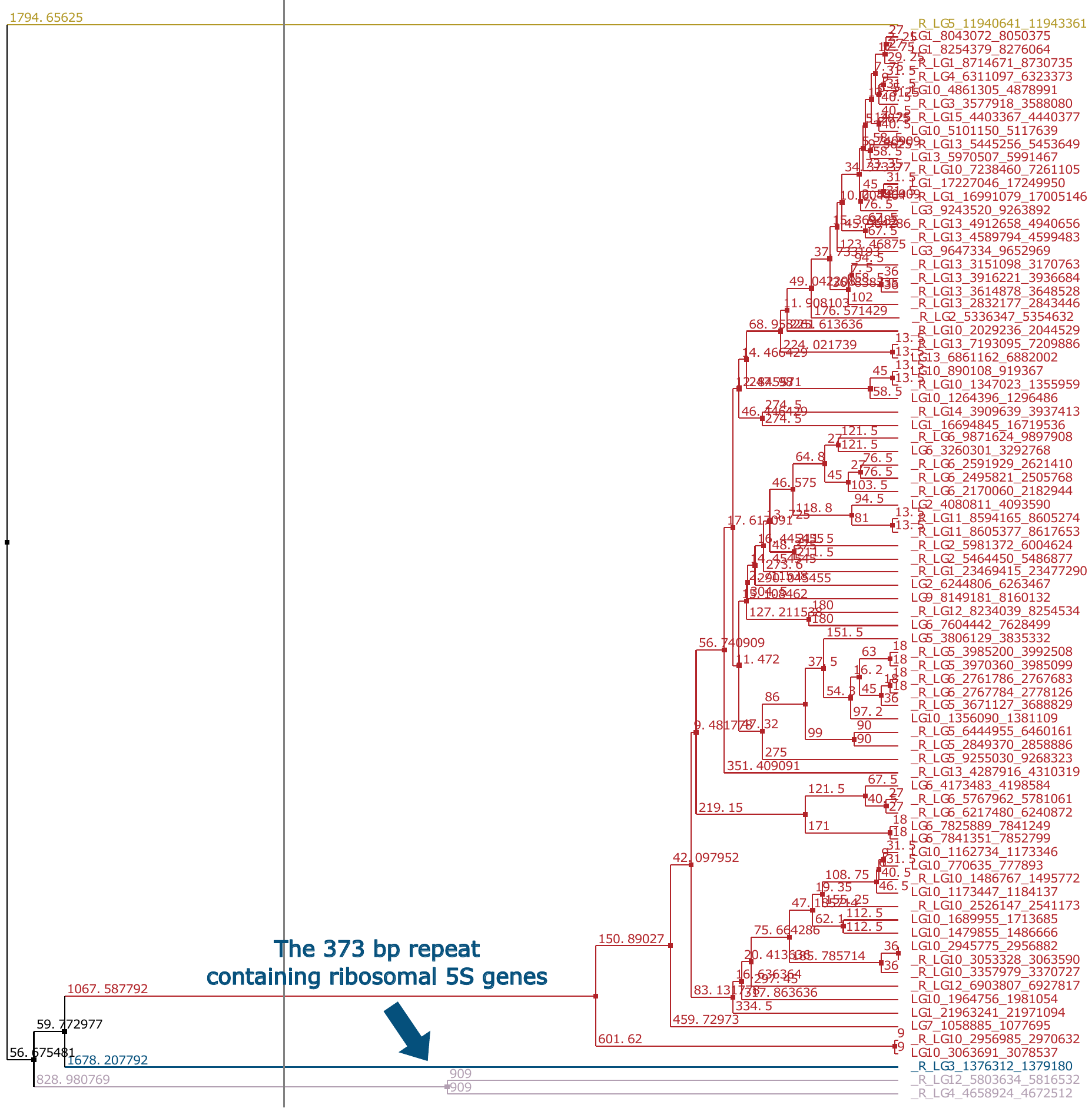

**Figure 15: Phylogenetic trees for the tandem repeats of period size 367-374 bp.** Tandem repeats with seven or more elements, such as detected by Tandem Repeat Finder, are included. The 373 bp repeat array containing 8 repeats of the 5S ribosomal genes is indicated by the blue arrow.

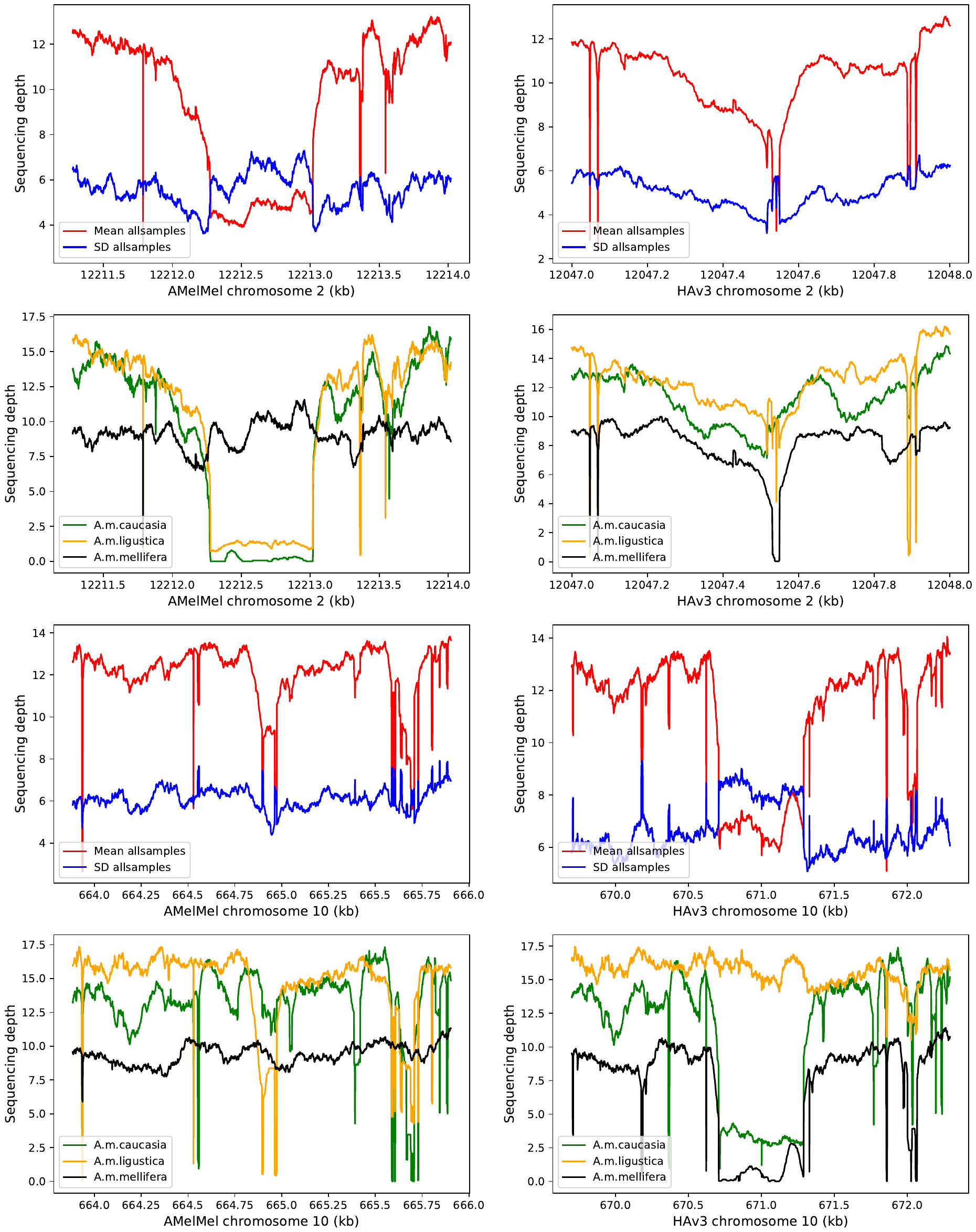

**Figure 16: Sequencing depth within and around insertions and deletions in *Apis mellifera* subspecies.** Left: a 745 bp variant present in AMelMel1.1 and absent in HAv3.1. Right: a 576 bp variant present in HAv3.1 and absent in AMelMel1.1. Red: mean sequencing depth for the alignment of 80 samples on AMelMel (left) and HAv3.1 (right). When the sequences are aligned to the reference genome having the insertion present, a drop of sequencing depth coincides with the position of the insertion, suggesting that a significant proportion of samples may lack the corresponding segment. Blue: standard deviation of the sequencing depth increases in the same region, confirming the heterogeneity of the samples for the presence or absence of the insertion. Mean sequencing depths are also indicated per subspecies, with *A. m. caucasia* (15 samples) in green, *A. m. ligustica* (30 samples) in yellow and *A. m. mellifera* (35 samples) in black. Results suggest most of the *A. m. mellifera* samples contain the insertion present in the AMelMel1.1 assembly on chromosome 2, as the sequencing depth remains constant throughout the region, and not the one present in the HAv3.1 assembly on chromosome 10, as indicated by a sequencing depth close to zero. Inversely, most of the *A. m. ligustica* samples contain the insertion present in the HAv3.1 assembly on chromosome 10 and not the one in the AMelMel1.1 assembly on chromosome 2. Most *A. m. caucasia* samples lack the insertion present in the AMelMel1.1 assembly and a few seem to have the insertion present in the HAv3.1 assembly.

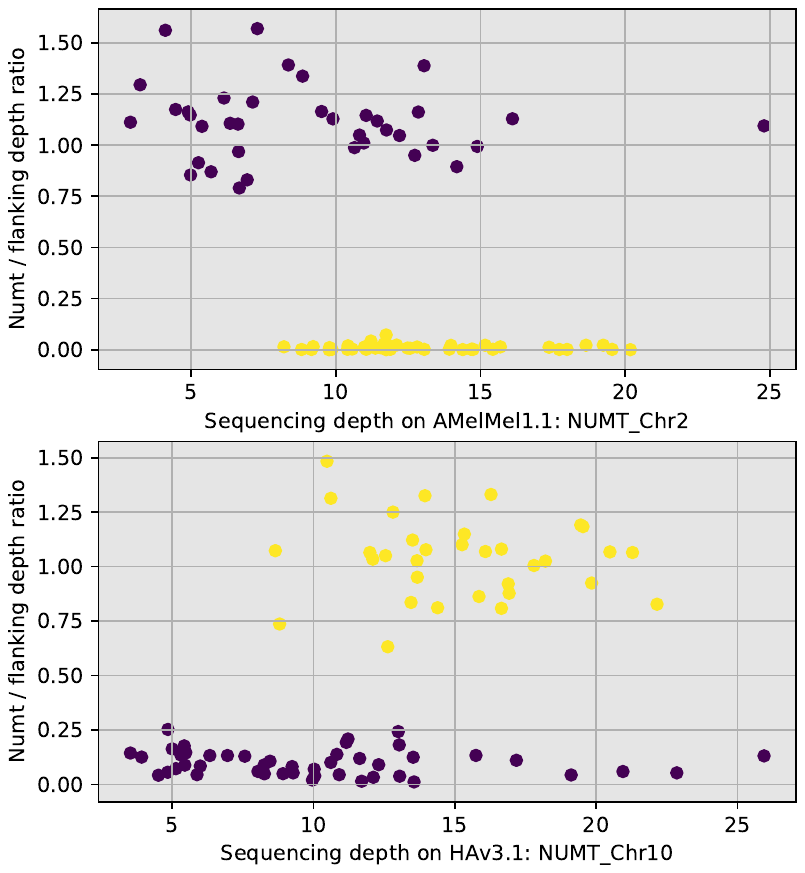

**Figure 17: Sequencing depth and genotype deduction in NUMT_Chr2 and NUMT_Chr10.** Each point represents a sample. X-axis: sequencing depth; Y-axis: ratio between the sequencing depth within the NUMT boundaries and in the flanking regions, using the alignments on the genome in which the NUMT is present. Colour represents the assignments by k-means clustering to the two groups considered.

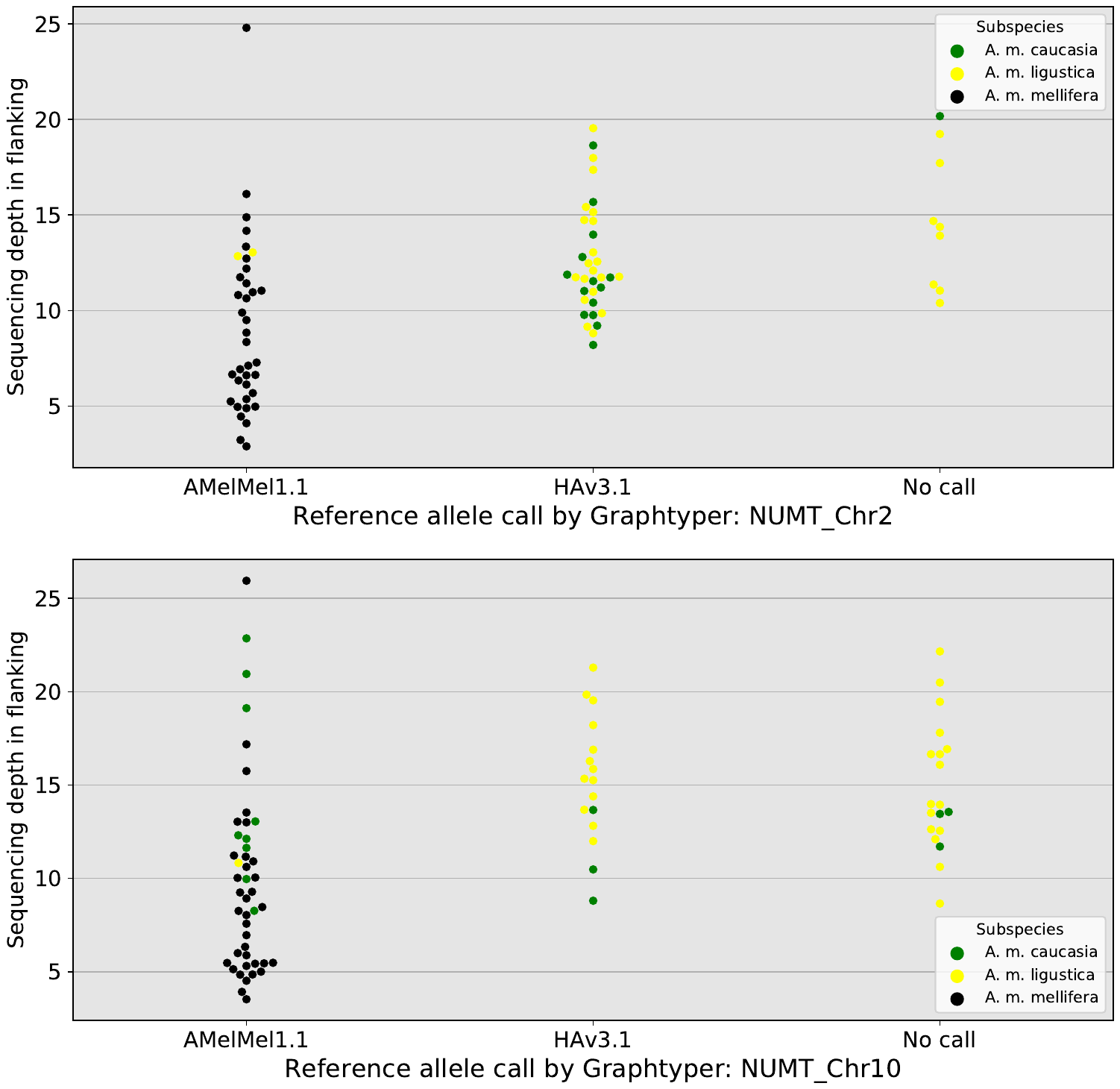

**Figure 18: Sequencing depth and call rate with Graphtyper 2.** Sequencing depth in flanking sequences does not seem to affect the call rate with Graphtyper 2, as some samples with a sequencing depth lower than 3 were successsfully called.
