## Supplementary material for "The black honey bee genome: insights on specific structural elements and a first step towards pan-genomes": Alignment of AMelMel to HAv3.1

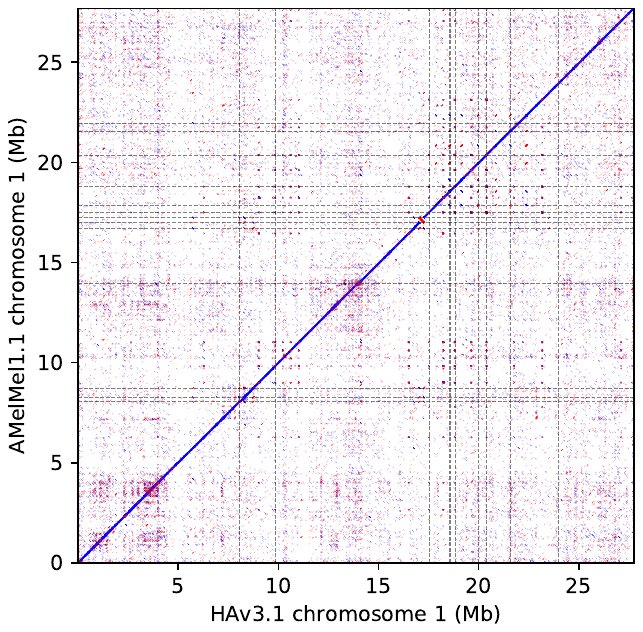

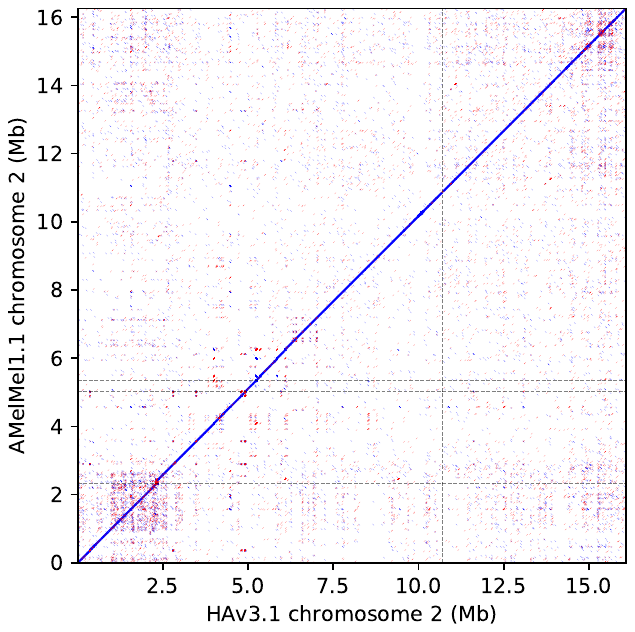
 **Chromosomes 1 and 2.** Dotplot representation of the alignment of HAv3.1 (x-axis) and AMelMel1.1 (y-axis). Blue: alignments in the same direction, red: alignments in inverted directions. Vertical and horizontal dotted lines represent contig boundaries on the two assemblies.

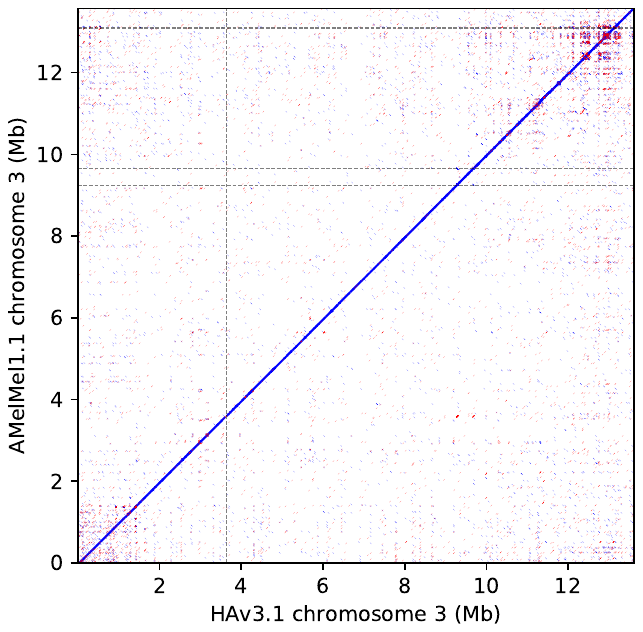

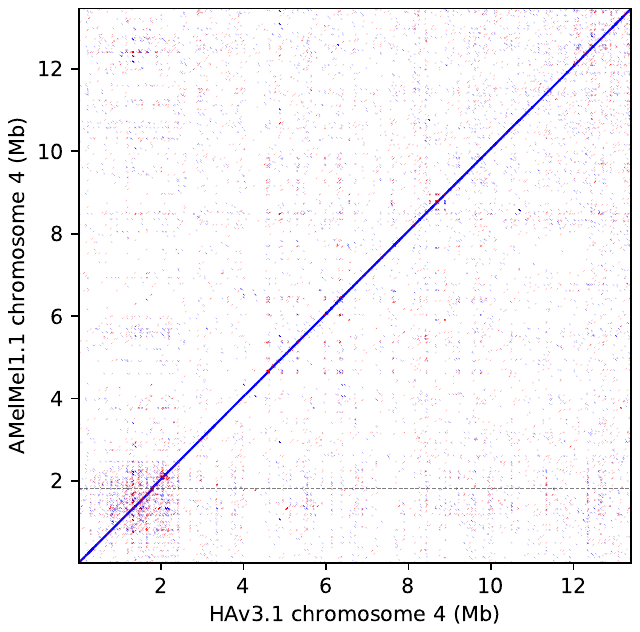
 **Chromosomes 3 and 4.** Dotplot representation of the alignment of HAv3.1 (x-axis) and AMelMel1.1 (y-axis). Blue: alignments in the same direction, red: alignments in inverted directions. Vertical and horizontal dotted lines represent contig boundaries on the two assemblies.

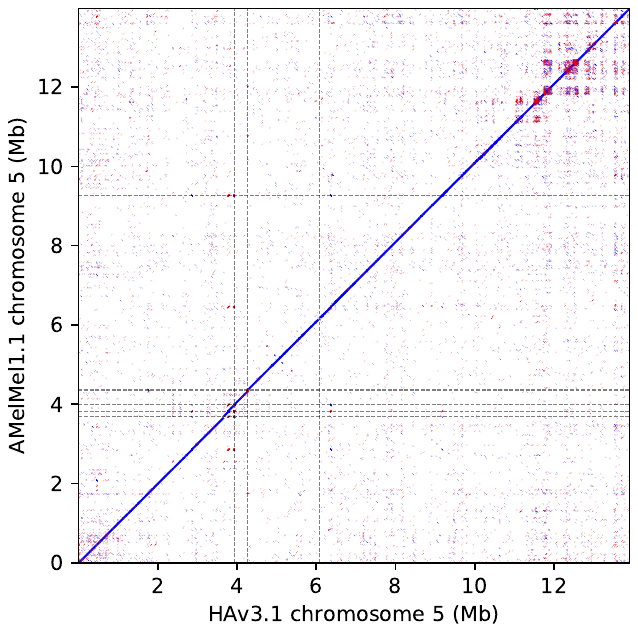

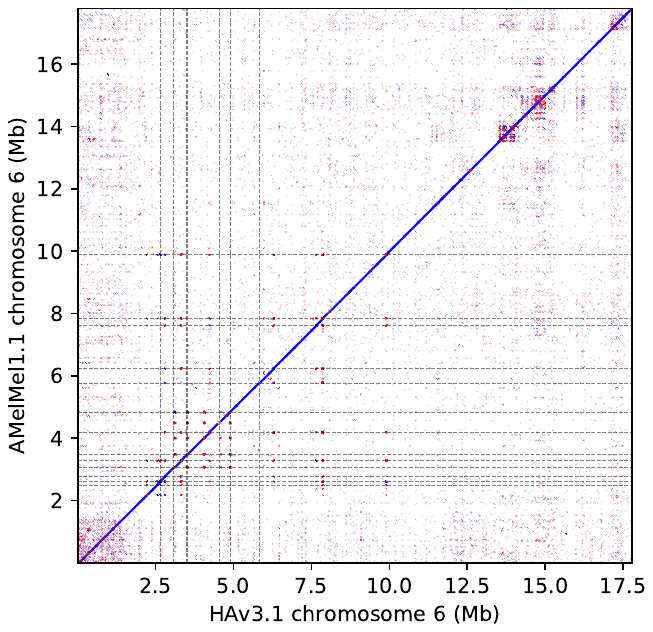
 **Chromosomes 5 and 6.** Dotplot representation of the alignment of HAv3.1 (x-axis) and AMelMel1.1 (y-axis). Blue: alignments in the same direction, red: alignments in inverted directions. Vertical and horizontal dotted lines represent contig boundaries on the two assemblies.
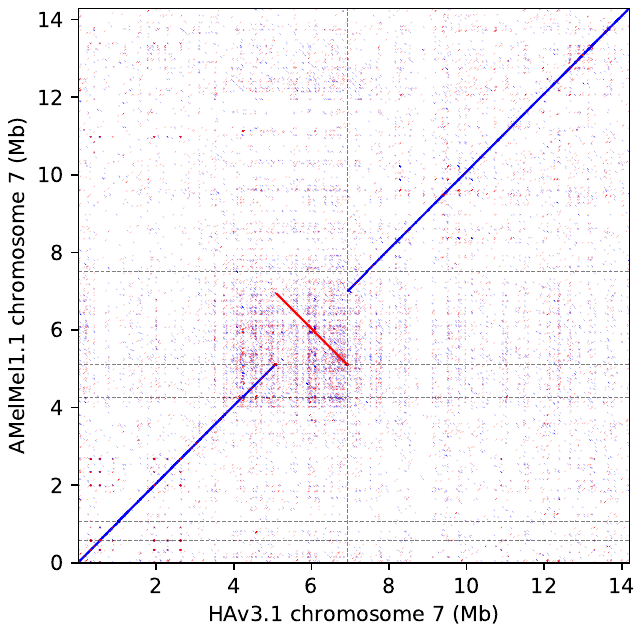

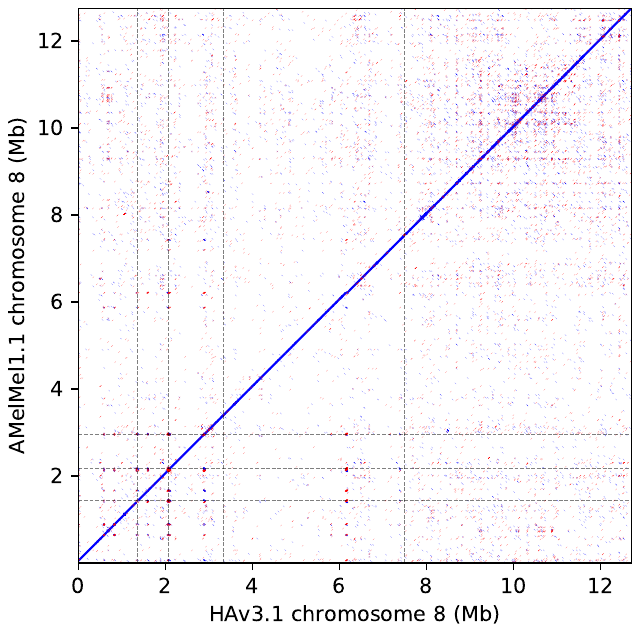
 **Chromosomes 7 and 8.** Dotplot representation of the alignment of HAv3.1 (x-axis) and AMelMel1.1 (y-axis). Blue: alignments in the same direction, red: alignments in inverted directions. Vertical and horizontal dotted lines represent contig boundaries on the two assemblies.
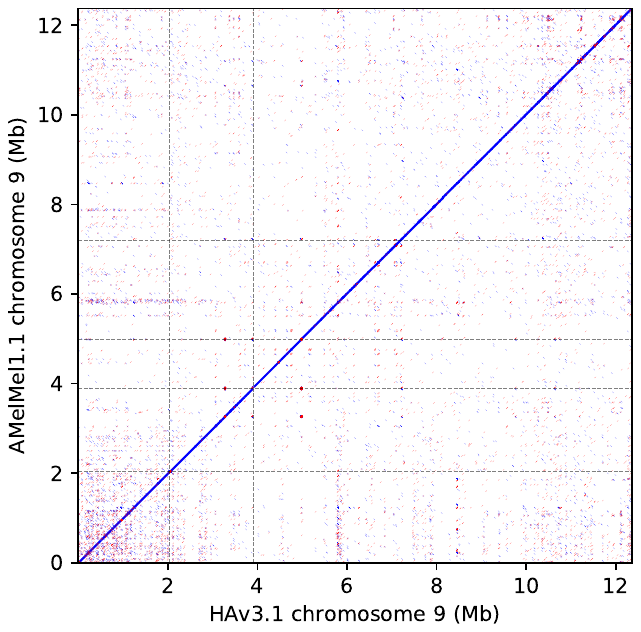

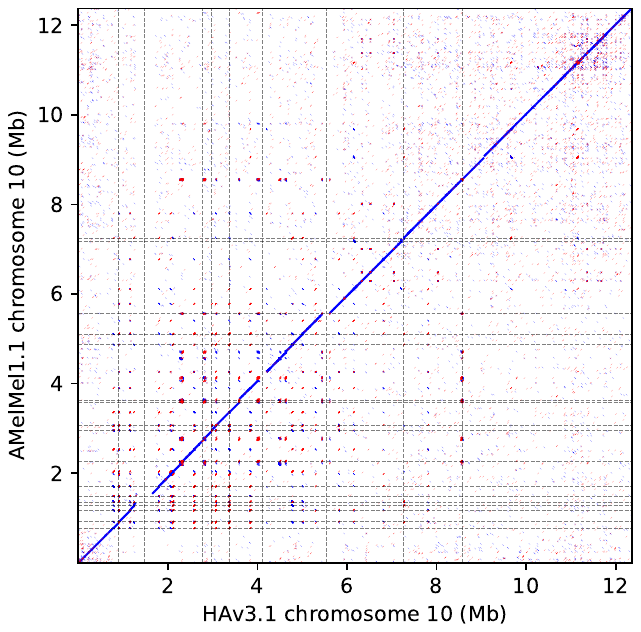
 **Chromosomes 9 and 10.** Dotplot representation of the alignment of HAv3.1 (x-axis) and AMelMel1.1 (y-axis). Blue: alignments in the same direction, red: alignments in inverted directions. Vertical and horizontal dotted lines represent contig boundaries on the two assemblies.
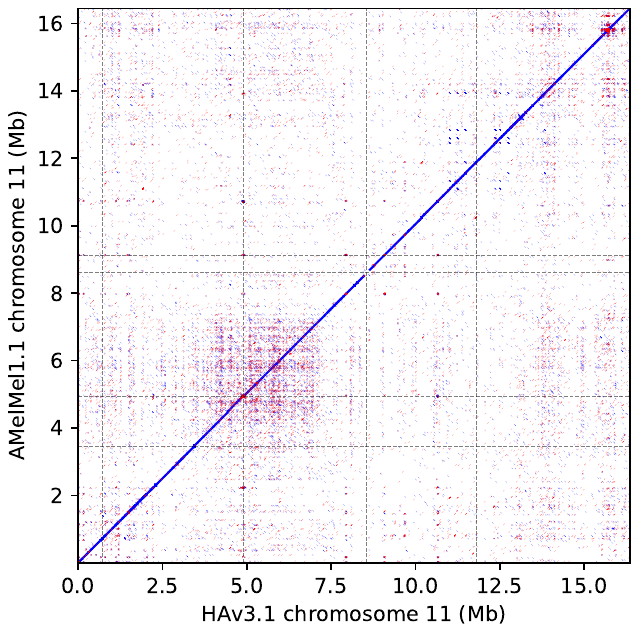

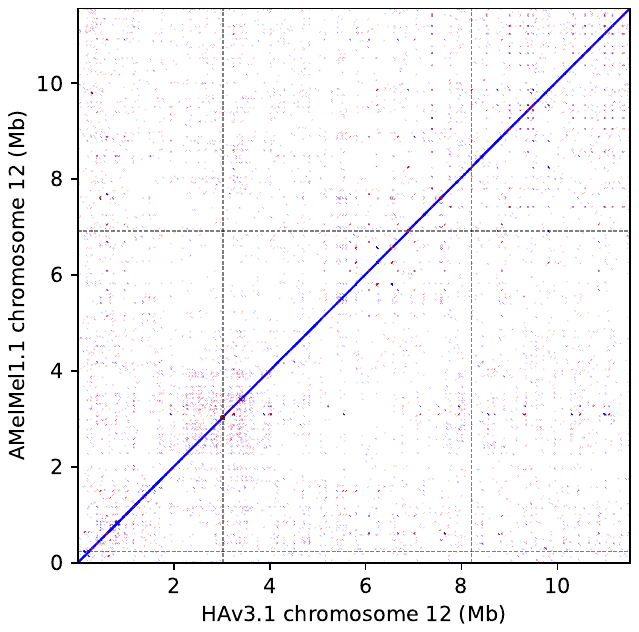
 **Chromosomes 11 and 12.** Dotplot representation of the alignment of HAv3.1 (x-axis) and AMelMel1.1 (y-axis). Blue: alignments in the same direction, red: alignments in inverted directions. Vertical and horizontal dotted lines represent contig boundaries on the two assemblies.
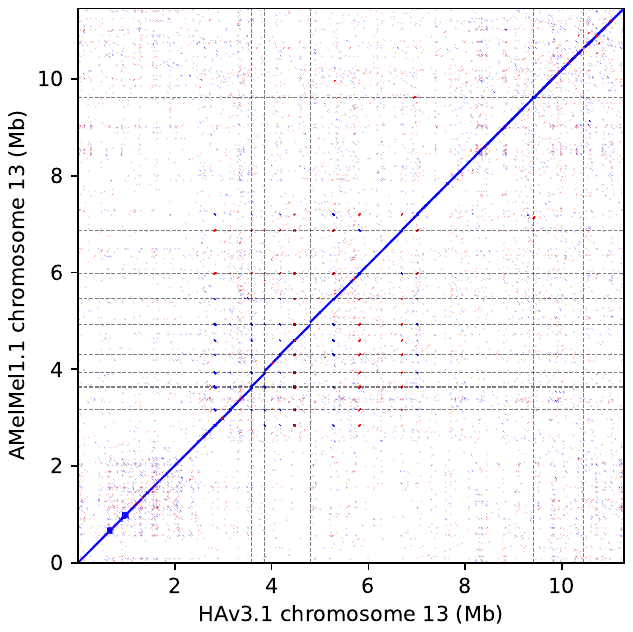

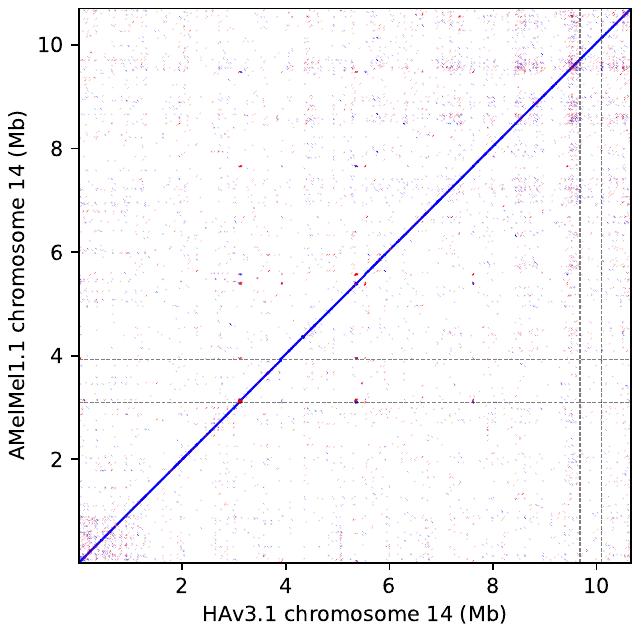
 **Chromosomes 13 and 14.** Dotplot representation of the alignment of HAv3.1 (x-axis) and AMelMel1.1 (y-axis). Blue: alignments in the same direction, red: alignments in inverted directions. Vertical and horizontal dotted lines represent contig boundaries on the two assemblies.
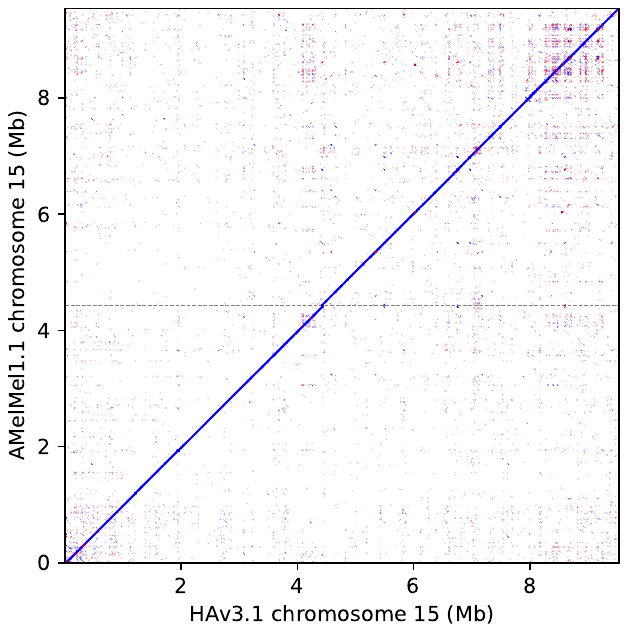

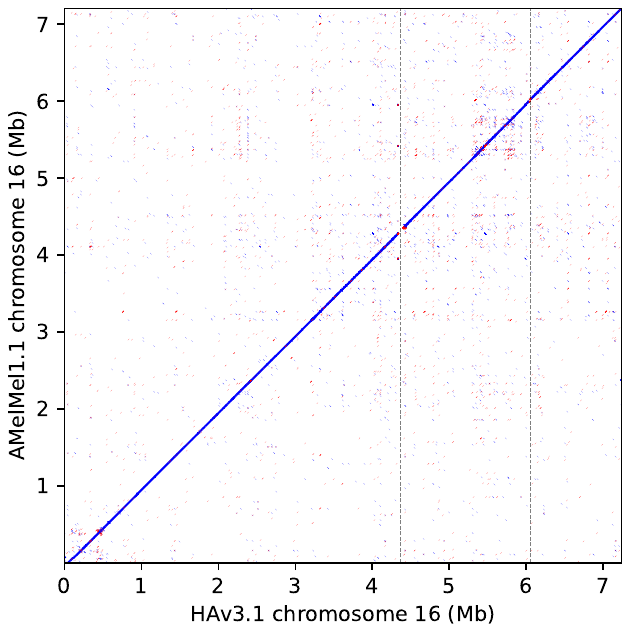
 **Chromosomes 15 and 16.** Dotplot representation of the alignment of HAv3.1 (x-axis) and AMelMel1.1 (y-axis). Blue: alignments in the same direction, red: alignments in inverted directions. Vertical and horizontal dotted lines represent contig boundaries on the two assemblies.
