## Supplementary material for "The black honey bee genome: insights on specific structural elements and a first step towards pan-genomes": Alignment of AMelMel to Amel4.5

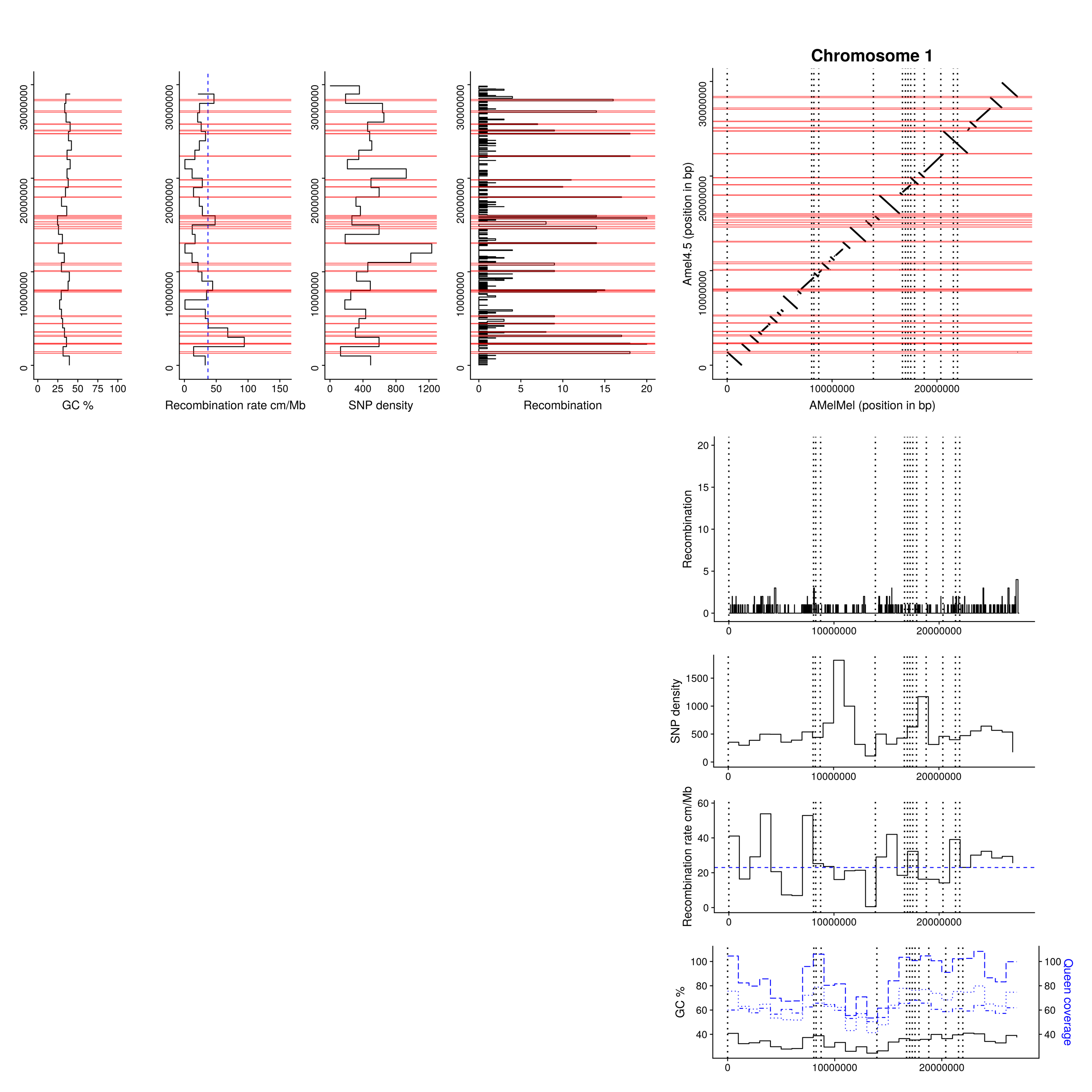

**Comparison of Amel4.5 and AMelMel assemblies: chromosome 1.** Abscissa: AMelMel, ordinate: Amel4.5. AMelMel contig borders are represented by vertical dotted lines. Additionally, for both assemblies, the position and number of recombination events detected along the chromosome are represented in each interval flanked by informative markers in the meiosis analysed. Average SNP density, recombination rate and GC% are given for 1Mb windows. Red zones represent recombination ‘hotspots’ regions where number of recombination events between two informative SNPs is higher than five events. Sequencing depth in 1 Mb windows for each of the 3 colonies analyzed to reconstruct the genetic map are in blue.

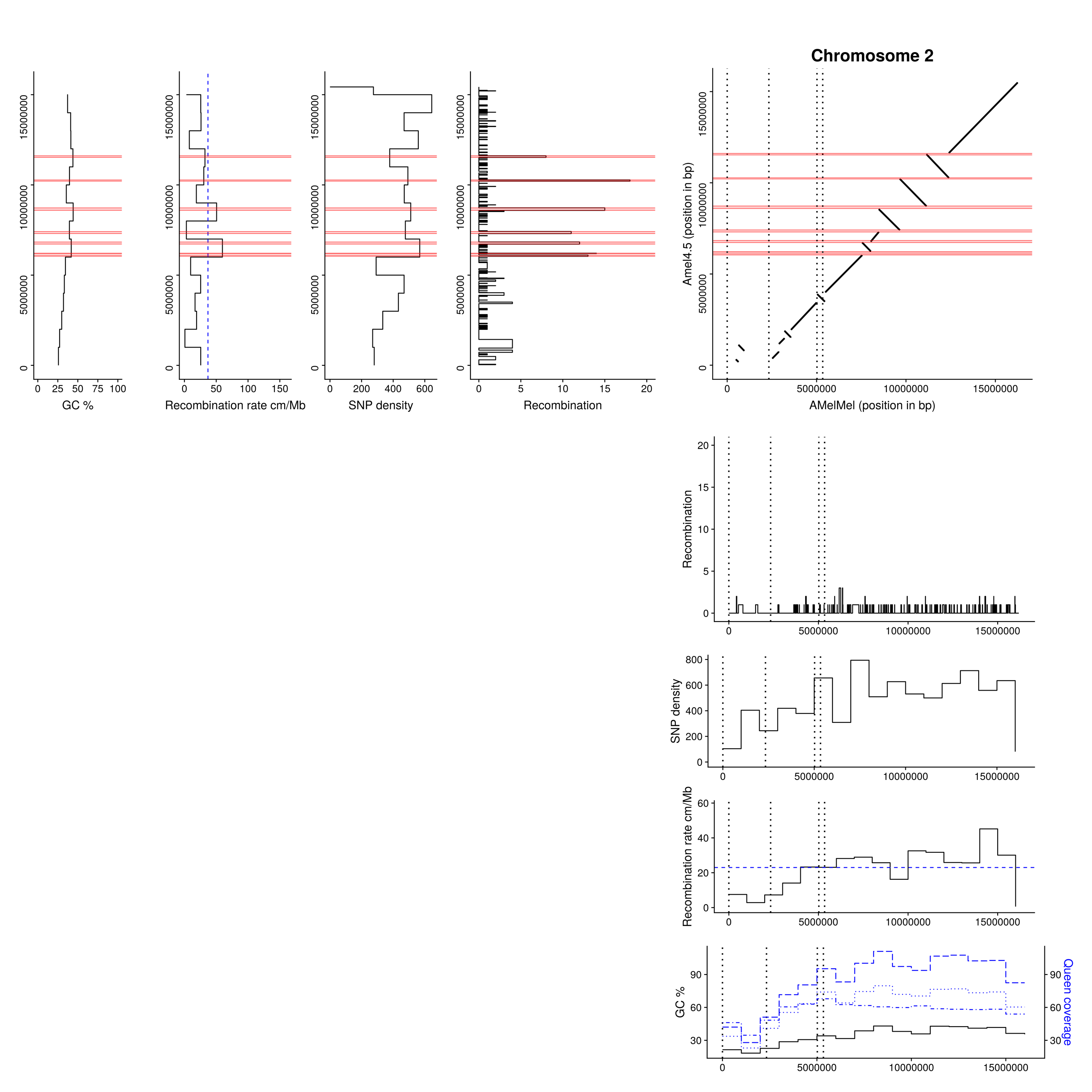

**Comparison of Amel4.5 and AMelMel assemblies: chromosome 2.** Abscissa: AMelMel, ordinate: Amel4.5. AMelMel contig borders are represented by vertical dotted lines. Additionally, for both assemblies, the position and number of recombination events detected along the chromosome are represented in each interval flanked by informative markers in the meiosis analysed. Average SNP density, recombination rate and GC% are given for 1Mb windows. Red zones represent recombination ‘hotspots’ regions where number of recombination events between two informative SNPs is higher than five events. Sequencing depth in 1 Mb windows for each of the 3 colonies analyzed to reconstruct the genetic map are in blue.

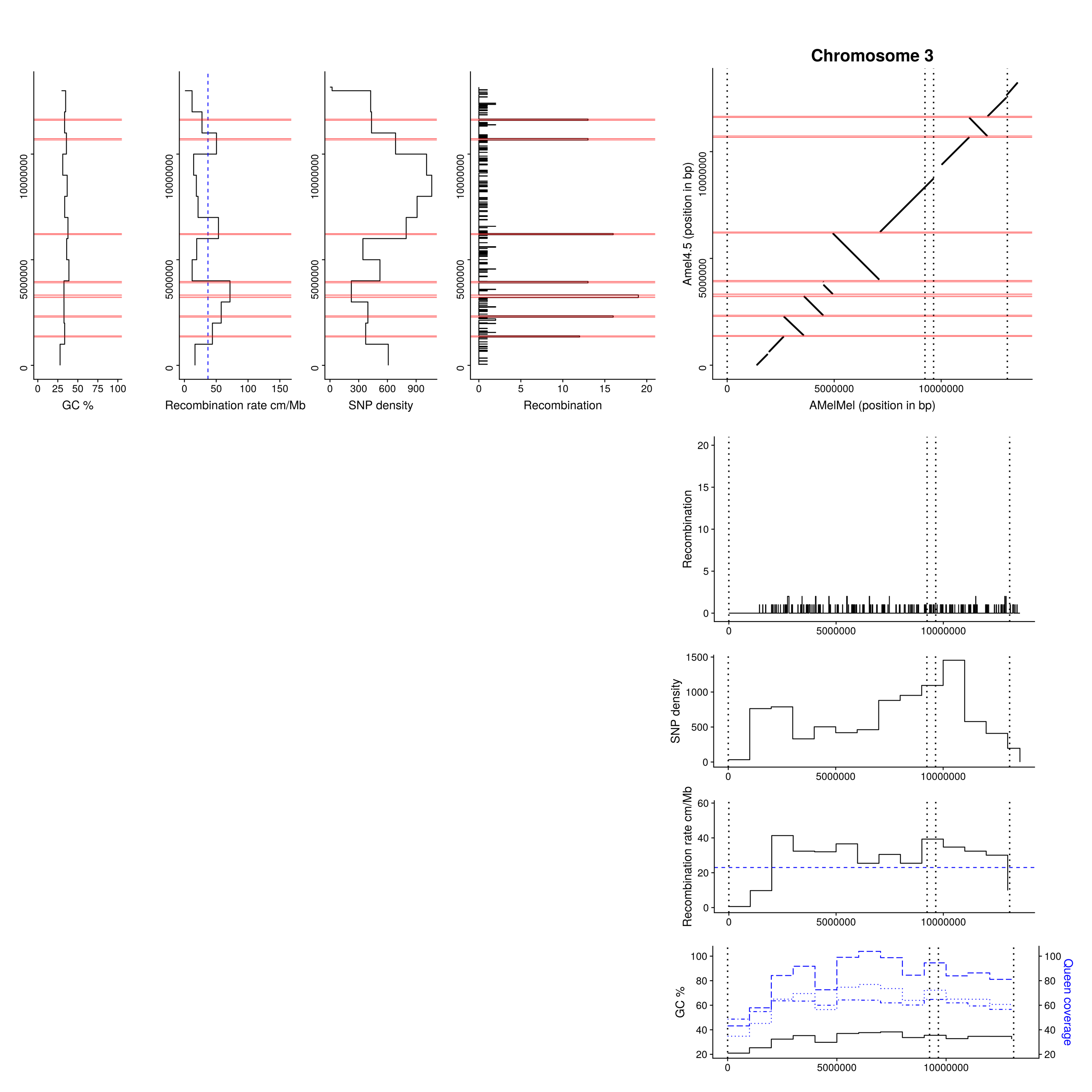

**Comparison of Amel4.5 and AMelMel assemblies: chromosome 3.** Abscissa: AMelMel, ordinate: Amel4.5. AMelMel contig borders are represented by vertical dotted lines. Additionally, for both assemblies, the position and number of recombination events detected along the chromosome are represented in each interval flanked by informative markers in the meiosis analysed. Average SNP density, recombination rate and GC% are given for 1Mb windows. Red zones represent recombination ‘hotspots’ regions where number of recombination events between two informative SNPs is higher than five events. Sequencing depth in 1 Mb windows for each of the 3 colonies analyzed to reconstruct the genetic map are in blue.

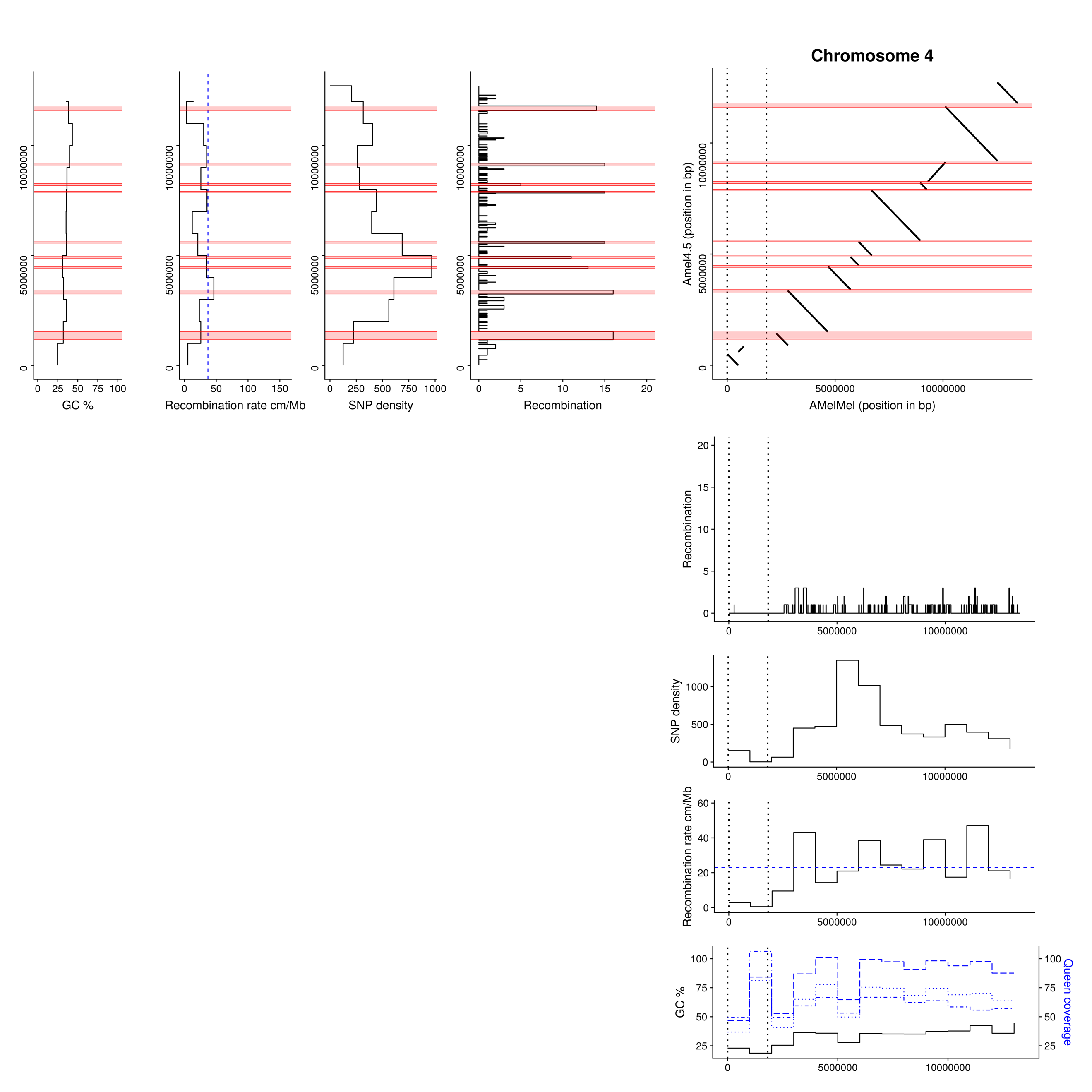

**Comparison of Amel4.5 and AMelMel assemblies: chromosome 4.** Abscissa: AMelMel, ordinate: Amel4.5. AMelMel contig borders are represented by vertical dotted lines. Additionally, for both assemblies, the position and number of recombination events detected along the chromosome are represented in each interval flanked by informative markers in the meiosis analysed. Average SNP density, recombination rate and GC% are given for 1Mb windows. Red zones represent recombination ‘hotspots’ regions where number of recombination events between two informative SNPs is higher than five events. Sequencing depth in 1 Mb windows for each of the 3 colonies analyzed to reconstruct the genetic map are in blue.

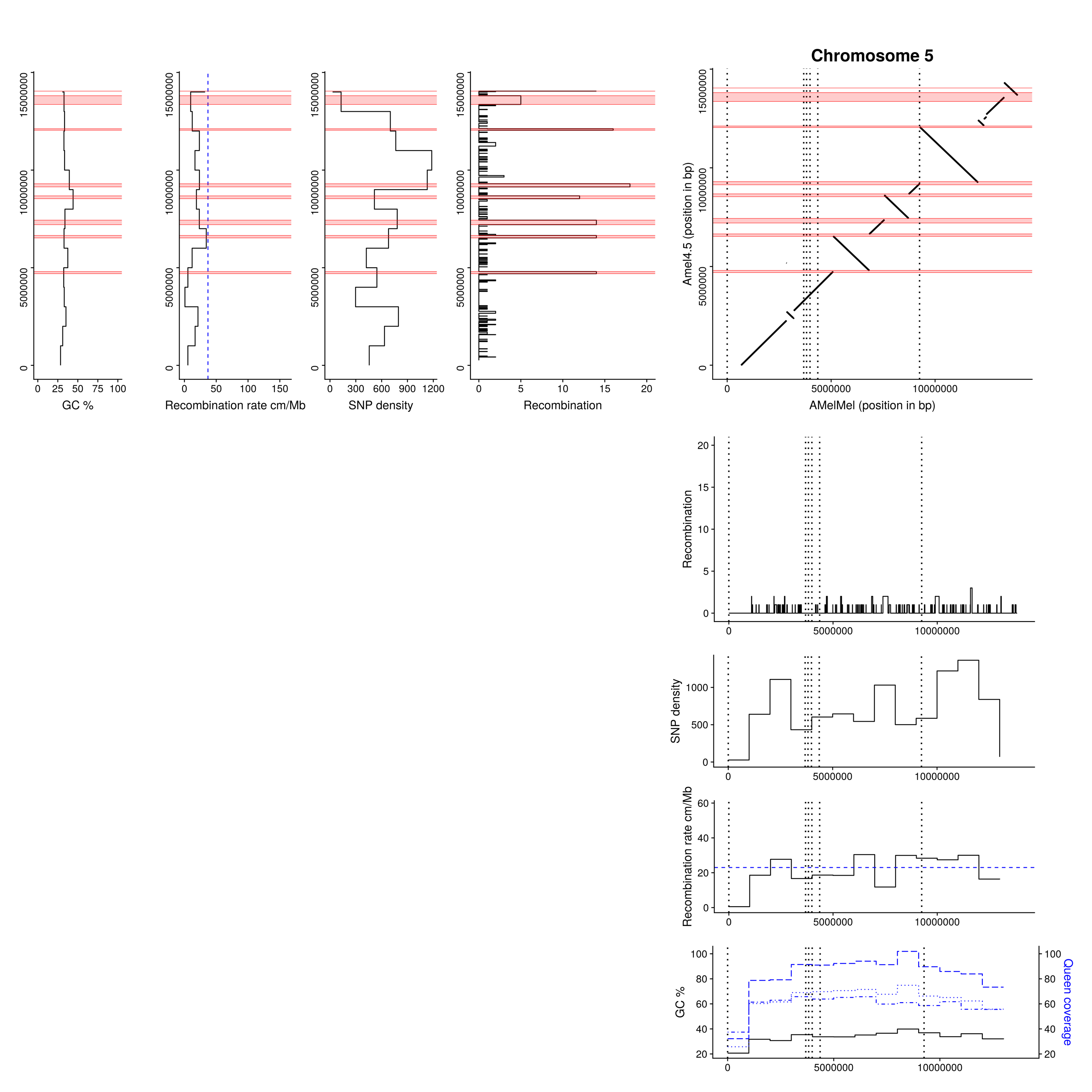

**Comparison of Amel4.5 and AMelMel assemblies: chromosome 5.** Abscissa: AMelMel, ordinate: Amel4.5. AMelMel contig borders are represented by vertical dotted lines. Additionally, for both assemblies, the position and number of recombination events detected along the chromosome are represented in each interval flanked by informative markers in the meiosis analysed. Average SNP density, recombination rate and GC% are given for 1Mb windows. Red zones represent recombination ‘hotspots’ regions where number of recombination events between two informative SNPs is higher than five events. Sequencing depth in 1 Mb windows for each of the 3 colonies analyzed to reconstruct the genetic map are in blue.

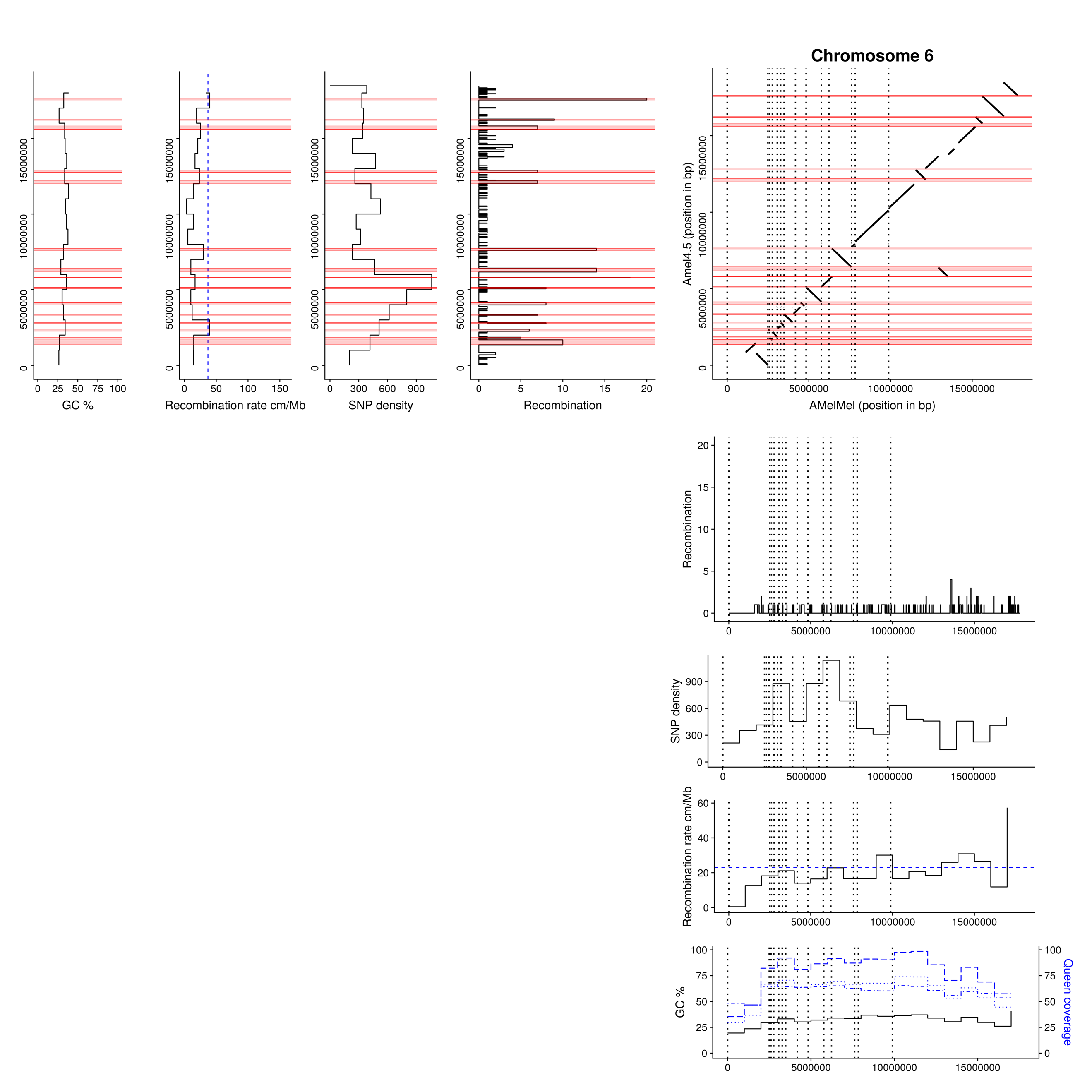

**Comparison of Amel4.5 and AMelMel assemblies: chromosome 6.** Abscissa: AMelMel, ordinate: Amel4.5. AMelMel contig borders are represented by vertical dotted lines. Additionally, for both assemblies, the position and number of recombination events detected along the chromosome are represented in each interval flanked by informative markers in the meiosis analysed. Average SNP density, recombination rate and GC% are given for 1Mb windows. Red zones represent recombination ‘hotspots’ regions where number of recombination events between two informative SNPs is higher than five events. Sequencing depth in 1 Mb windows for each of the 3 colonies analyzed to reconstruct the genetic map are in blue.

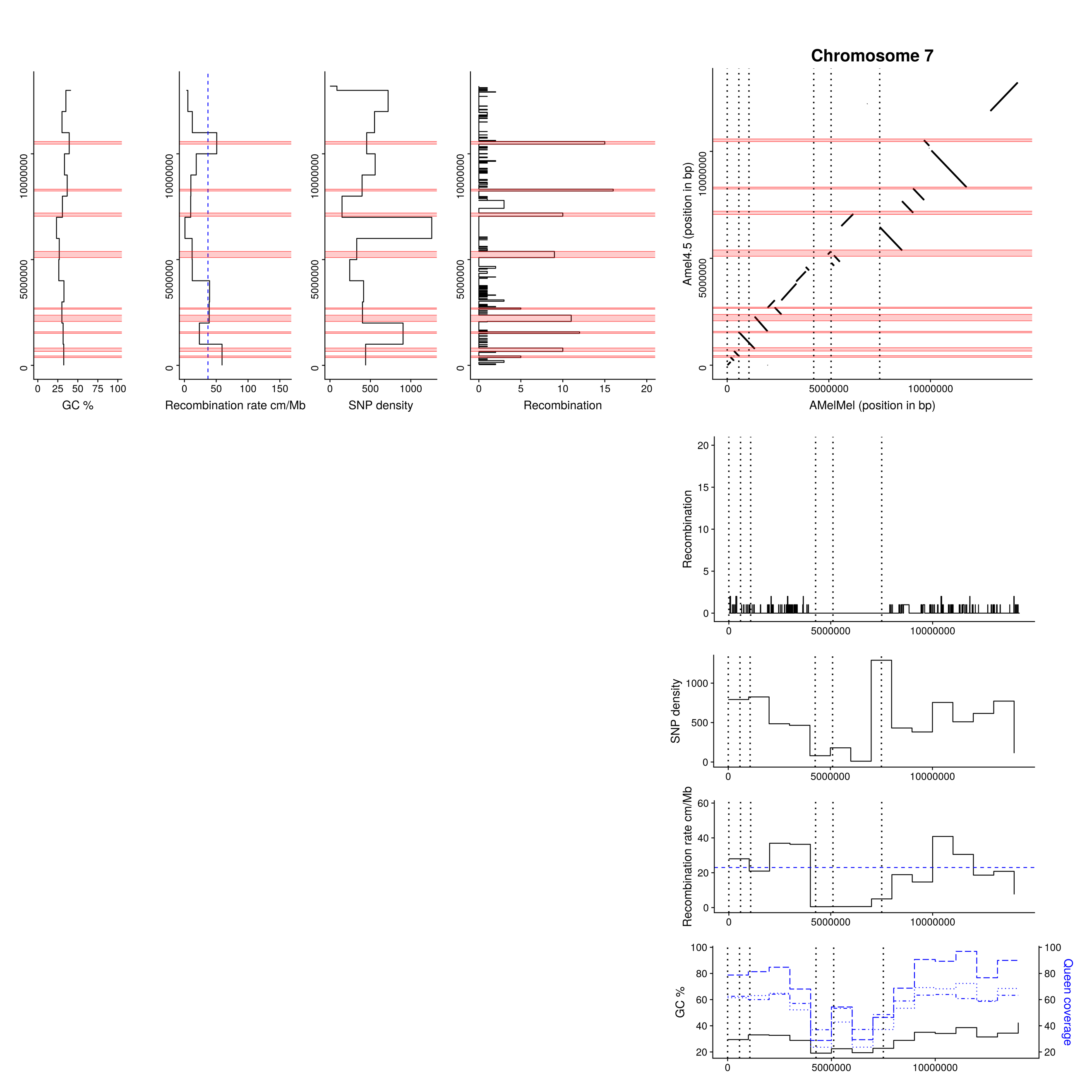

**Comparison of Amel4.5 and AMelMel assemblies: chromosome 7.** Abscissa: AMelMel, ordinate: Amel4.5. AMelMel contig borders are represented by vertical dotted lines. Additionally, for both assemblies, the position and number of recombination events detected along the chromosome are represented in each interval flanked by informative markers in the meiosis analysed. Average SNP density, recombination rate and GC% are given for 1Mb windows. Red zones represent recombination ‘hotspots’ regions where number of recombination events between two informative SNPs is higher than five events. Sequencing depth in 1 Mb windows for each of the 3 colonies analyzed to reconstruct the genetic map are in blue.

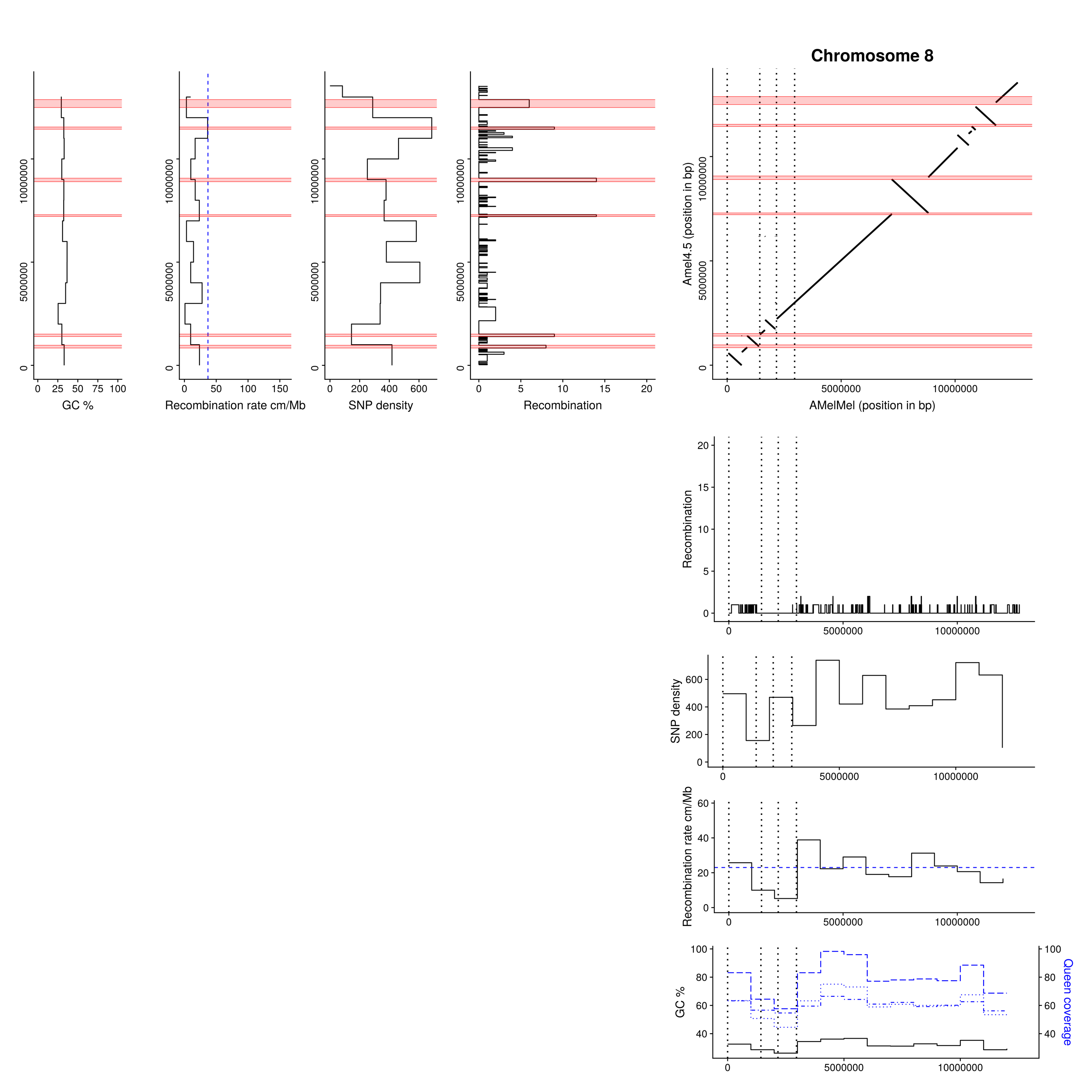

**Comparison of Amel4.5 and AMelMel assemblies: chromosome 8.** Abscissa: AMelMel, ordinate: Amel4.5. AMelMel contig borders are represented by vertical dotted lines. Additionally, for both assemblies, the position and number of recombination events detected along the chromosome are represented in each interval flanked by informative markers in the meiosis analysed. Average SNP density, recombination rate and GC% are given for 1Mb windows. Red zones represent recombination ‘hotspots’ regions where number of recombination events between two informative SNPs is higher than five events. Sequencing depth in 1 Mb windows for each of the 3 colonies analyzed to reconstruct the genetic map are in blue.

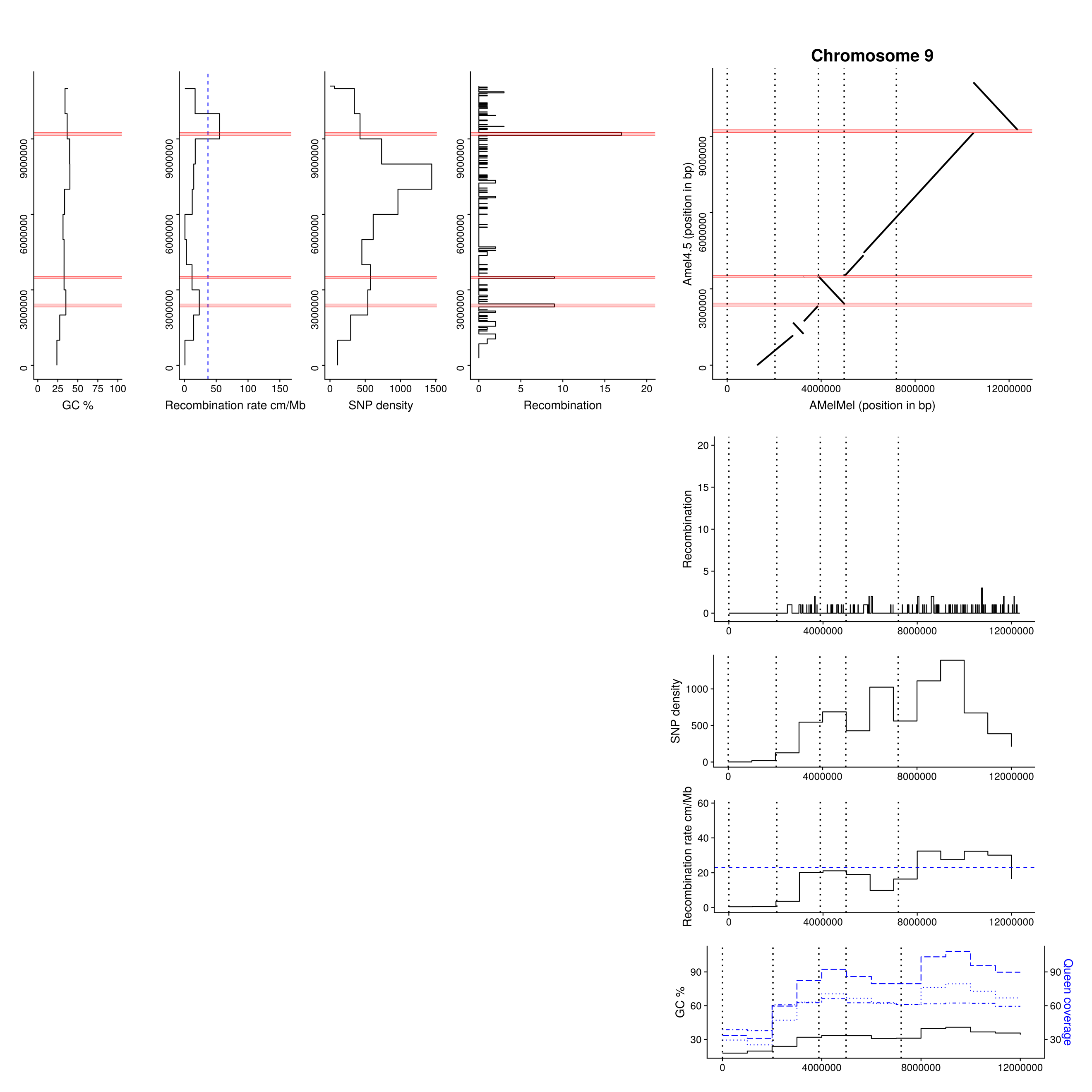

**Comparison of Amel4.5 and AMelMel assemblies: chromosome 9.** Abscissa: AMelMel, ordinate: Amel4.5. AMelMel contig borders are represented by vertical dotted lines. Additionally, for both assemblies, the position and number of recombination events detected along the chromosome are represented in each interval flanked by informative markers in the meiosis analysed. Average SNP density, recombination rate and GC% are given for 1Mb windows. Red zones represent recombination ‘hotspots’ regions where number of recombination events between two informative SNPs is higher than five events. Sequencing depth in 1 Mb windows for each of the 3 colonies analyzed to reconstruct the genetic map are in blue.

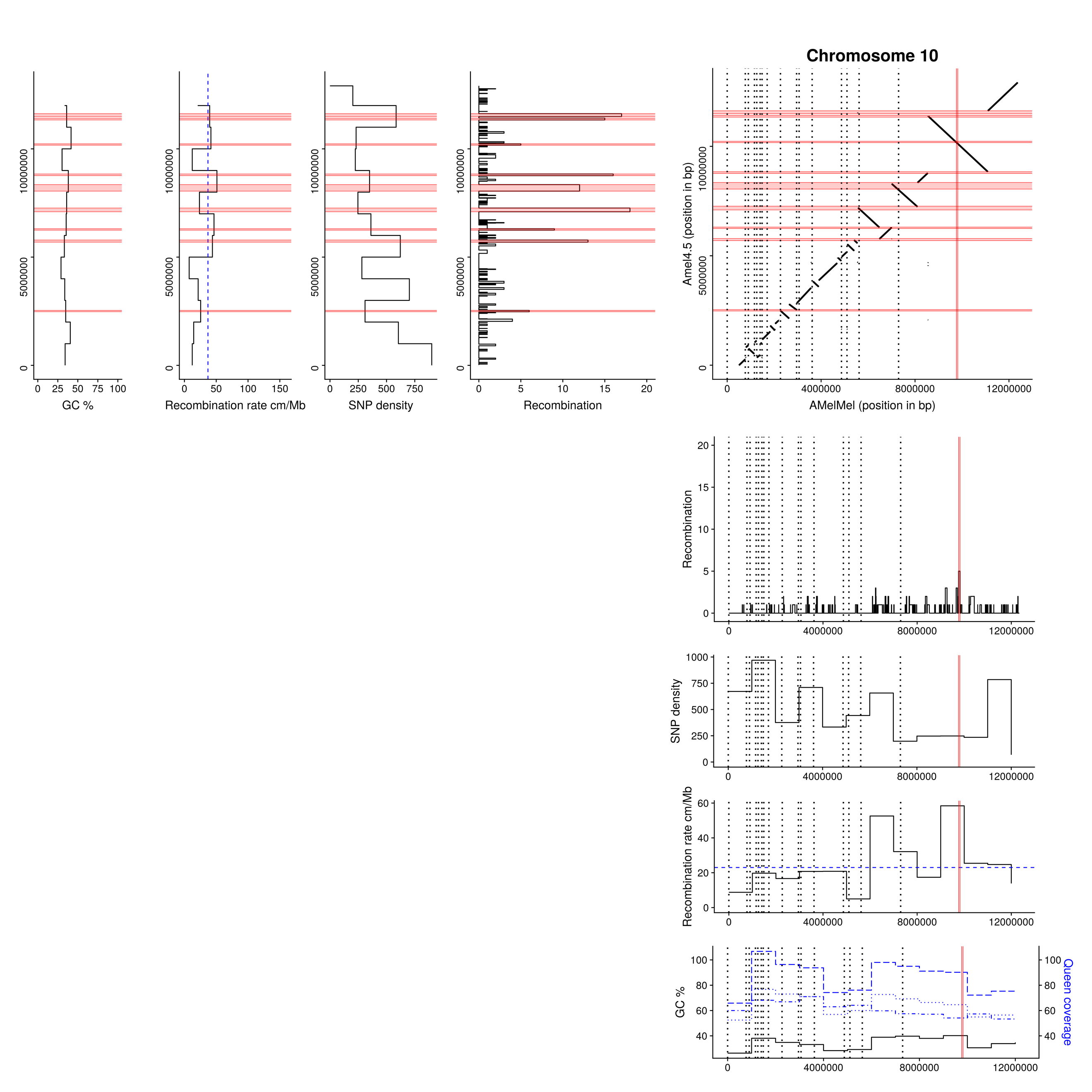

**Comparison of Amel4.5 and AMelMel assemblies: chromosome 10.** Abscissa: AMelMel, ordinate: Amel4.5. AMelMel contig borders are represented by vertical dotted lines. Additionally, for both assemblies, the position and number of recombination events detected along the chromosome are represented in each interval flanked by informative markers in the meiosis analysed. Average SNP density, recombination rate and GC% are given for 1Mb windows. Red zones represent recombination ‘hotspots’ regions where number of recombination events between two informative SNPs is higher than five events. Sequencing depth in 1 Mb windows for each of the 3 colonies analyzed to reconstruct the genetic map are in blue.

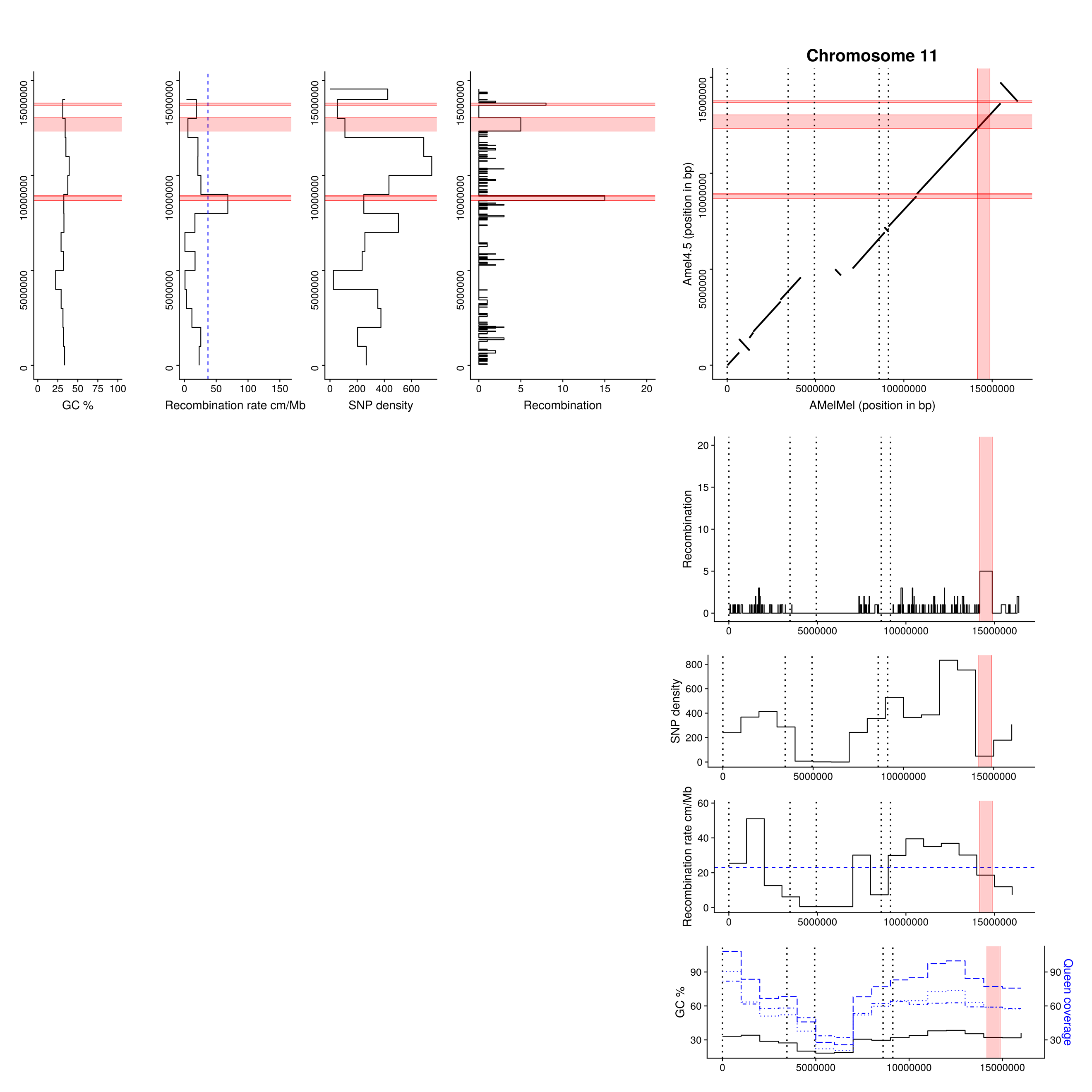

**Comparison of Amel4.5 and AMelMel assemblies: chromosome 11.** Abscissa: AMelMel, ordinate: Amel4.5. AMelMel contig borders are represented by vertical dotted lines. Additionally, for both assemblies, the position and number of recombination events detected along the chromosome are represented in each interval flanked by informative markers in the meiosis analysed. Average SNP density, recombination rate and GC% are given for 1Mb windows. Red zones represent recombination ‘hotspots’ regions where number of recombination events between two informative SNPs is higher than five events. Sequencing depth in 1 Mb windows for each of the 3 colonies analyzed to reconstruct the genetic map are in blue.

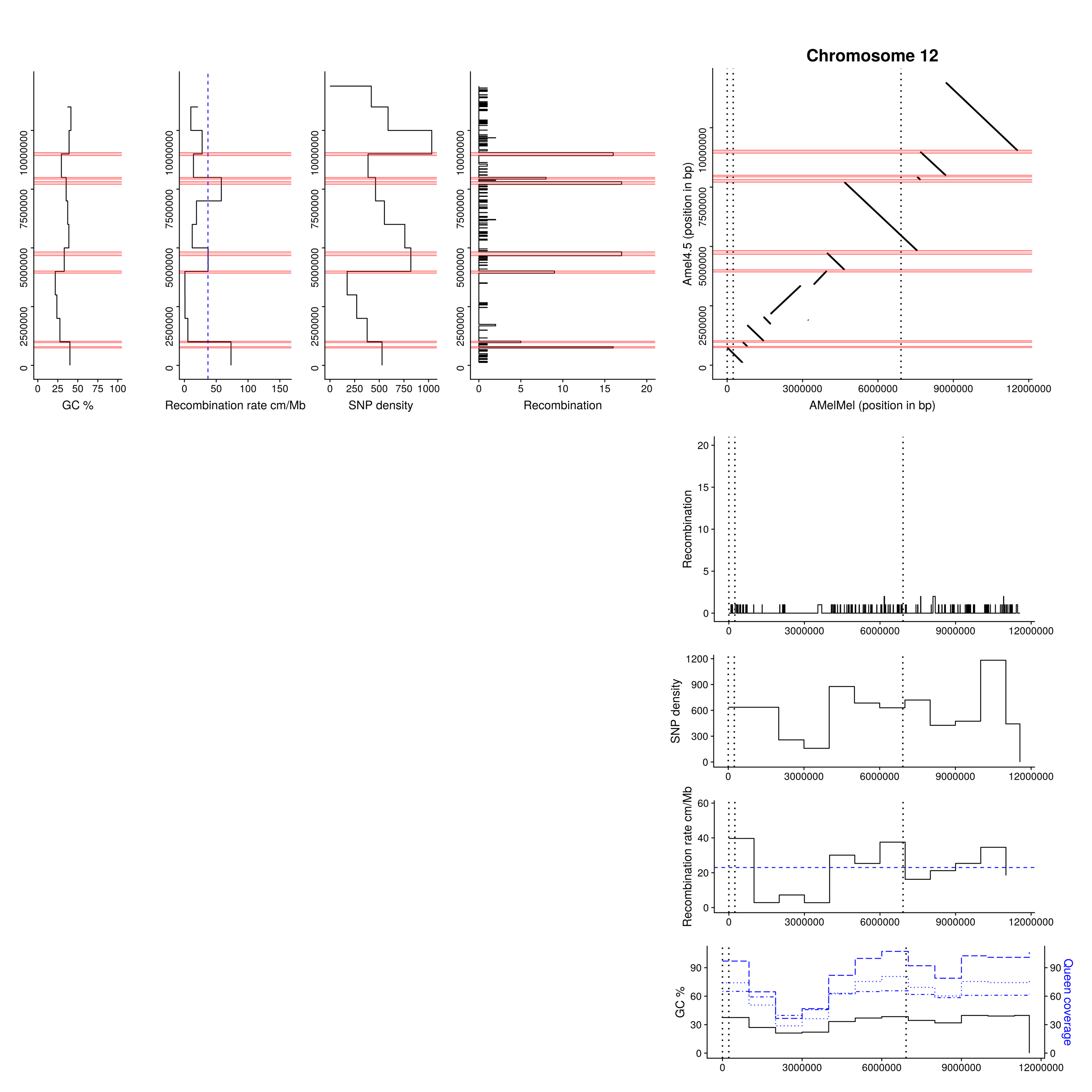

**Comparison of Amel4.5 and AMelMel assemblies: chromosome 12.** Abscissa: AMelMel, ordinate: Amel4.5. AMelMel contig borders are represented by vertical dotted lines. Additionally, for both assemblies, the position and number of recombination events detected along the chromosome are represented in each interval flanked by informative markers in the meiosis analysed. Average SNP density, recombination rate and GC% are given for 1Mb windows. Red zones represent recombination ‘hotspots’ regions where number of recombination events between two informative SNPs is higher than five events. Sequencing depth in 1 Mb windows for each of the 3 colonies analyzed to reconstruct the genetic map are in blue.

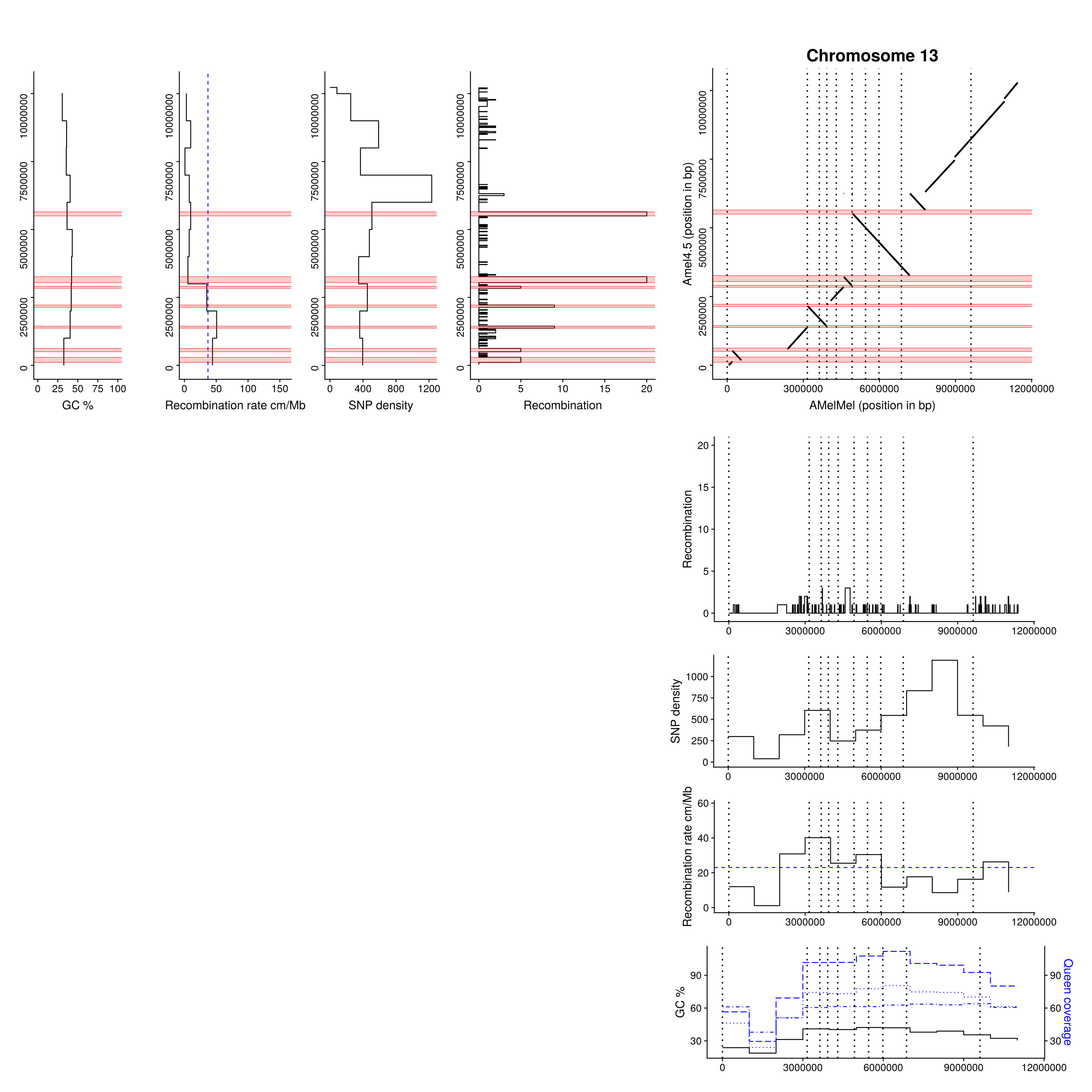

**Comparison of Amel4.5 and AMelMel assemblies: chromosome 13.** Abscissa: AMelMel, ordinate: Amel4.5. AMelMel contig borders are represented by vertical dotted lines. Additionally, for both assemblies, the position and number of recombination events detected along the chromosome are represented in each interval flanked by informative markers in the meiosis analysed. Average SNP density, recombination rate and GC% are given for 1Mb windows. Red zones represent recombination ‘hotspots’ regions where number of recombination events between two informative SNPs is higher than five events. Sequencing depth in 1 Mb windows for each of the 3 colonies analyzed to reconstruct the genetic map are in blue.

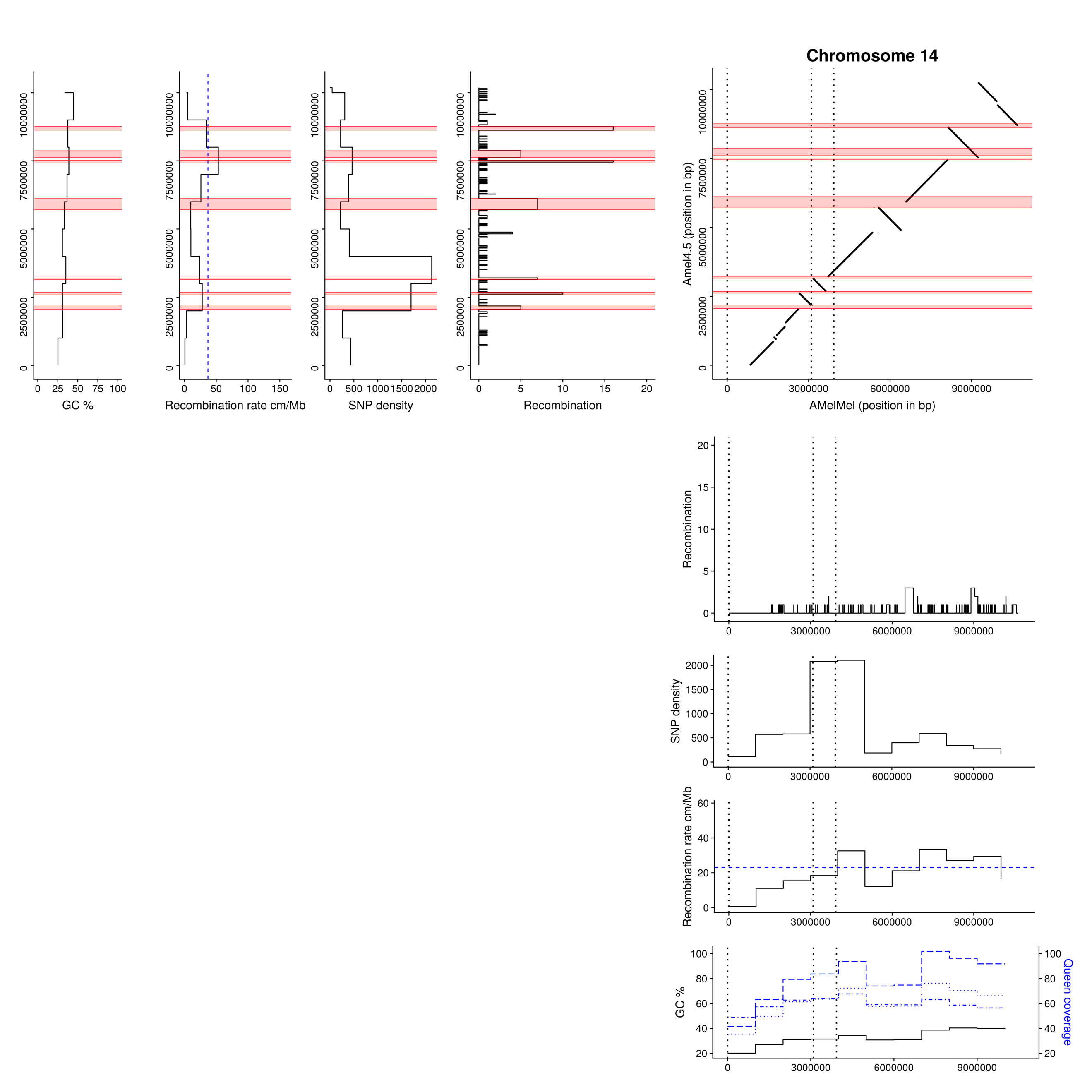

**Comparison of Amel4.5 and AMelMel assemblies: chromosome 14.** Abscissa: AMelMel, ordinate: Amel4.5. AMelMel contig borders are represented by vertical dotted lines. Additionally, for both assemblies, the position and number of recombination events detected along the chromosome are represented in each interval flanked by informative markers in the meiosis analysed. Average SNP density, recombination rate and GC% are given for 1Mb windows. Red zones represent recombination ‘hotspots’ regions where number of recombination events between two informative SNPs is higher than five events. Sequencing depth in 1 Mb windows for each of the 3 colonies analyzed to reconstruct the genetic map are in blue.

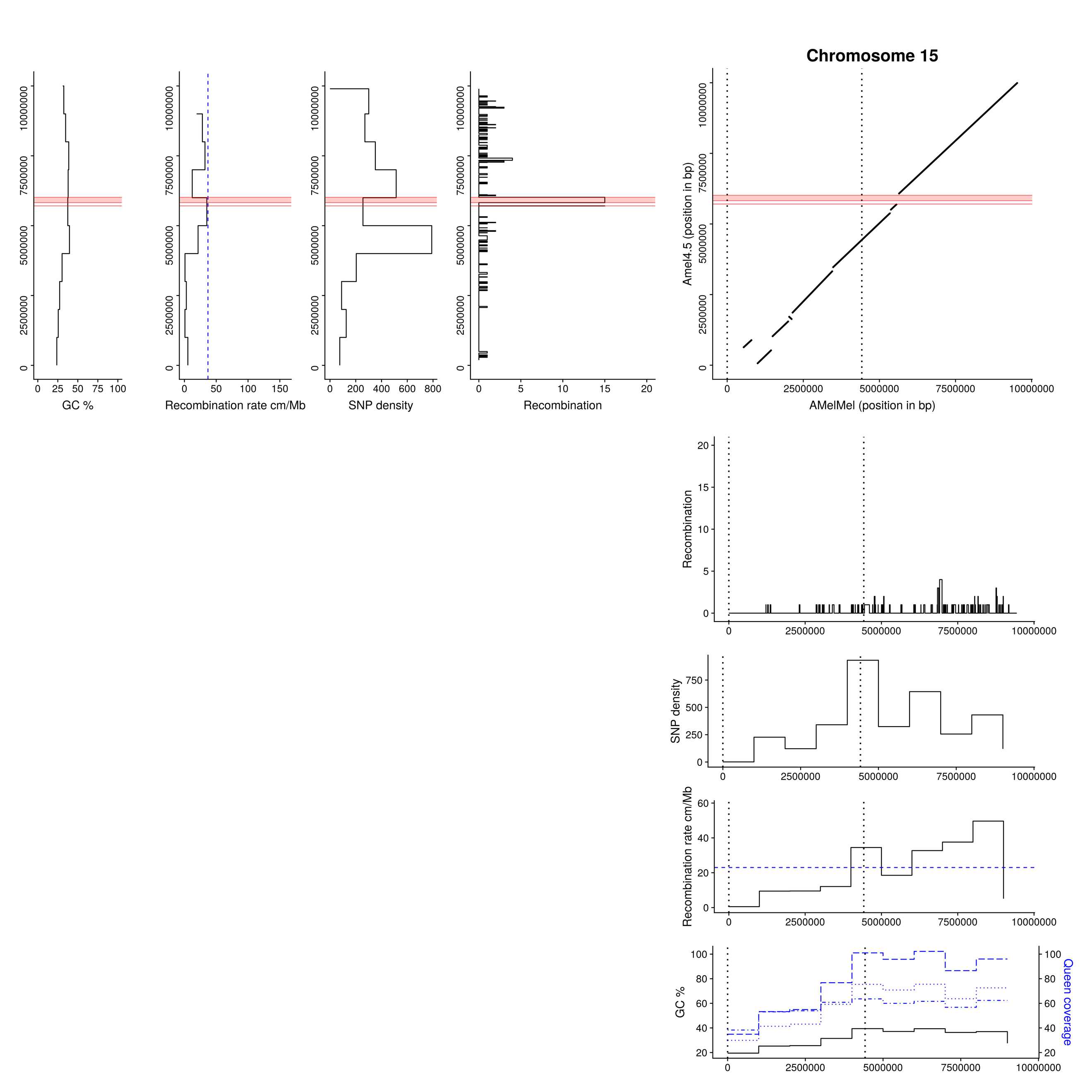

**Comparison of Amel4.5 and AMelMel assemblies: chromosome 15.** Abscissa: AMelMel, ordinate: Amel4.5. AMelMel contig borders are represented by vertical dotted lines. Additionally, for both assemblies, the position and number of recombination events detected along the chromosome are represented in each interval flanked by informative markers in the meiosis analysed. Average SNP density, recombination rate and GC% are given for 1Mb windows. Red zones represent recombination ‘hotspots’ regions where number of recombination events between two informative SNPs is higher than five events. Sequencing depth in 1 Mb windows for each of the 3 colonies analyzed to reconstruct the genetic map are in blue.

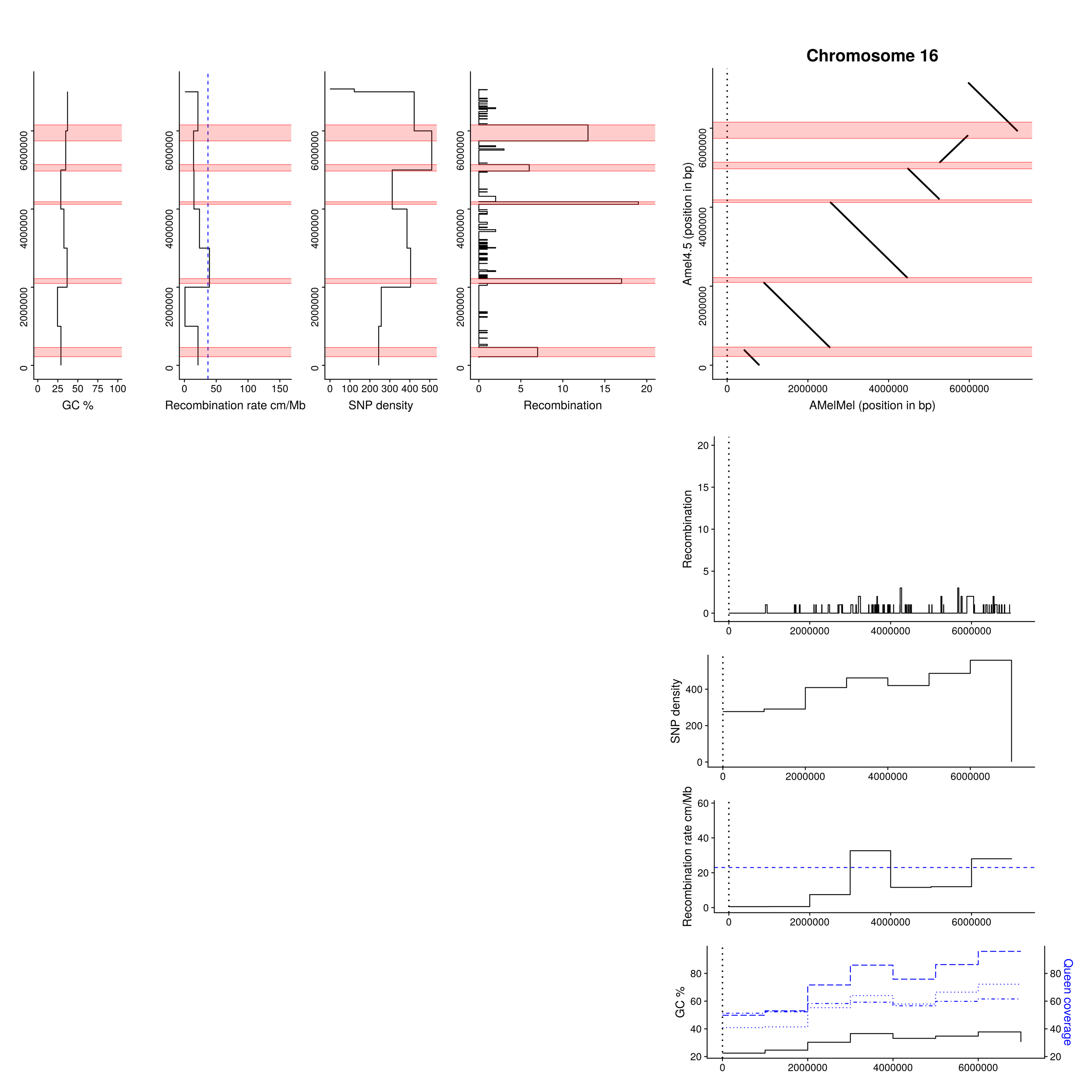

**Comparison of Amel4.5 and AMelMel assemblies: chromosome 16.** Abscissa: AMelMel, ordinate: Amel4.5. AMelMel contig borders are represented by vertical dotted lines. Additionally, for both assemblies, the position and number of recombination events detected along the chromosome are represented in each interval flanked by informative markers in the meiosis analysed. Average SNP density, recombination rate and GC% are given for 1Mb windows. Red zones represent recombination ‘hotspots’ regions where number of recombination events between two informative SNPs is higher than five events. Sequencing depth in 1 Mb windows for each of the 3 colonies analyzed to reconstruct the genetic map are in blue.
