## Supplementary material for "The black honey bee genome: insights on specific structural elements and a first step towards pan-genomes": Chromosomal inversions between AMelMel and HAv3.1

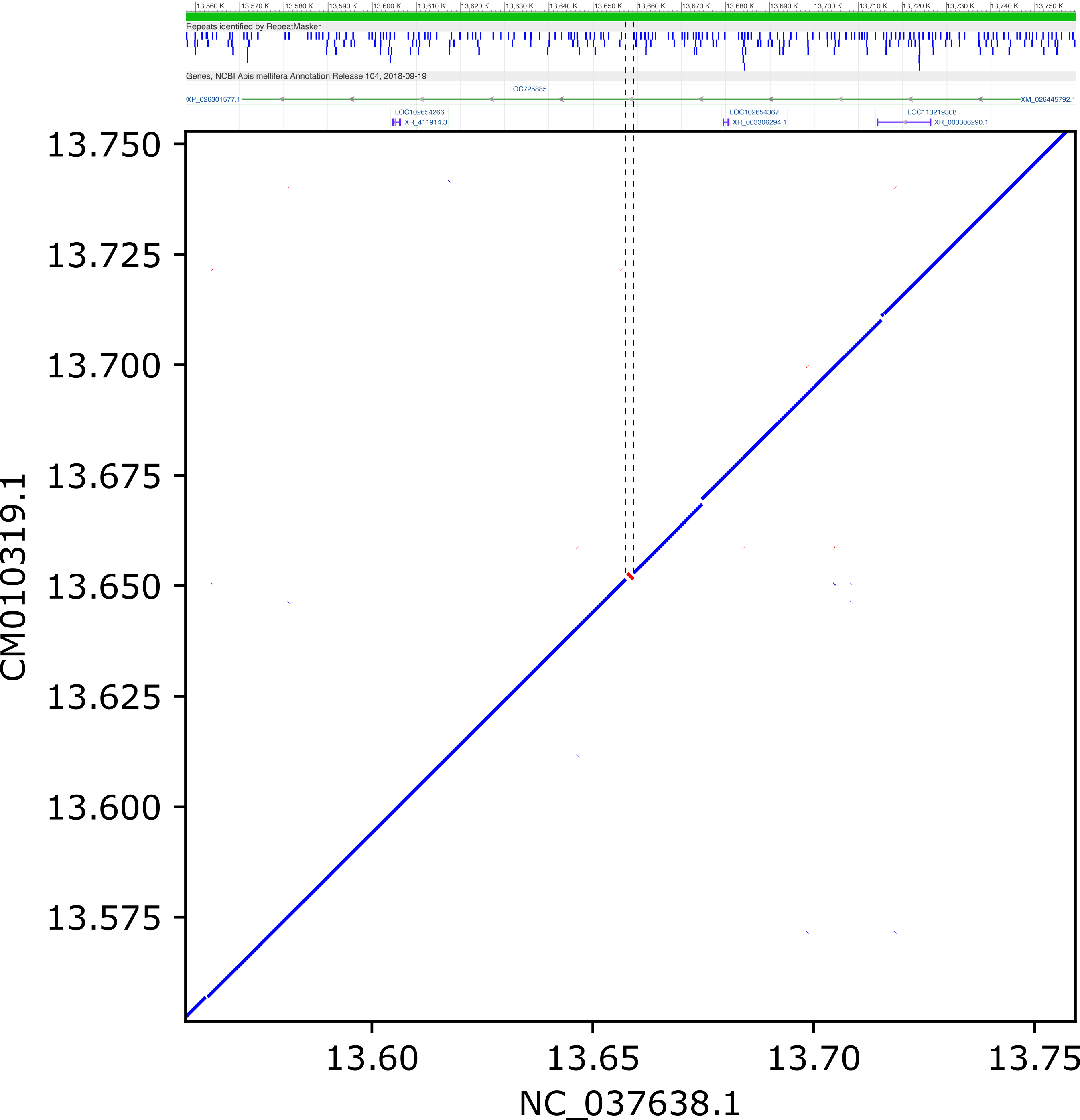

**Inversions > 1 kb between HAv3.1 and AMelMel1.1 on chromosome 1, detected by LAST and distant from contig breakpoints**

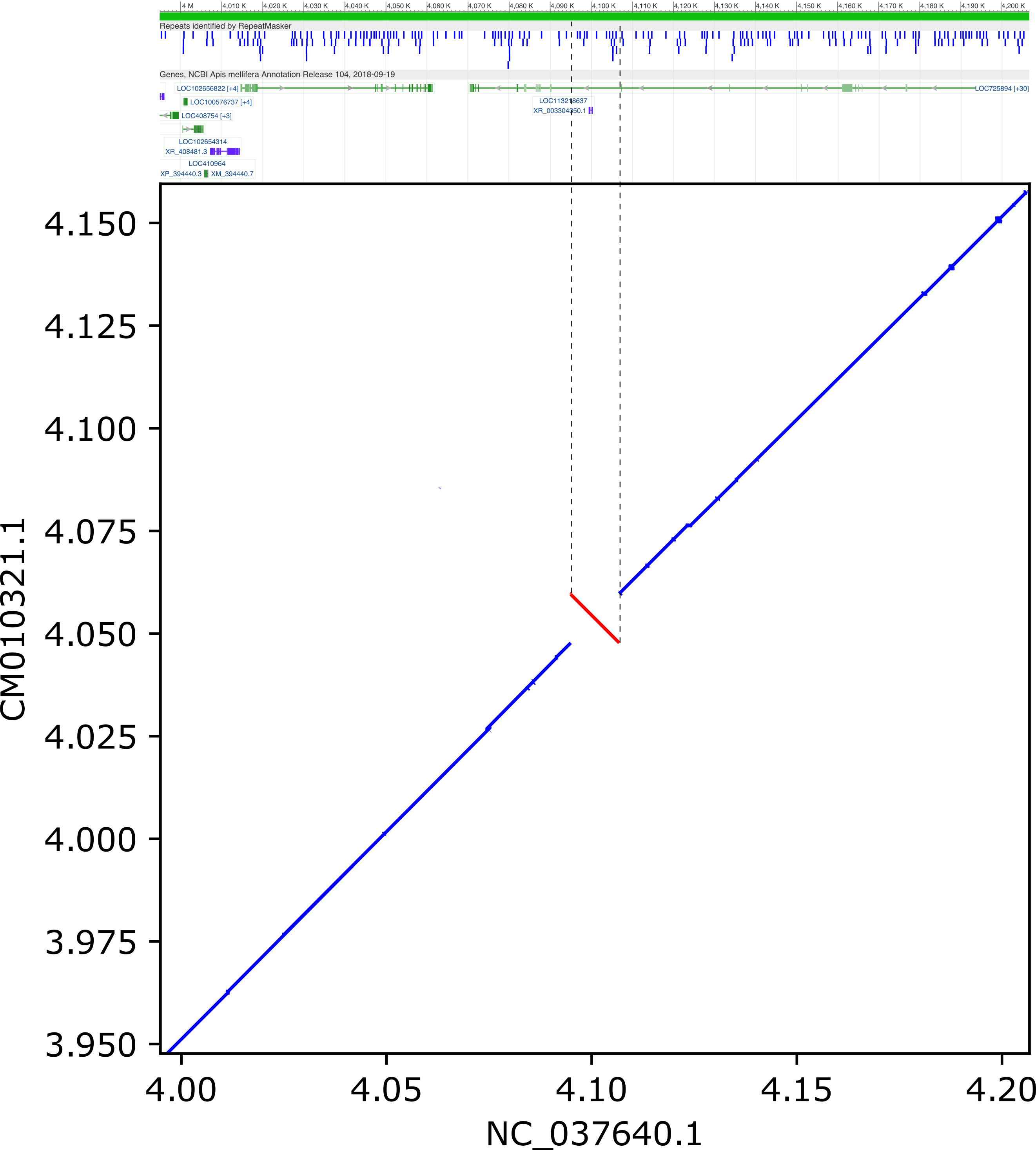

**Inversions > 1 kb between HAv3.1 and AMelMel1.1 on chromosome 3, detected by LAST and distant from contig breakpoints**

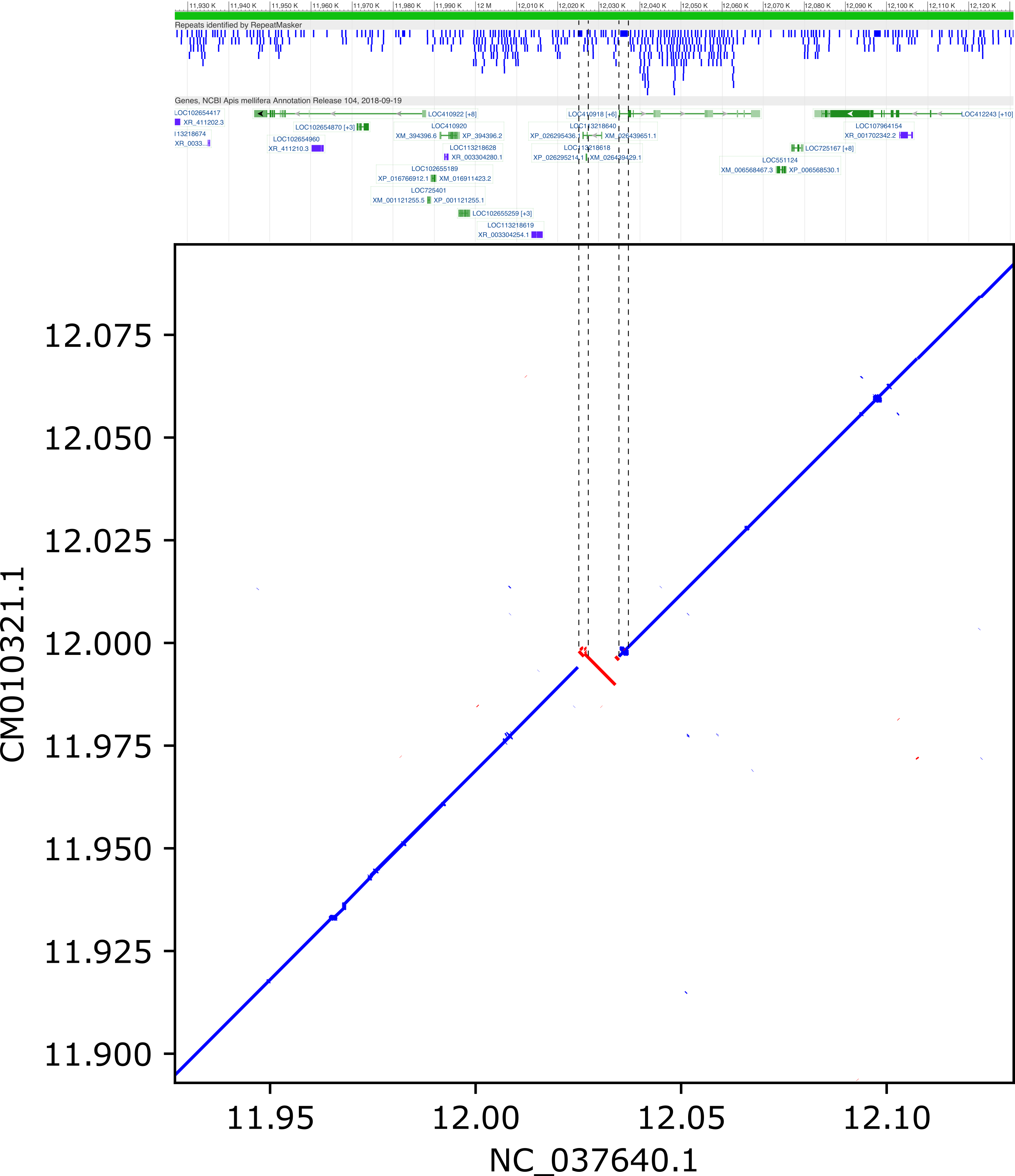

**Inversions > 1 kb between HAv3.1 and AMelMel1.1 on chromosome 3, detected by LAST and distant from contig breakpoints**

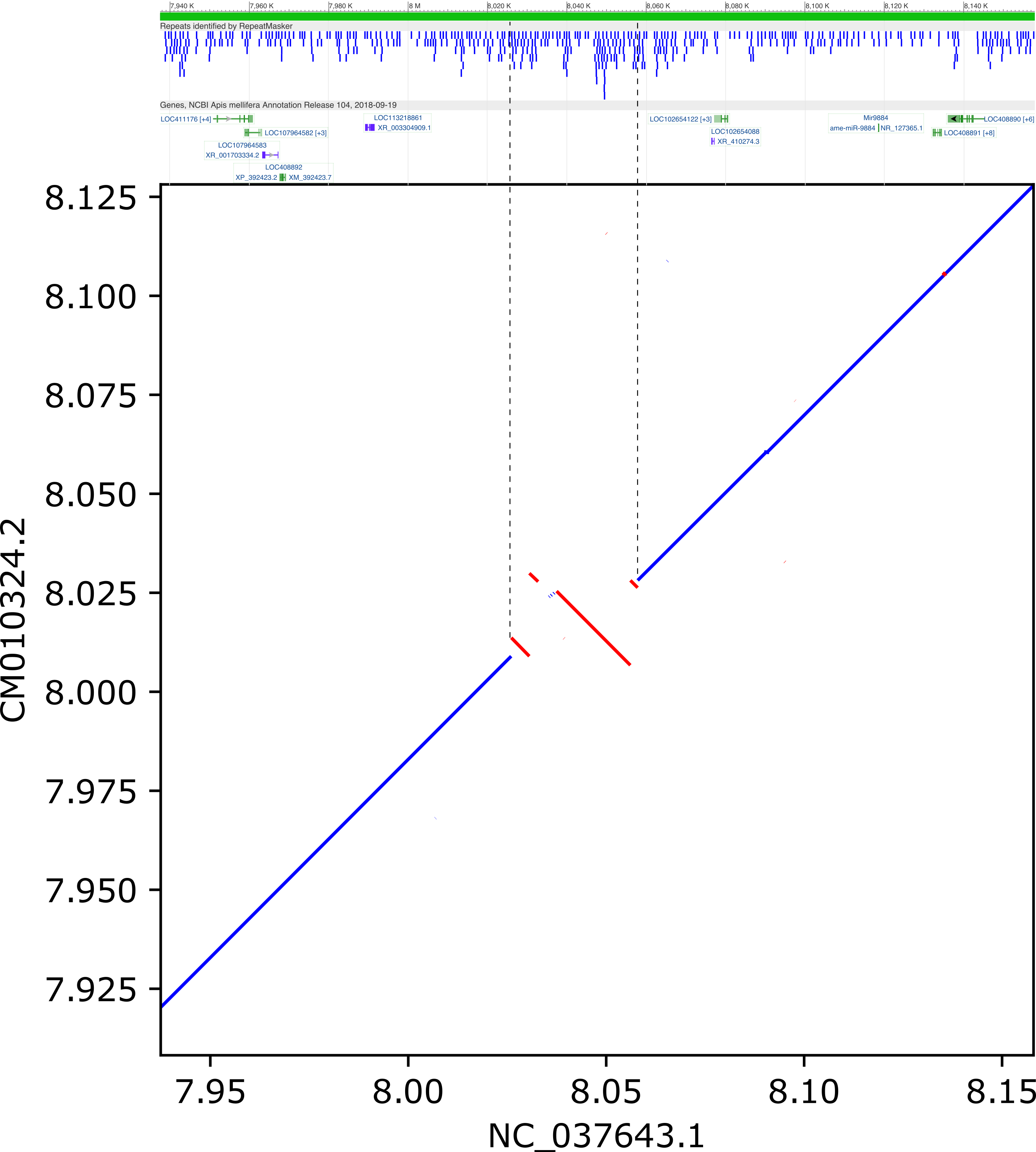

**Inversions > 1 kb between HAv3.1 and AMelMel1.1 on chromosome 6, detected by LAST and distant from contig breakpoints**

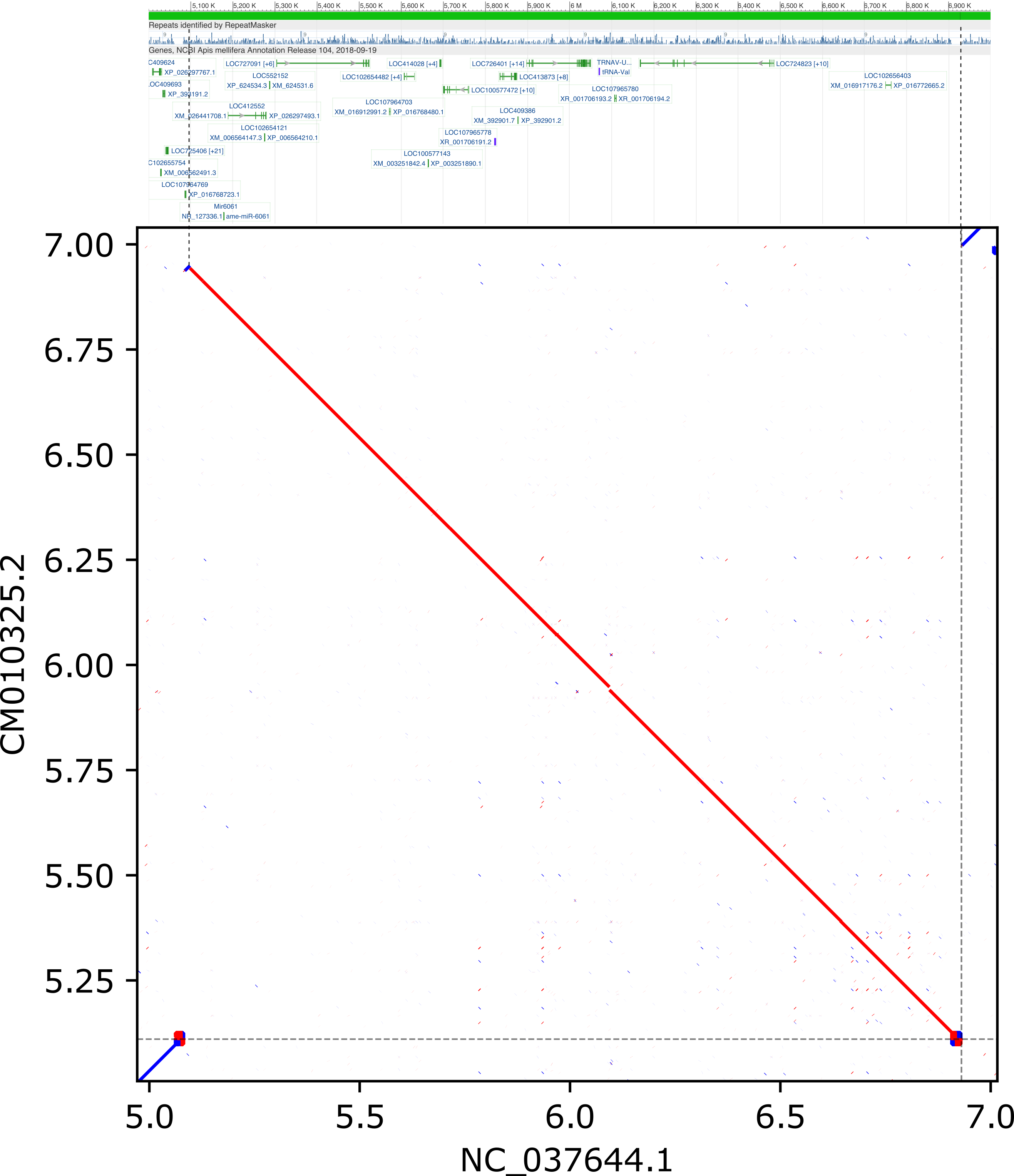

**Inversions > 1 kb between HAv3.1 and AMelMel1.1 on chromosome 7, detected by LAST and distant from contig breakpoints**

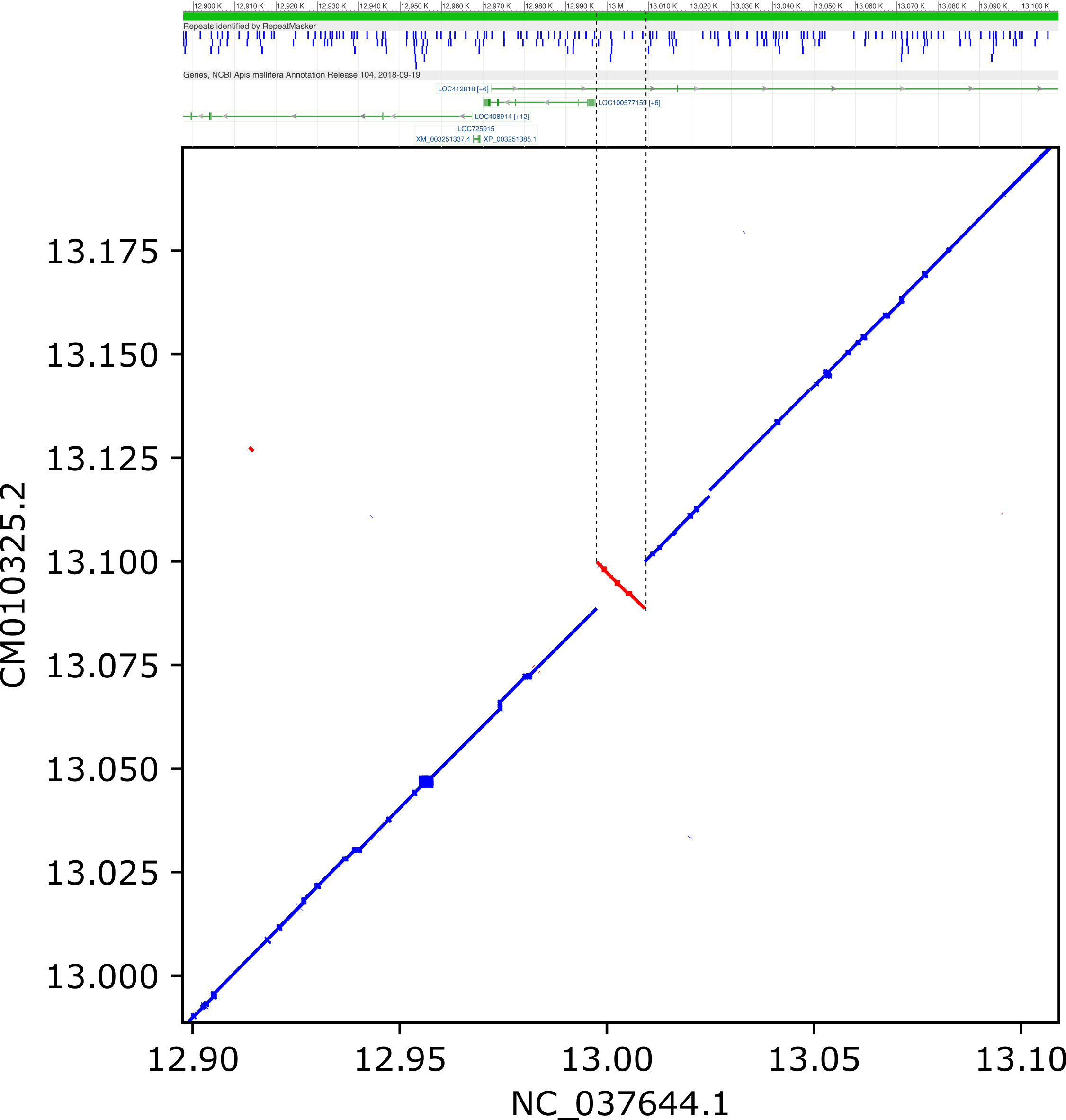

**Inversions > 1 kb between HAv3.1 and AMelMel1.1 on chromosome 7, detected by LAST and distant from contig breakpoints**

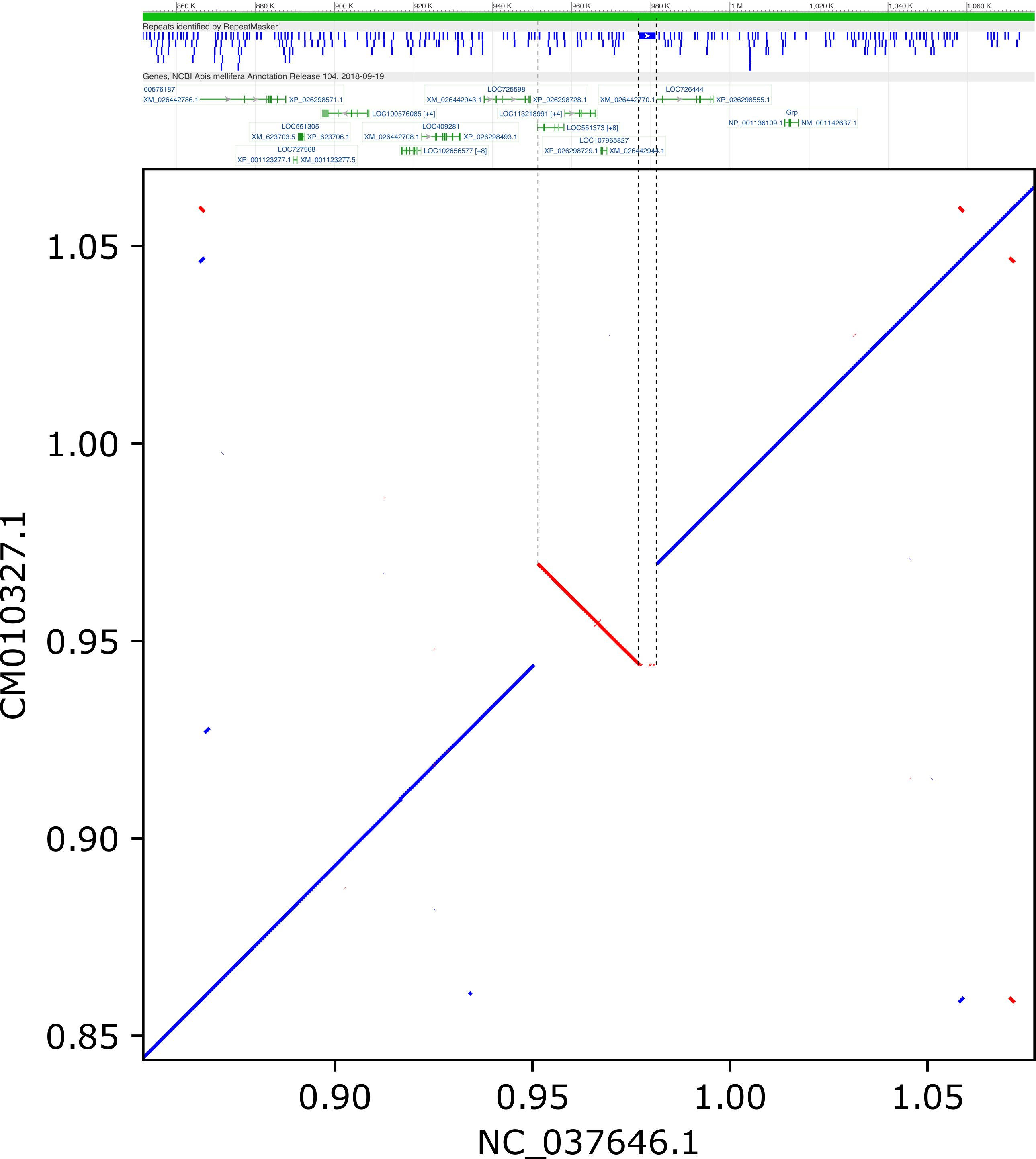

**Inversions > 1 kb between HAv3.1 and AMelMel1.1 on chromosome 9, detected by LAST and distant from contig breakpoints**

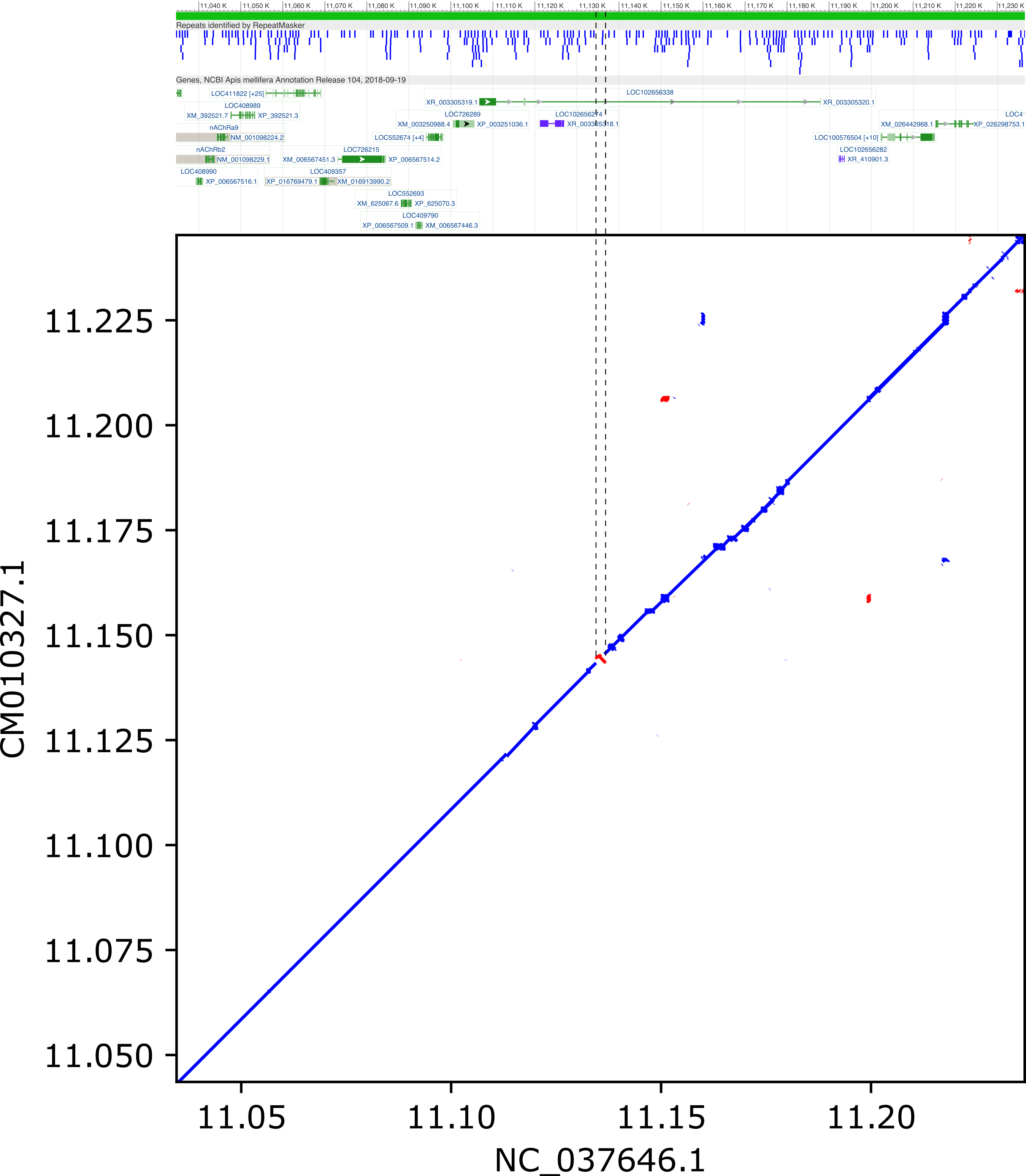

**Inversions > 1 kb between HAv3.1 and AMelMel1.1 on chromosome 9, detected by LAST and distant from contig breakpoints**

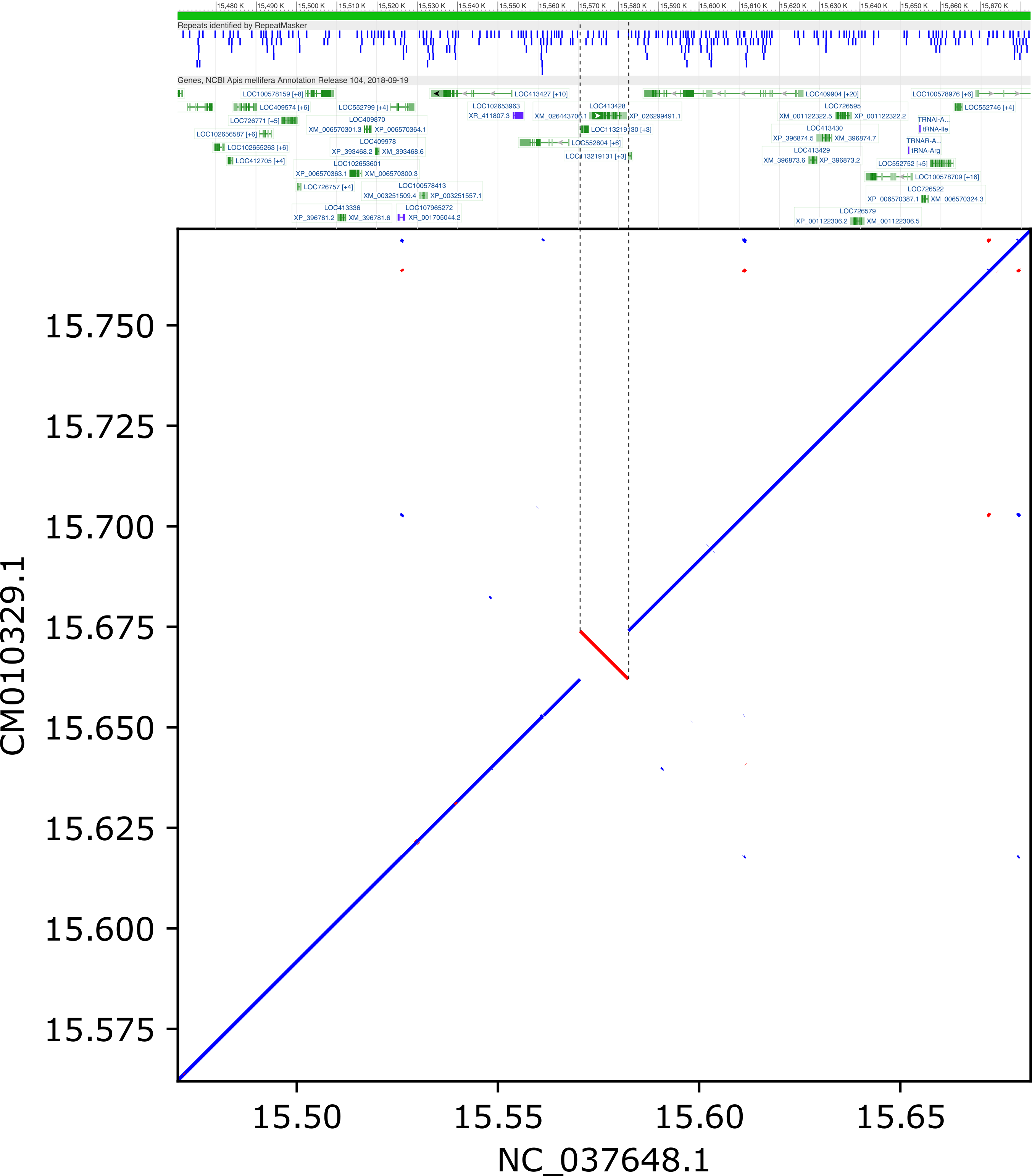

**Inversions > 1 kb between HAv3.1 and AMelMel1.1 on chromosome 11, detected by LAST and distant from contig breakpoints**

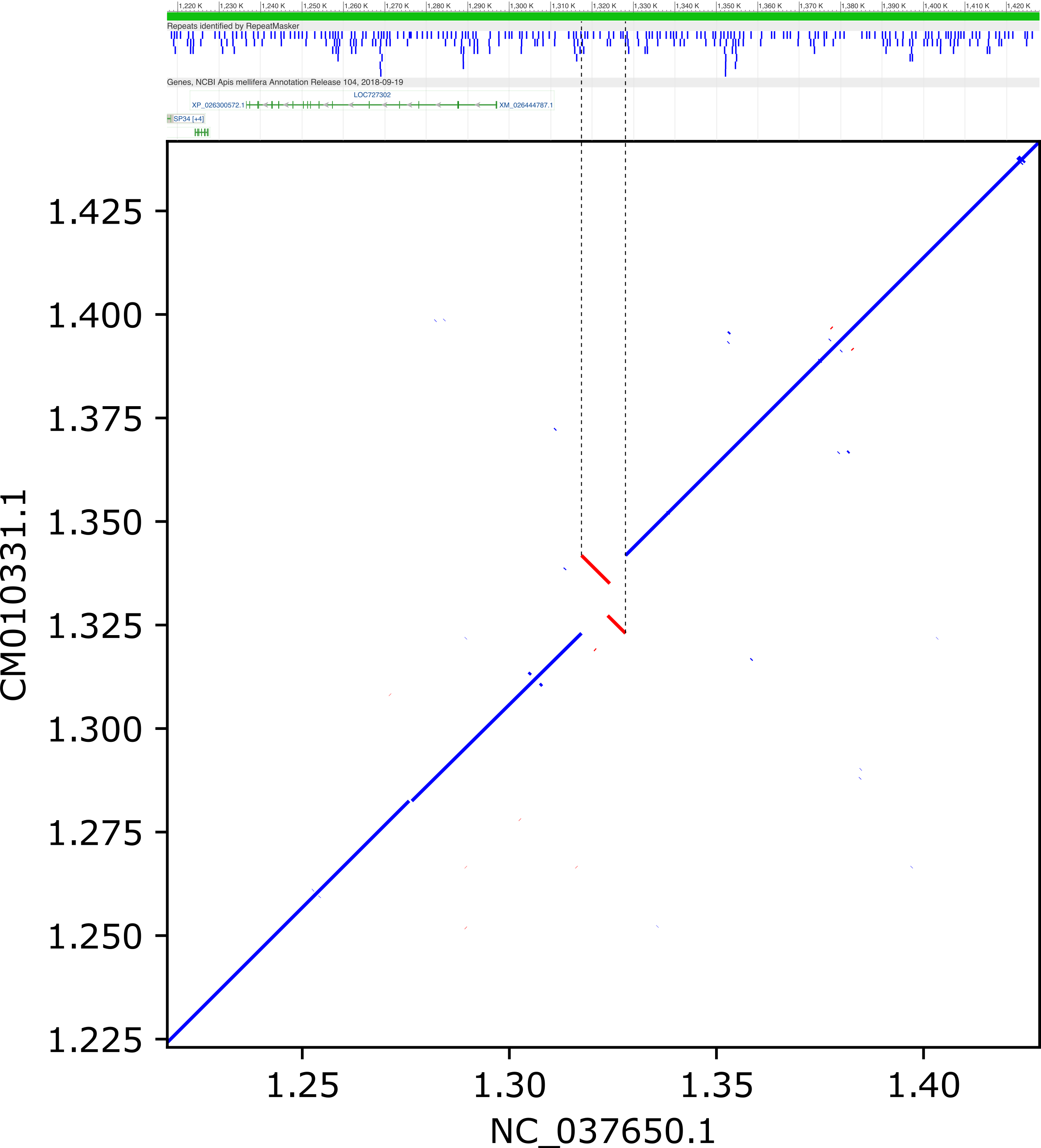

**Inversions > 1 kb between HAv3.1 and AMelMel1.1 on chromosome 13, detected by LAST and distant from contig breakpoints**

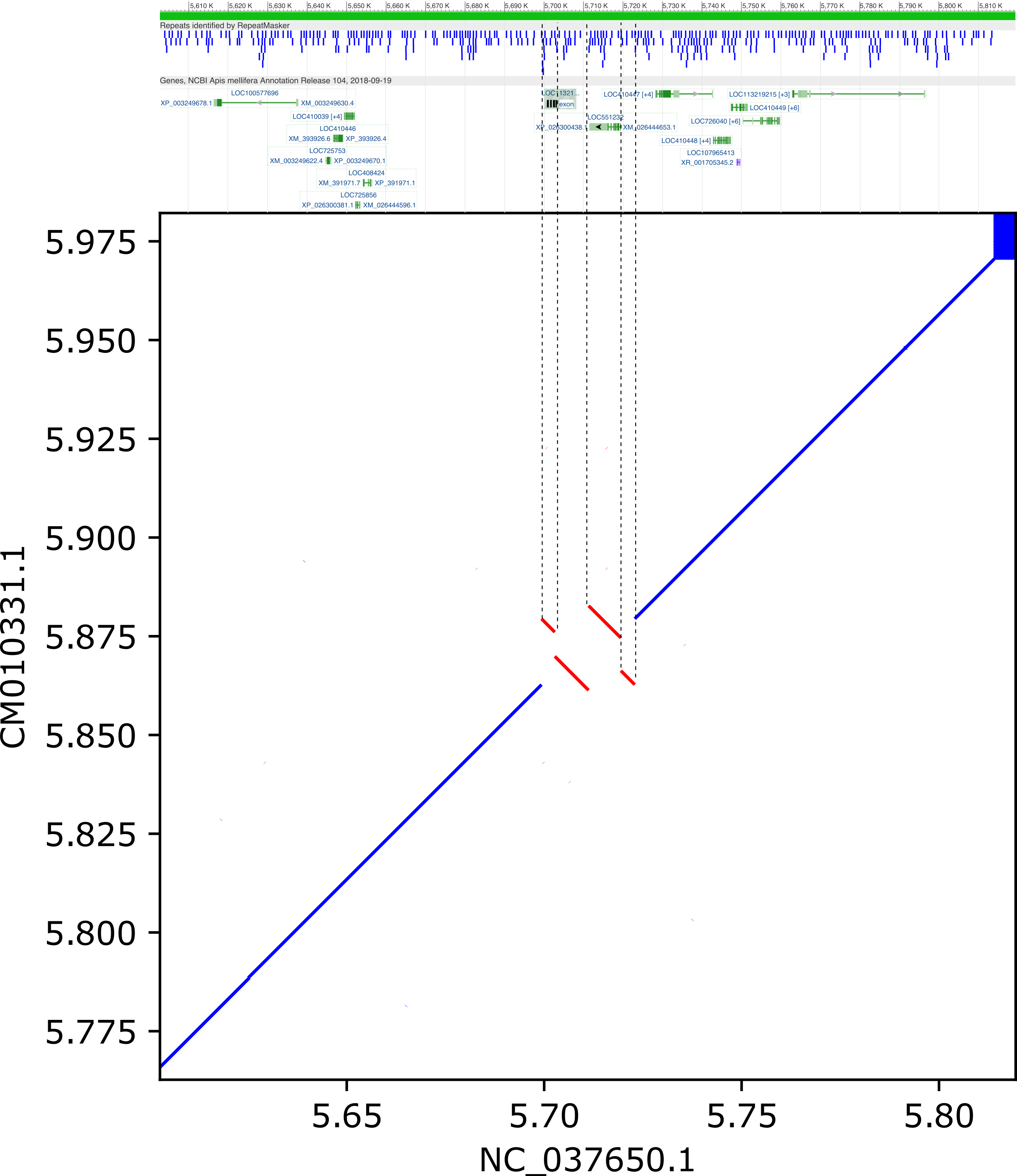

**Inversions > 1 kb between HAv3.1 and AMelMel1.1 on chromosome 13, detected by LAST and distant from contig breakpoints**

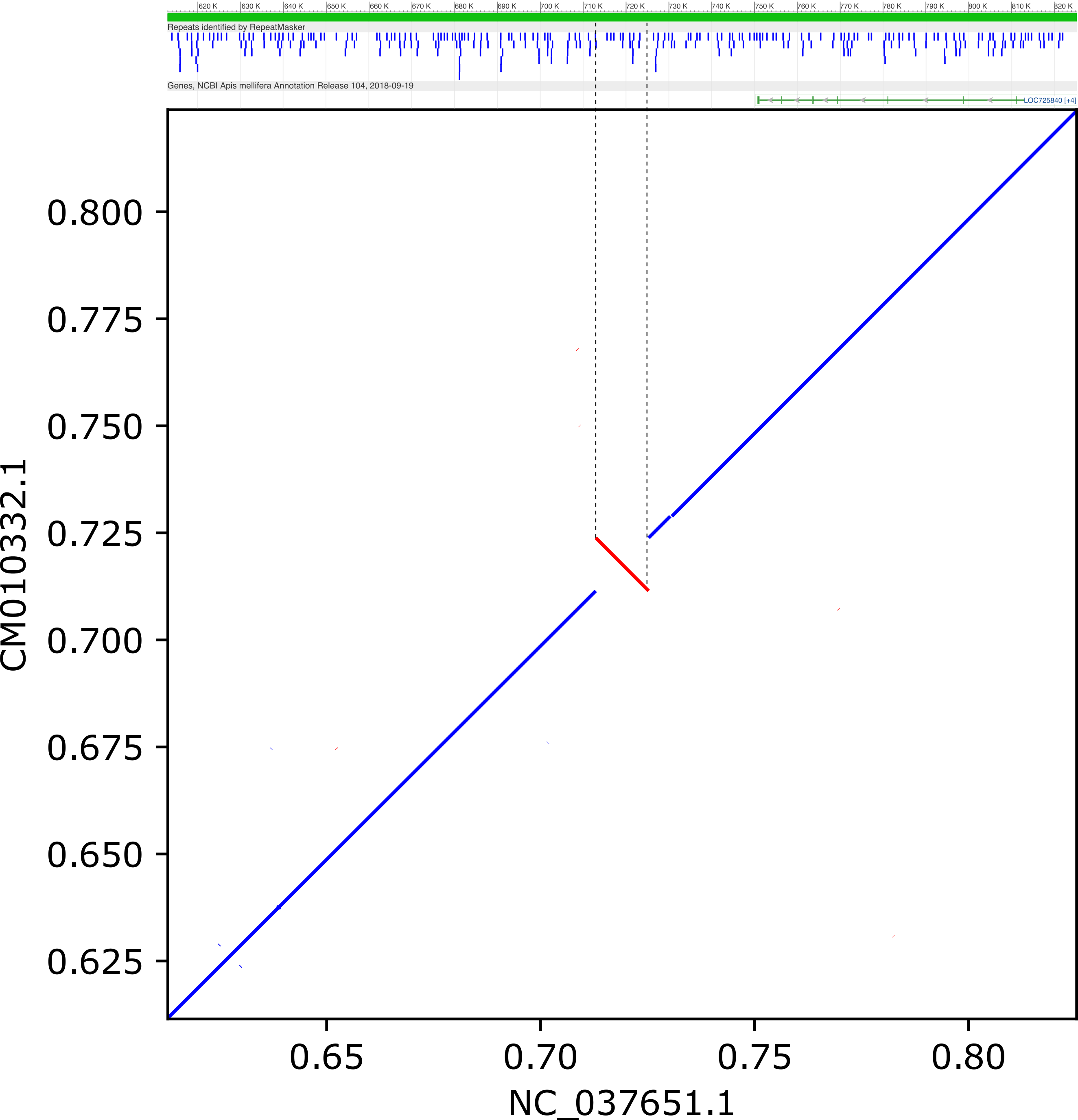

**Inversions > 1 kb between HAv3.1 and AMelMel1.1 on chromosome 14, detected by LAST and distant from contig breakpoints**

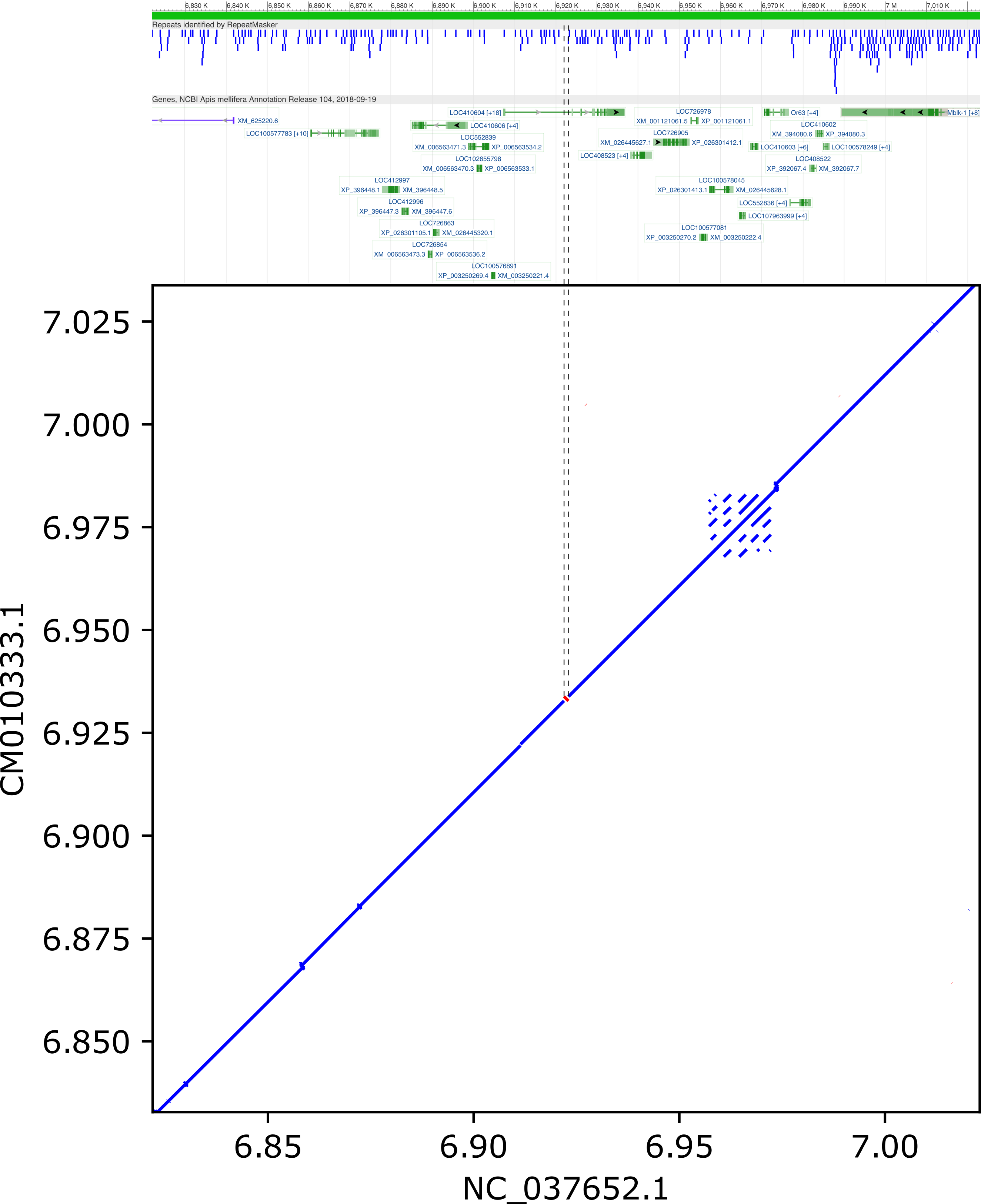

**Inversions > 1 kb between HAv3.1 and AMelMel1.1 on chromosome 15, detected by LAST and distant from contig breakpoints**
